# Trifunctional linkers enable improved visualization of actin by expansion microscopy

**DOI:** 10.1101/2023.05.10.540261

**Authors:** Gang Wen, Matthew Domenic Lycas, Yuqing Jia, Volker Leen, Markus Sauer, Johan Hofkens

## Abstract

Expansion Microscopy (ExM) revolutionized the field of super-resolution microscopy by allowing for subdiffraction resolution fluorescence imaging on standard fluorescence microscopes. However, it has been found that it is hard to visualize actin filaments efficiently using ExM. To improve actin imaging, multifunctional molecules have been designed, however, with moderate success. Here, we present optimized methods for phalloidin conjugate grafting that have high efficiency for both cellular and tissue samples. Our optimized strategy improves anchoring and signal retention by ∼10 times. We demonstrate the potential of trifunctional linkers (TRITON) for actin imaging in combination with immunolabeling using different ExM protocols. 10x ExM of actin labeled with TRITON enabled us to visualize the periodicity of actin rings in cultured hippocampal neurons and brain slices by Airyscan confocal microscopy. Thus, TRITON linkers provide an efficient grafting method, especially in cases where the concentration of target-bound monomers is insufficient for high-quality ExM.

---

Expansion microscopy (ExM) enables nanoscale imaging of structural details on conventional diffraction-limited microscopes through physically expanding samples in an isotropic fashion.^1–4^ Since its invention in 2015, ExM has been successfully applied to visualize various cellular components, such as proteins, RNA, and membranes in cultured cells or tissues,^3–8^ and organoids,^9,10^ as well as clinical specimens^11, 12^. Expansion factors of, *e.g.*, 10-20-fold^13–15^ or the combination of ExM with super-resolution microscopy methods^16–18^ such as stimulated emission depletion (STED) microscopy, structured illumination microscopy (SIM) and single-molecule localization microscopy (SMLM) have enabled multicolor fluorescence imaging of cells and tissue with spatial resolutions of ∼20 nm or better. In ExM, the performance of target visualization in the expanded state is influenced by both fluorescent label retention and the sufficient linkage of biological targets of interest into the polymer. Post-expansion immunostaining has been demonstrated as a good method for detecting endogenous proteins, as it not only avoids radical-induced fluorescent signal loss, but also increases epitope accessibility and reduces the linkage error caused by antibodies.^18–20^ In this strategy, endogenous proteins are covalently grafted into the polymer matrix with the succinimidyl ester of 6-((Acryloyl)amino)hexanoic acid (AcX), a commonly used anchoring reagent. Following homogenization with denaturation or proteolytic cleavage using proteinase K and expansion, preserved and newly exposed antigens can then be immunolabeled. As a result, protein organization in cells can be visualized with so far unmatched structural resolution,^18–20^ but even refined ExM strategies can currently not be applied for high-end visualization of actin filaments.

Actin is one of the most abundant proteins in eukaryotic cells and plays a vital role in cellular structure and functions, such as maintenance of cell morphology,^21^ cell motility,^22^ cell division,^23, 24^ transcriptional regulation,^25, 26^ endocytosis,^27^ intracellular trafficking,^28^ and muscle contraction^29^. Therefore, visualization of actin filaments using ExM has also garnered significant interest.^30–33^ However, due to the lack of an anchorable unit in commonly used phalloidin conjugates, retention of phalloidin labels remains challenging using the original ExM protocol.^30, 32^ To overcome this limitation, three phalloidin-actin complex anchoring strategies have been developed (Figure 1).^30–32^ The first strategy involves synthesizing a trivalent phalloidin linker (termed TRITON), which contains rhodamine B (Rh B), phalloidin and an acryloyl unit (monomer) for gel-grafting to enable visualization of actin filaments after expansion.^30, 33^ The second strategy, proposed by Park *et al*, uses anti-fluorophore antibodies to stain the fluorophore of phalloidin conjugates, and demonstrated ExM imaging of actin filament organization using the original ExM procedure.^32^ The third strategy, proposed by Trinks and coworkers, involves staining F-actin with a biotinylated phalloidin conjugate (phall-XX-biotin), followed by conjugation with fluorescently labeled streptavidin, in which streptavidin can provide lysine residues to introduce polymerizable moieties for gel-anchoring.^31^ However, in the last two methods, the size of antibodies (∼150 kDa) and streptavidin (∼60 kDa) compared to TRITON (<2 kDa) causes an increase in linkage error between actin and fluorescence of 17.5 nm and 5 nm, respectively, and these numbers are proportionally increased after expansion. Furthermore, due to the big size of antibodies and streptavidin, some actin-bound phalloidin conjugates cannot bind with antibodies and streptavidin, and thus are washed out after expansion as a result of insufficient grafting, reducing the achievable fluorescence intensity.^32^

**Figure 1.**
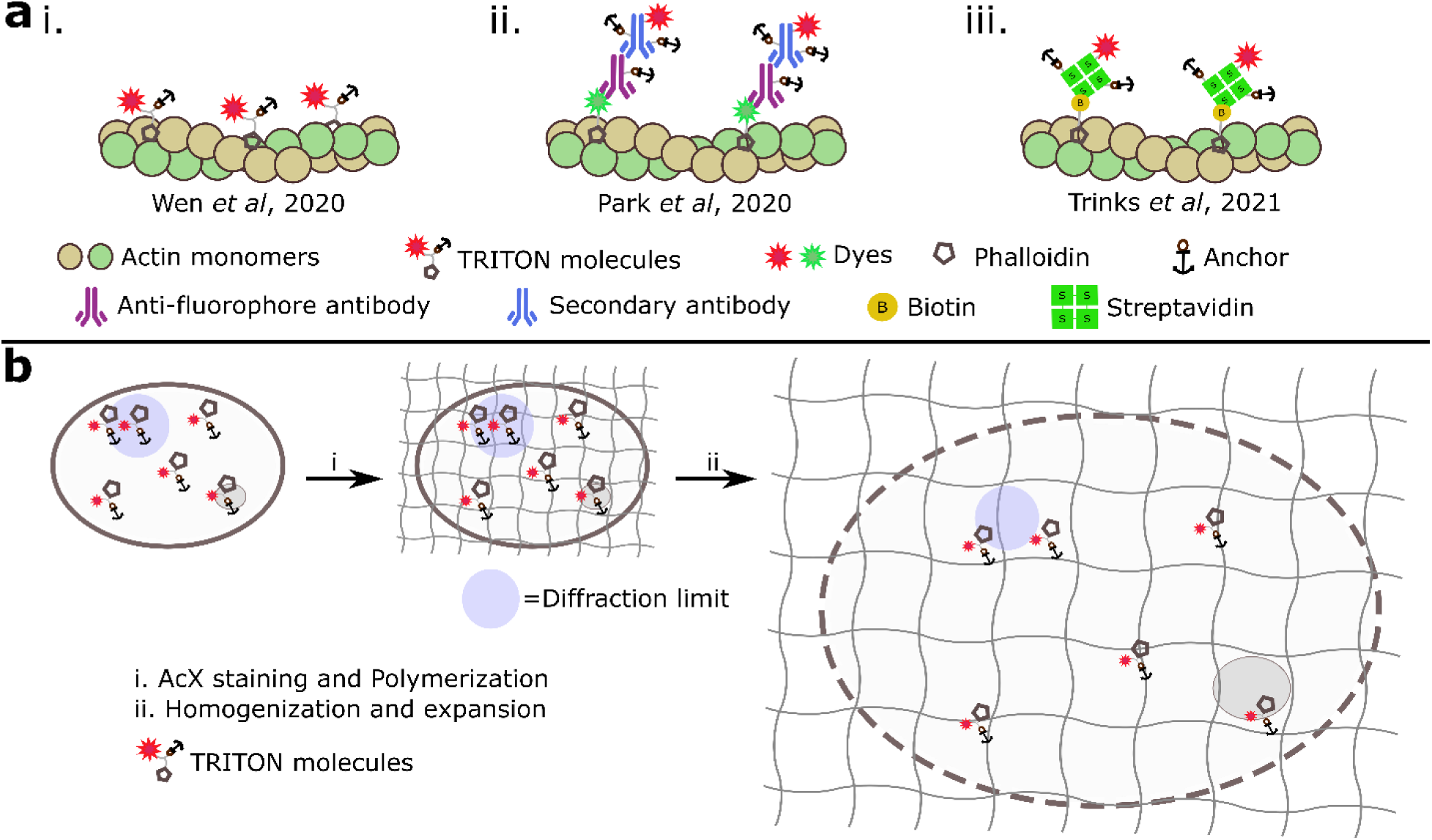
Schematic diagram of actin filament visualization in ExM. (a) Schematic of various grafting strategies of phalloidin conjugates *via* TRITON molecules (i), anti-fluorophore antibodies (ii) and fluorescently labeled streptavidin (iii). (b) Workflow of visualization of actin filaments using TRITON molecules in ExM. F-actin is stained by phalloidin TRITON molecules, linked into the gel, and expanded using standard ExM protocols.

Here, we examined various conjugation strategies to construct trivalent molecules and measured their effectiveness in actin staining experiments. We also optimized trivalent molecules with multiple anchors and corresponding staining protocols. We demonstrate that our TRITON linkers are compatible with various labeling strategies and multicolor staining. Finally, we evaluated their performance in cultured neurons and tissue samples. Using TRITON molecules and optimized staining protocols, we were able to maximize fluorescence signal retention, and thus visualize fine details of actin filament organization, such as periodicities of ∼185 nm by standard fluorescence microscopy even in brain slices.

## RESULTS AND DISCUSSION

### Evaluation of TRITON with different conjugation strategies

We first compared the post-digestion signal retention of a series of TRITON molecules, constructed *via* different conjugation strategies. To achieve this, we synthesized a trivalent linker, building on our previously reported TRITON molecules (compound **2**, Figure 2a).^30^ A triazine-based trivalent linker was also evaluated as part of this study (compound **3**, Figure 2a). In addition, triazine was chosen as a chemical scaffold to construct a maleimide-containing trivalent linker. A postulated benefit of this novel trivalent linker would be late-stage introduction of another fluorescent dye *via* a thiol-maleimide reaction (compound **4**, Figure 2a). We also synthesized a divalent linker by direct conjugation of phalloidin to Rh B using the same linker as compound **2**, as a control reference (compound **1**, Figure 2a). We then assessed their performance on signal retention post-digestion in cultured Hela cells. Therefore, fixed cells were stained with phalloidin TRITON molecules and their fluorescence intensity measured before and after expansion. After staining cells were gelated and digested following the original ExM protocol.^3^ After digestion, samples were washed three times by shrinking buffer (×5 min) to shrink the hydrogel and the fluorescence intensity of the same cell was measured again to quantify signal retention (signal retention = signal intensity_post-digestion_/signal intensity_pre-gelation_).

**Figure 2.**
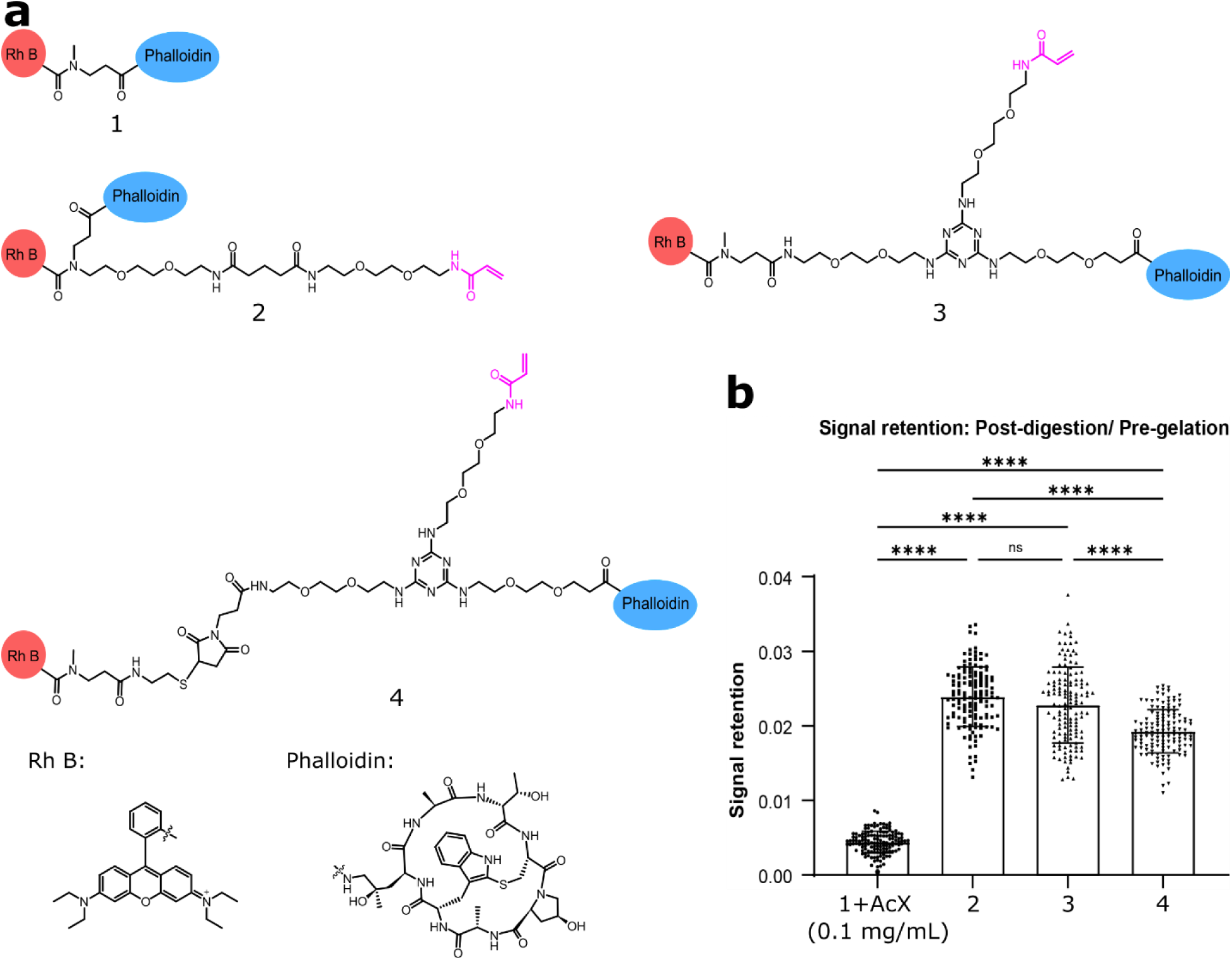
Quantification of signal retention of TRITON molecules. Comparison of different conjugation linkers clearly highlights the positive impact of multifunctional linkers for actin staining. (a) Molecular structures of different TRITON molecules. (b) Comparison of signal retention of different TRITON molecules. Bars represent the mean value and error bars represent the standard deviation. Statistical significance was assessed by one-way ANOVA test. **** p < 0.0001, ns (non-significant)= 0.0568. From left to right, mean values are 0.004 ± 0.001 (mean ± standard deviation), 0.024 ± 0.004, 0.023 ± 0.005, and 0.019 ± 0.003 (n= 137), respectively.

In line with literature observations,^32^ the fluorescence signal of compound **1** was almost completely lost post-digestion due to the lack of polymerizable moieties or primary or secondary amines required for linkage into the gel (Figure 2b). As a result, signal retention of compound **1** was only 0.004 ± 0.001 (mean ± standard deviation). Our TRITON molecules increased signal retention 6-fold (up to 0.024, compound **2**) and no significant difference of signal retention between two chemical skeletons was noticed (compound **2** and **3**), with 5-fold increased retention observed for compound **4** when compared to the staining with compound **1** and AcX.

### Optimization of TRITON grafting strategies

Due to the low signal retention of our previously developed phalloidin TRITON molecules and the volumetric dilution of signal observed with samples expansion,^30^ actin filament imaging required high laser intensities. While we had previously determined a survival rate of Rh B during radical-induced polymerization of around 55%,^30^ we attributed the observed low post-digestion signal retention to inefficient grafting of TRITON molecules into the hydrogel. Consequently, the majority of TRITON molecules washes out during the expansion process. Based on spatial binding models between phalloidin and F-actin,^34–37^ phalloidin conjugates bind actin filaments by burying deep into a cleft within the actin subunits. Hence, actual grafting of the acrylate to the propagating polymer mesh might be hindered. To improve the grafting efficiency, we explored two strategies to increase the actual acryloyl concentration bound to actin filaments. The first strategy was to introduce multiple acryloyl units to our previously reported TRITON molecule. Therefore, we used compound **2** (Figure 2a) and synthesized TRITON molecules with two and three acryloyl units (compound **5** and **6**, respectively, Figure 3a). In addition, based on the crystal structure of the actin-phalloidin complex (PDB ID: 6C1D),^34^ every actin subunit contains 19 lysine residues that can also be used to introduce the acryloyl moieties by applying Acryloyl-X (SE, Life Technologies, abbreviated AcX) to specimens prior to the gelation process. In principle, the latter strategy can not only facilitate the grafting of TRITON molecules, but also effectively reduce the loss of F-actin fragments post expansion, resulting from non-specific proteolytic digestion (proteinase K) and thus maximize the retention of actin filaments and accordingly the fluorescence signal measured for expanded actin filaments.

**Figure 3.**
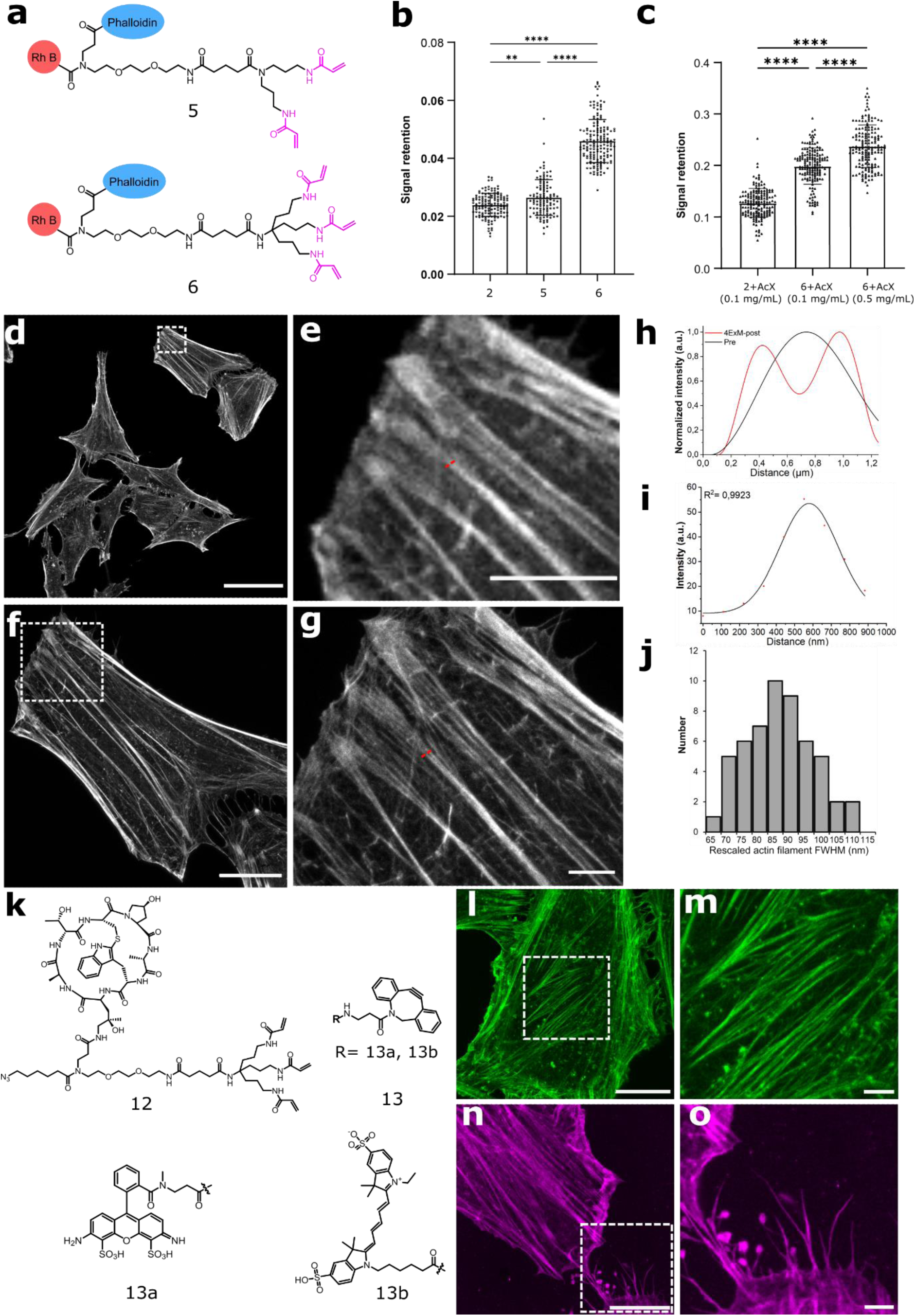
Optimization of TRITON-actin grafting efficiency in ExM. (a) Structures of optimized TRITON linkers equipped with multiple acryloyl units. (b) Comparison of post-digestion signal retention of actin filaments stained with TRITON linkers containing different acryloyl units. Bars represent the mean value and error bars represent the standard deviation. Statistical significance was assessed by one-way ANOVA test. ** p < 0.01; **** p < 0.0001. From left to right, mean values are 0.024 ± 0.004 (mean ± standard deviation, n= 137), 0.026 ± 0.006 (n= 112), 0.046 ± 0.007 (n= 162), respectively. (c) Comparison of post-digestion signal retention of actin filaments stained with TRITON linkers and AcX. Bars represent the mean value and error bars represent the standard deviation. Statistical significance was assessed by one-way ANOVA test. **** p < 0.0001. From left to right, mean values are 0.127 ± 0.028, 0.199 ± 0.035, 0.243 ± 0.049 (n= 162 from three independent samples), respectively. (d) Pre-expansion confocal fluorescence image of actin filaments stained with TRITON molecule **6** in 4x ExM. (e) Magnified views of the boxed regions in panel d. (f) Post-expansion confocal fluorescence image of actin filaments in HeLa cells. (g) Magnified views of the boxed regions in panel f. (h) Fluorescence intensity profiles along the red dotted lines in panel e (black line, pre-expansion) and g (red line, post-expansion). (i) A representative cross-sectional intensity profile of actin filaments with Gaussian fitting (solid line). (j) Distribution of Gaussian-fitted full width at half maximum (FWHM) of actin filament intensity profiles, yielding a resolution of 89 ± 11 nm (mean ± s.d., n= 53). (k) Chemical structures of compound **12** and **13** for post-gelation labeling. (l, m) Post-expansion confocal fluorescence image of actin filaments stained with compound **12** and **13a** (ATTO 488-DBCO) in 4x ExM. (n, o) Post-expansion confocal fluorescence image of actin filaments stained with compound **12** and **13b** (Sulfo-Cy5-DBCO) in 4x ExM. Scale bars, 50 μm (d, f, l, n), and 10 μm (e, g, m, o).

Next, we assessed the post-digestion signal retention of the improved TRITON molecules in fixed cells following the abovementioned protocol. Indeed, we observed a small increase in signal retention from 0.024 to 0.026 and 0.046 when increasing the acryloyl numbers of our TRITON molecules from one to three (compounds **2**, **5** and **6**, Figure 3b). In addition, signal retention improved substantially when we applied AcX to phalloidin TRITON-stained cells prior to gelation (Figure 3c). For example, by staining cells with AcX (0.1 mg/mL in PBS) prior to gelation, signal retention increased from 0.024 to 0.127 (5-fold) and 0.046 to 0.199 (4-fold) for compound **2** and **6**, respectively (Figure 3c). We rationalized that AcX signal retention improvement is due to the increased actin surface-bound acryloyl concentration enabling more efficient grafting of both, actin filaments and TRITON molecules.^38, 39^ Furthermore, we noticed a further improvement of signal retention of compound **6** from 0.199 to 0.243 when increasing the AcX concentration from 0.1 mg/mL to 0.5 mg/mL (Figure 3c). This finding is also consistent with the very recent paper of Ria and coworkers.^40^ Inspired by these results, we explored the impact of crosslinking agents on signal retention of actin filaments in more detail. Unfortunately, we did not observe an improved signal retention when modifying the general crosslinker, *e.g.*, by replacing the acrylamide moiety of AcX with methacrylamide (compound **7** in Figure S1) or introducing multiple acryloyl units to AcX (compound **8** in Figure S1). In addition, we also constructed phalloidin TRITON linkers with linear three or four acryloyl units (Figure S2). Compared to compound **6** containing branched three acryloyl units, compound **9** showed a slightly higher signal retention of 0.216 and no significant difference was observed between compound **9** and compound **10**, indicating that L-lysine can also tolerate enzymatic digestion with proteinase K (Figure S2). Interestingly, signal retention was decreased to 0.167 when four linear acryloyl units were introduced to phalloidin TRITON linkers (compound **11** in Figure S2). Finally, building on these insights, the scaffold of compound **6** was selected to construct optimized phalloidin TRITON linkers and its performance evaluated in biological systems.

### ExM of actin filaments with TRITON

Fluorescence labeling of Hela cells with compound **6** allowed us to visualize details of actin filament organization at an improved spatial resolution (Figure 3d-h). Comparing the same actin filaments pre- and post-expansion, we determined an expansion factor of 3.29 ± 0.14 (n= 34) (Figure S3), corresponding to a resolution of 89 ± 11 nm (mean ± standard deviation, n= 53) by rescaling the average Gaussian-fitted full width at half maximum (FWHM) with the expansion factor (Figure 3i,j and Figure S4).

We then broadened the application of TRITON molecules. Radical-induced polymerization is known to destroy organic fluorophores at a varying degree. For example, cyanine dyes (*e.g.*, Cy5) suffer from nearly complete destruction and thus cannot be used in pre-gelation labeling ExM.^2, 30^ Therefore, we constructed the azide-modified TRITON molecule **12** that enables post-gelation actin fluorescence labeling. Here fluorophores can be introduced after digestion *via* a strain-promoted click reaction, thus avoiding dye destruction during polymerization (Figure 3k-o). Advantageously, such phalloidin-actin staining can also be easily combined with immunolabeling for multicolor staining. To demonstrate this, we simultaneously visualized different biological targets in Hela cells using 4x ExM. Following a first immunofluorescence step for other proteins, we stained fixed cells with phalloidin TRITON. Next, cells were incubated with AcX and expanded. Afterwards, different cytoskeletal structures (actin filaments and microtubules) were clearly visualized at nanoscale (Figure S5a-d). Similar results were observed for simultaneous imaging of actin filaments and mitochondria (Figure S5e-h).

To further increase the spatial resolution, we combined ExM with structured illumination microscopy (SIM). As expected, we obtained an improved resolution showing more fine structures of actin fibers (Figure 4a-d). In addition, we also evaluated the performance of TRITON molecules in the recently developed Ten-fold Robust Expansion Microscopy (TREx)^15^ and achieved an average experimental expansion factor of 7.34 ± 0.23 (n= 22). Here, again a clear and continuous fluorescence signal of actin filaments was observed after expansion. Considering the experimental expansion factor, we estimated a spatial resolution of 39 ± 7 nm (n= 152) achieved for action imaging on a confocal fluorescence microscope (Figure 4e-j and Figure S6).

**Figure 4.**
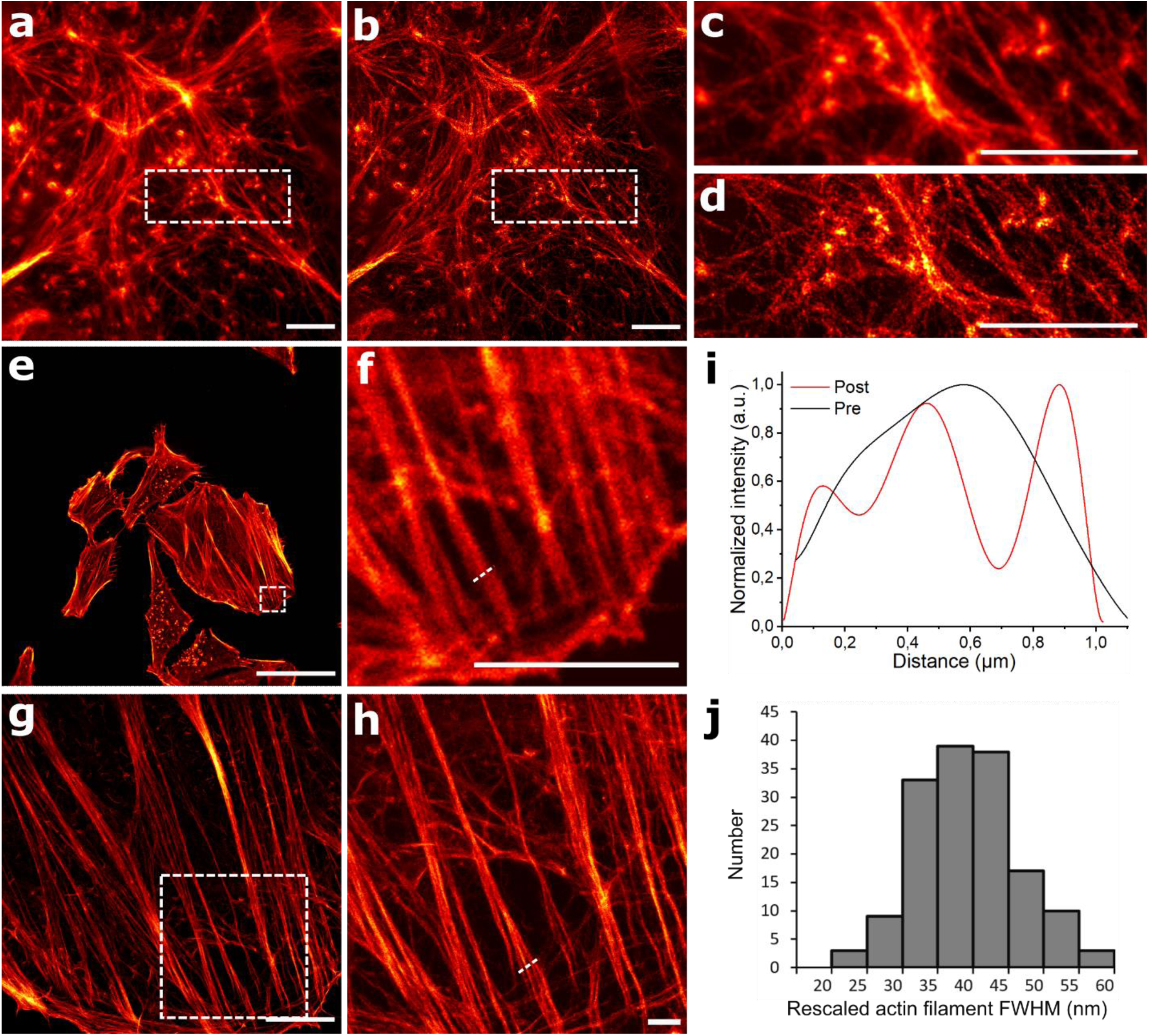
Actin filaments visualized by 4x ExM-SIM and 10x ExM. (a, b) Comparison of widefield (a) and SIM images (b) of 4x expanded actin filaments in Hela cells stained with TRITON molecule **6**. (c, d) Magnified view of the boxed regions in (a, b). (e) Pre-expansion confocal fluorescence image of actin filaments stained with TRITON molecule **6**. (f) Magnified view of the boxed region in (e). (g) Post-expansion confocal fluorescence image of actin filaments of the same cell as shown in (e, f) using TREx with an experimentally determined expansion factor of 7.34 ± 0.23 (n= 22). (h) Magnified view of the boxed region in (g), showing homogeneously labeled actin filaments at high spatial resolution. (i) Fluorescence intensity profiles along the white dotted lines in (f, h) (black line, pre-expansion) and (red line, post-expansion), highlighting the superior resolution. (j) Distribution of Gaussian-fitted full width at half maximum (FWHM) of TREx expanded actin filament intensity profiles resulting in an average value of 39 ± 7 nm (mean ± s.d., n= 152). Scale bars, 50 μm (e, g), and 10 μm (a-d, f, h).

### ExM imaging of actin filaments in neurons

Actin forms periodic rings along neuronal extensions with distances of approximately 185 nm and these rings require super-resolution imaging to resolve.^41^ By labeling actin in hippocampal neurons with TRITON compound **12**, the periodic structure could be clearly discerned by Airyscan confocal microscopy using the TREx expansion protocol (Figure 5). Fluorescence labeling of compound **12** was performed post-gelation by copper-free click-labeling with CF568-BCN. Being able to resolve the periodic structure of actin rings enabled us to determine a “molecular” expansion factor of the structure under investigation. Intensity profiles for TRITON staining were obtained and averaged along linear sections of neuronal extensions, showing the periodicity of actin (Figure 5a,b). The periodicity was captured by measuring the power spectra as a function of distance and averaging them across all regions of interest (Figure 5c,d). Although multiple peaks were obtained, the peak closest to 0 was chosen. This peak was found at 0.928 μm, and provided that the unexpanded distance for these structures is 185 nm, we determined an expansion factor of 5x for actin filaments in hippocampal neurons using the TREx protocol. In addition, we labeled pre- and postsynaptic proteins Bassoon and Homer, respectively, by immunolabeling, demonstrating the versatility of the TRITON labeling concept for multicolor super-resolution imaging (Figure 5e-j). To conclude, our data clearly show that TRITON molecules in combination with immunolabeling is ideally suited to study 3D synapse organization at high spatial resolution.

**Figure 5.**
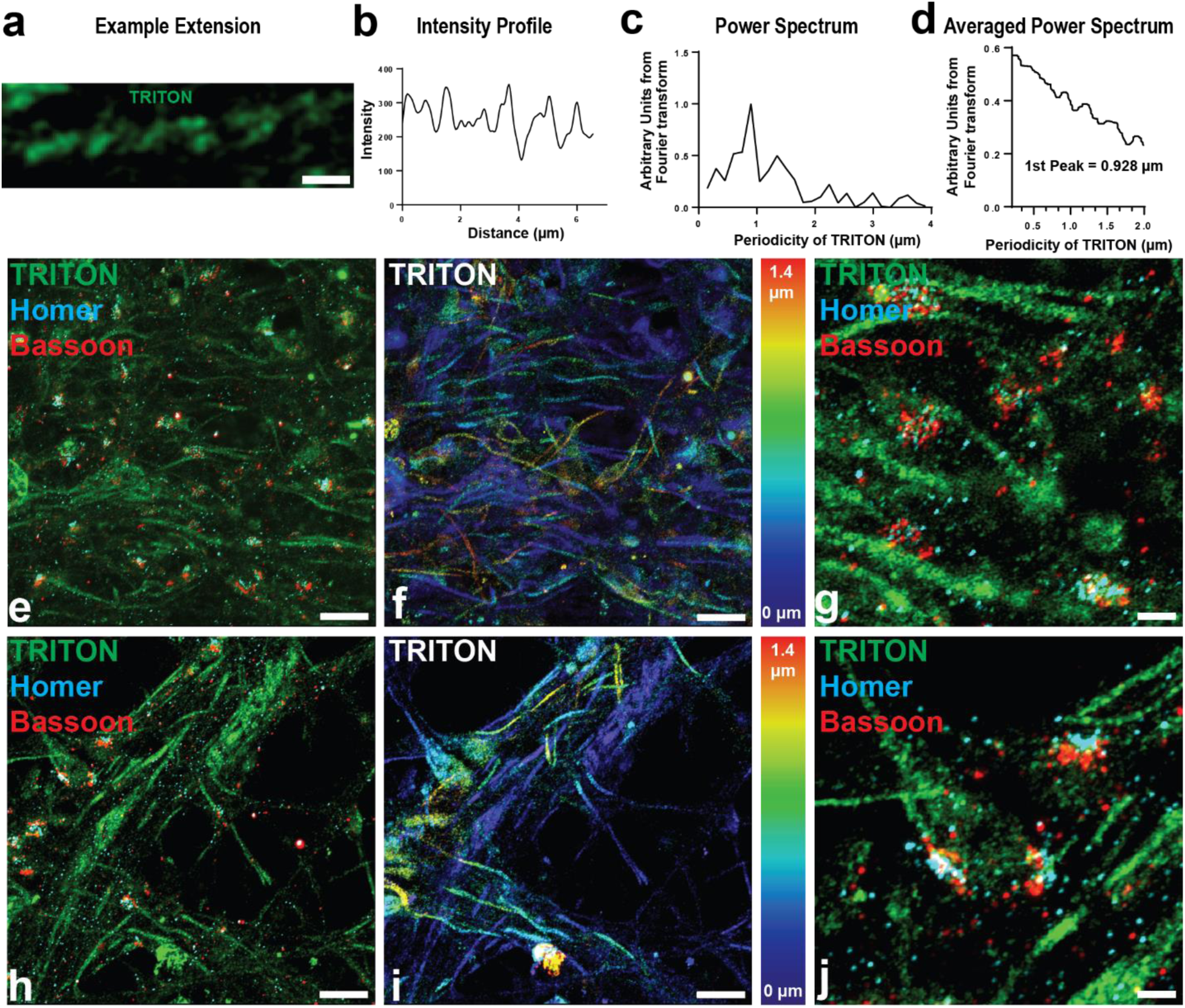
Super-resolution imaging of actin filaments and two synaptic proteins in neuronal cultures. (a-d) Determination of the expansion factor by identifying periodicity of actin signal in neuronal extensions. (a) Example neuronal extension from primary hippocampal cultures, expanded using the TREx protocol and labeled with compound **12** (post-gelation labeling of the azide group of **12** with CF568-BCN). (b) Intensity profile of the TRITON signal along the axis of the neuronal extension and averaged across a width of 15 pixels. (c) Power spectrum of the intensity profile shown in panel b, demonstrating that the TRITON signal shows a periodicity. (d) Averaged power signals across 35 regions of interest from 11 images to identify the periodicity of the actin signal in the neuronal extensions present in the images. This obtained value, 0.928 μm, was used to determine the actual expansion factor of the hippocampal neurons. (e-j) Airyscan confocal images of actin with Homer and Bassoon immunolabeling in expanded primary hippocampal cultures. (e, h) Representative three-color images of primary hippocampal cultures. (f, i) Corresponding 3D TRITON channel images. The z-position is indicated by the color coding. (g, j) Magnified images in e, h, showing actin, Homer, and Bassoon. Scale bars, 2 μm (e, f, h, i), 1 μm (a), and 0.5 μm (g).

### ExM imaging of actin filaments in mouse brain tissues

Finally, we assessed the performance of our TRITON linkers in labeling actin filaments in mouse brain tissues. Actin filaments were visualized using direct conjugation of phalloidin TRITON with organic dyes (compound **6**) and post-labeling of an azide group *via* a strain-promoted click reaction (compound **12**). Following the TREx expansion protocol, mouse brain slices were expanded 7.2x by comparing the same feature of tissue section pre- and post-expansion (Figure 6a,b). Here, both pre- and post-digestion labeling approaches allowed to visualize actin filaments with sufficient signal intensity after expansion (Figure 6c,d). Through combining TREx and Airyscan confocal microscopy, individual neuronal extensions were clearly distinguished in the striatum (Figure 6e-h). Many of these extensions showed the periodicity of actin labeling, as measured from the intensity profiles along different extensions (Figure 6i-l).

**Figure 6.**
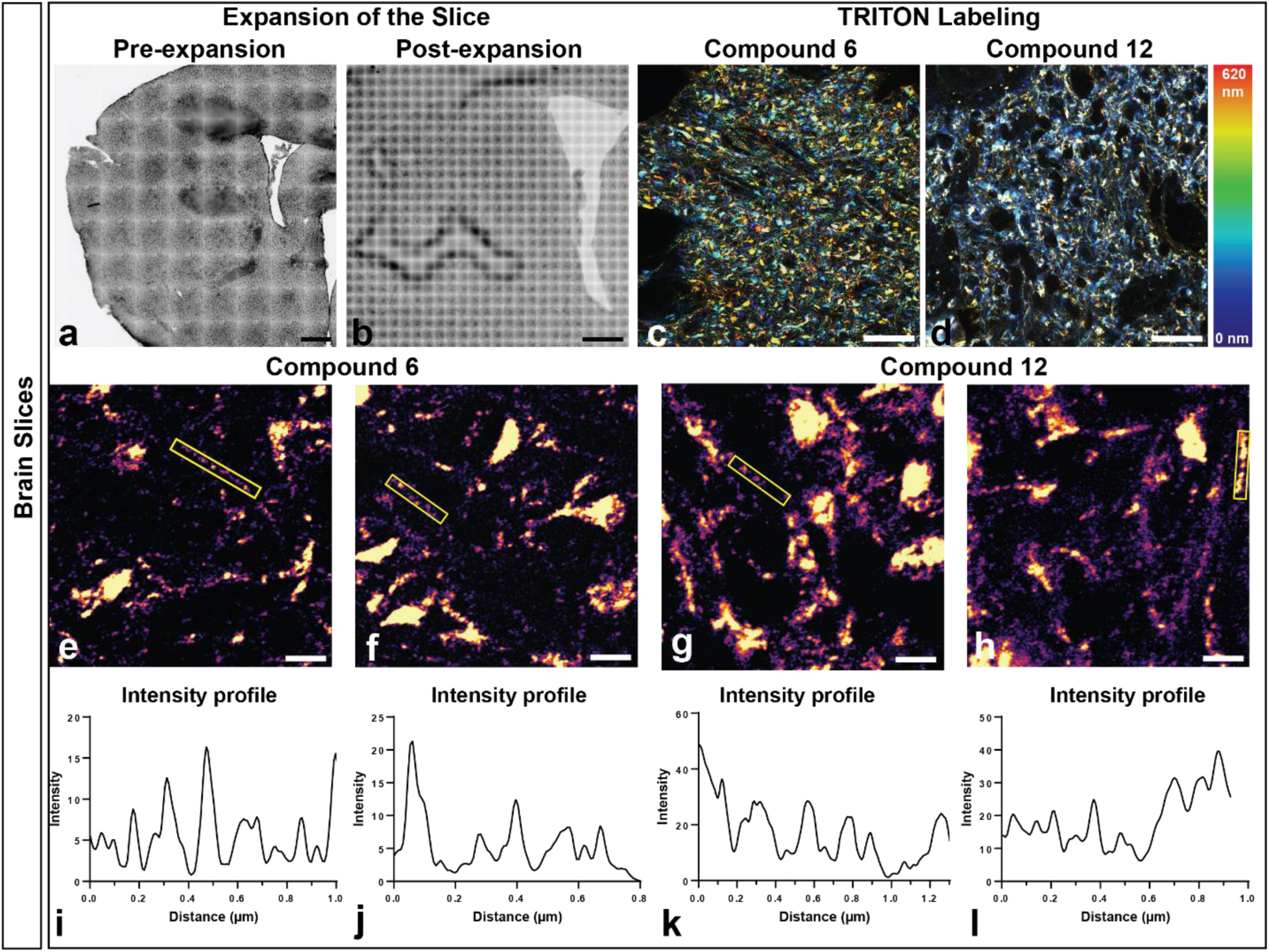
Super-resolution imaging of actin filaments in brain tissues with TRITON linkers. (a, b) Brightfield image of a mouse brain section before and after TREx expansion. (c, d) Airyscan confocal images of actin filaments in expanded neuronal extensions of the striatum labeled with compounds **6** and **12**. The z-position is color encoded. (e-h) Representative images of actin filaments in the striatum labeled with compounds **6** and **12**. (i-l) Intensity profiles taken along extensions found in e-h, demonstrating the periodicity of actin stained with TRITON linkers. Scale bars, 2000 μm (b), 500 μm (a), 5 μm (c, d), and 0.5 μm (e-h).

## CONCLUSIONS

In this work, we designed and synthesized a series of optimized phalloidin TRITON molecules with multiple anchors for improved actin polymer linkage and visualization of actin filaments in ExM. Our results demonstrate that the combination of phalloidin TRITON linkers with additional anchoring reagents (*e.g.*, AcX) improves the fluorescence signal retention, thus enabling the visualization of continuous actin filament organizations after 4x and 10x expansion. TRITON linkers with multiple anchors can be readily combined with other staining approaches, such as immunofluorescence staining, where we successfully performed multicolor imaging of actin filaments in combination with proteins, *e.g.*, microtubules and mitochondria. In combination with Airyscan confocal microscopy and 10x expansion, TRITON linkers allow fluorescence imaging of the actin cytoskeleton with superior spatial resolution using either directly fluorophore labeled TRITON linkers or post-gelation fluorescence labeling by strain-promoted click chemistry. Finally, we demonstrate the potential of TRITON linkers by resolving the periodicity of actin rings in cultured neurons and brain tissues. We expect that our phalloidin TRITON is a powerful tool to simplify and strengthen the visualization of actin filaments in cultured cells and various tissues.

## Methods

### Cell culture

HeLa cells (ATCC) were cultured in Dulbecco’s Modified Eagle Medium (DMEM; Life Technologies, high glucose (4.5 g/L), no glutamine, no phenol red) supplemented with 10% (*v*/*v*) fetal bovine serum, 50 μg/mL gentamicin (Life technologies) and 1% glutamax at 37 °C in a humidified 5% CO_2_ incubator. When cells reached to a confluency of 70-80%, they were washed three times with 1x DPBS (no calcium, no magnesium; Life Technologies), followed by detaching with 10x TrypLETM Enzyme (Life Technologies). Cells were seeded on 22 mm x 22 mm #1.5 coverslips at a density of 4 x 10^4^ cells/cm^2^ and incubated for 24 h prior to fixation.

### Fixation and staining of Hela cells

For actin imaging, cells were fixed 10 min with 4% paraformaldehyde (PFA) in 1x PBS and washed three times for 5 minutes with 1x PBS. The cells were then quenched with 100 mM NH_4_Cl in 1x PBS for 20 min, followed by washing three times for 5 minutes with 1x PBS. After permeabilization with 0.25% Triton X-100 in 1x PBS for 15 min, they were washed three times for 5 minutes with 1x PBS. The cells were then stained with 0.25 μM our phalloidin TRITON molecule in 1x PBS at room temperature for 1 h. After incubation, samples were rinsed twice with PBS and stained with 1 μg/mL DAPI and then washed three times with PBS prior to imaging. For grafting optimization of TRITON molecules, the cells were incubated with AcX (0.1 mg/mL) in 1x PBS overnight and then washed twice with 1x PBS for 15 minutes each prior to the gelation step.

For simultaneous imaging of actin and other proteins, cells were washed three times with pre-warmed 1x PBS, followed by fixation with 3% paraformaldehyde (PFA) and 0.1% glutaraldehyde (GA) in 1x PBS for 10 min. The cells were then quenched for 7 min with 0.1% sodium borohydride in 1x PBS and washed three times with 1x PBS. After permeabilization with 0.25% Triton X-100 and 3% BSA in 1x PBS for 15 min, cells were incubated with a solution of primary antibodies (mouse anti α-tubulin, Abcam, ab7291 or rabbit anti-TOM20) in staining buffer (PBS, 3% BSA) at a concentration of 2 μg/mL for 1 hour at room temperature and washed with PBS three times for 5 minutes each. Samples were then incubated with TRITON (ATTO 647N, acrylates, and TFP)-labeled secondary antibodies (Goat Anti-Mouse IgG, Abcam ab or

Goat Anti-Rabbit IgG, Abcam ab) for 1 hour in staining buffer with a dilution of 1:50. After incubation, samples were rinsed twice with PBS and washed twice with blocking buffer for 5 minutes each. After that, samples were stained with 0.25 μM phalloidin TRITON molecule (Rh 6G, acrylates, and phalloidin) in 1x PBS at room temperature for 1 h. After incubation, samples were rinsed twice with PBS and stained with 1 μg/mL DAPI, followed by washing three times with PBS. Next, stained cells were incubated with AcX (0.1 mg/mL) in 1x PBS overnight and then washed twice with 1x PBS for 15 minutes each prior to the gelation step.

### Gelation, digestion and expansion

For 4x ExM, gelation, digestion and expansion were performed as previously described.^30^ Briefly, stained samples were first washed with 80 μL monomer solution (2 M NaCl, 8.625% (w/w) sodium acrylate, 2.5% (w/w) acrylamide, 0.15% (w/w) N,N’-methylenebisacrylamide and 0.01% 4-hydroxy-TEMPO in PBS) supplemented with tetramethylenediamine (TEMED, 0.15% w/w) and ammonium persulfate (APS, 0.15% w/w) and the gelation was performed on the gelation chamber with another 80 μL gelation solution at 37 °C for 1.5 h. Next, gels were digested overnight with proteinase K (New England Biolabs) at a concentration of 8 units/mL in digestion buffer (50 mM Tris (pH 8.0), 1 mM EDTA, 0.5% TritonX-100, 0.8 M guanidine HCl) at room temperature. For 10x ExM, gelation and digestion were performed following the ten-fold robust ExM protocol. The gelation solution was prepared with 1.1 M sodium acrylate, 2.0 M acrylamide, 90 μg/mL N,N′-methylenebisacrylamide in 1x PBS, supplemented with 1.5 mg/mL APS, 1.5 mg/mL TEMED, and 15 μg/mL 4-hydroxy TEMPO and kept on ice before use. Next, stained samples were quickly washed with 80 μL freshly prepared gelation solution and the gelation was performed on the gelation chamber with another 80 μL gelation solution at 37 °C for 1.5 h. Following gelation, gels were digested overnight with proteinase K at a concentration of 8 units/mL in digestion buffer (50 mM Tris (pH 8.0), 2 mM CaCl_2_, 0.5% TritonX-100, and 0.8 M guanidine HCl) at 37 °C. After digestion, samples were re-stained with 1 μg/mL DAPI in 1x PBS and washed three times with 1x PBS. Finally, the digested gels were expanded with double-deionized H_2_O for 10 minutes. This step was repeated 3-4 times with fresh double-deionized until gels were fully expanded.

### Stabilization of expanded hydrogels

To stabilize expanded hydrogels during image acquisition, they were mounted on a poly-D-lysine-coated 6 well plate (Cellvis, Product #: P06-1.5H-N) following the previous procedure.^6^ The glass bottom of 6 well plate was treated with 1 mL 0.1% (w/v) poly-D-lysine at room temperature for 20 minutes. The glass bottom was then rinsed three times with 1 mL double-deionized H_2_O and was air-dried at room temperature before use. Next, the expanded gel was transferred to the coated plate and a coverslip was placed on the top of the gel for 5 min. Finally, the top coverslip was removed prior to imaging.

### Evaluation of post-digestion signal retention of various TRITON molecules

The phalloidin-actin staining protocol was mentioned above. After staining, samples were imaged as pre-gelation image, followed by gelation or incubating with acryloyl X (AcX, 0.1 mg/mL in PBS or 0.5 mg/mL in PBS) prior to the gelation step. Next, gels were digested with proteinase K at room temperature for 12 h. After rinsing samples with 1x PBS, they were re-stained with 1 μg/mL DAPI in 1x PBS and washed with 1x PBS. Samples were then washed three times with shrinking buffer (1 M NaCl and 60 mM MgCl_2_ in double-deionized water) and post-digestion imaging was performed using the same parameters as pre-gelation image to determine the fluorescent signal retention. We calculated the signal retention by dividing the total fluorescence intensity post-digestion by total fluorescence intensity obtained before gelation.

### Primary neuron culture

Primary hippocampal cultures were prepared from P0-P1 C57BL/6J mice. Hippocampi were dissected from the pups and placed in an eppendorf tube with 250 μL of reconstituted Neuronal Isolation Enzyme with papain (Thermo 88285) for 30 minutes at 37 °C. The hippocampal isolates were next titrated with flame treated Pasteur pipettes yielding single cell suspensions. The neurons were then applied to 24 mm poly-D-lysine treated coverslips (0.1 mg/mL poly-D-lysine in PBS for 1 hour at 37 °C, washed 5x in H_2_O) at a density of 100,000 cells per well. Cells were maintained for 16 days in Neurobasal-A medium (Thermo Fisher) supplemented with B-27 (Thermo) 2%, Glutamax 1% (Thermo Fisher), and gentamycin 5 mg/mL (Sigma) at 37 °C in 7.4% CO_2_.

### Primary neuron sample preparation

Cultured primary neuron were fixed at day 16 in paraformaldehyde (4%) for 15 minutes, followed by washing twice in PBS. Permeabilization was next performed with Triton-X-100 at 0.5% for 5 minutes in PBS. Samples were blocked for 20 minutes in blocking buffer consisting of 3% BSA in PBS. Primary antibody was next applied for 1 hour in blocking buffer (Bassoon SYSY 141 111 [1:200], Homer SYSY 160 003 [1:200]. The samples were washed three times for 5 minutes with blocking buffer. Secondary antibody and TRITON compounds were then applied. Secondary antibodies were applied at a concentration of 5 μg/mL, and compound **12** was applied at 1 μM. Samples were washed twice for 5 minutes with blocking buffer. Following this, samples were washed twice with PBS. To optimize the grafting of samples, they were then incubated in 0.1 mg/mL AcX in PBS with 200 mM NaHCO_3_ overnight at room temperature. Stained neuron samples were first washed in PBS and then gelated in monomer solution consisting of 1.1 M sodium acrylate, 2.0 M acrylamide, 90 μg/mL N,N′-methylenebisacrylamide in PBS following the TREx protocol.^15^ Prior to gelation, 1.5 mg/mL TEMED and 1.5 mg/mL APS were added to the monomer solution. The sample was placed cell side down over monomer solution on parafilm, and this situation was maintained over ice for 5 minutes, followed by being placed in 37 °C for 1 hour. Following gelation, the sample was digested in Proteinase K solution (1:100 Proteinase K (Ambion AM2546), 50 mM TRIS pH 8, 800 mM guanidine HCl, 2 mM CaCl_2_, 0.5% Triton-X-100) at 50 °C overnight. Gels were immersed in PBS for three rounds over the course of an hour to remove digestion buffer. Next, gels were placed in 10 μM CF568-BCN (Biotium) for 2 hours. Gels were washed with PBS 4 times for total time of 1 hour to remove free dye. Samples were then expanded through multiple water washed until the gel expanded no further. Subsections of the gels were mounted onto poly-D-lysine-coated Labteks for imaging.

### Mouse brain slice preparation

Mouse brain slices were obtained from male C57BL/6 mice 12–16 weeks old. Mice were anesthetized isoflurane and perfused with paraformaldehyde (4%) in PBS. Brains were removed and placed in paraformaldehyde (4%) in PBS for 24 hours followed by sucrose (30%) in PBS for 24 hours at 4 °C. Brains were snap frozen on dry ice and stored at −80 °C. Brains were sliced on a cryostat at a thickness of 10 μm, and slices were transferred to 3-aminopropyltriethoxysilane-coated coverslips. Samples were then stored at −20 °C in 50:50 glycerol:PBS. Staining of the slices commenced with washing twice for 15 minutes with glycine (20 mM) and NH_4_Cl (50 mM) in PBS. Slices were then permeabilized for 30 minutes with 0.3% Triton-X-100 in 5% BSA. Next, samples were labeled with compound **6** (0.25 μM) or compound **12** (1 μM) in blocking buffer (PBS with 5% BSA and 0.3% Triton-X-100) for 5 hours on a shaker. Samples were washed in blocking buffer in increments of 10, 10, 10, 90 minutes, following by washing three times for 10 minutes with PBS. Hoechst 34580 was applied (10 μg/mL) for 30 minutes. Samples were then washed three times with PBS to remove free dye. To optimize the grafting of samples, they were placed in 0.1 mg/mL AcX in PBS with 200 mM NaHCO_3_ overnight at room temperature. Gelation, digestion, fluorophore labeling, and expansion were performed identically to the neuron culture gels except that ATTO 488-DBCO was used to introduce fluorescence for post-labeling.

### Microscopy and image analysis

Confocal imaging and original data were acquired on an inverted Leica true confocal scanner SP8 X system (Wetzlar, Germany) using a HCPLAPO CS2 63× 1.2 NA water-immersion objective for all images. The nuclei stained with DAPI were activated by a 405 nm pulsed diode laser. A supercontinuum white light laser (470 nm-670 nm, SuperK EXTREME/FIANIUM, NKT photonics, Birkerød, Denmark) was used at 561 nm for excitation of rhodamine B. Airyscan confocal data was captured on a Zeiss LSM 980 with Aryscan 2. Neuron culture data was capture with a frame size of 76.99 μm^2^ across 1810 pixels^2^. Images were taken with sampling equal to 2. The 488, 561, and 647 laser lines were used, scanning by frame with a pixel time of 2.3E-06 seconds. Images were taken with a z step size of 150 nm. Brain slice data was taken with a frame size of 201.64 μm^2^ across 4072 pixels^2^. The 488 laser line was used, scanned by frame with a pixel time of 2.1E-06 seconds. Data was taken with a z step size of 180 nm. The objective used for all experiments was the plan-apochromat 63×/1.40 Oil DIC M27. Airyscan processing was done with ZEN (blue edition) version 3.5.093.00000. SIM data was captured on a Zeiss Elyra 7 with a Plan-Apochromat 63×/1.4 oil DIC M27 objective and the 1.6x tub lens. Grating period was 819.4 nm, 3D SIM processing was used with settings of 20 iterations, medium filter, 0.025 regularization weight. Imaging was done with excitation lasers of 488 nm, 561 nm, 642 nm light imaged individually. Laser emissions were filtered by BP 495-550, BP 570-620, and LP 655 filters, respectively. Images were analyzed with ImageJ after acquisition. Expansion factor was obtained by dividing the expanded distance by the original distance of the actin filaments in Hela cells. The experimental image resolution was determined with ImageJ FWHM_line plugin with the Gaussian fitting and recalculated by expansion factor.

## ASSOCIATED CONTENT

### Supporting Information

The Supporting Information is available free of charge.

Detailed synthesis process and characterization of compounds, raw data, expansion factor calculation and resolution quantification.

## AUTHOR INFORMATION

### Author Contributions

G.W., V.L., M.S., and J.H. designed the research. G.W. performed synthesis and cellular experiments. M.L. performed neuron and tissue staining. G.W., Y.J. and M.L. analyzed the data together. G.W. V.L., and M.L. wrote the article with input from all other authors.

### Notes

The authors declare no competing financial interests.

## Supporting information

Supporting information includes chemcial characteristics and raw images

## ACKNOWLEDGMENT

This work was supported by funding from the China Scholarship Council to G.W. (Nr 201806210078), from the Flemish government through long-term structural funding Methusalem (CASAS2, Meth/15/04) to J.H., and from the European Research Council (ERC) under the European Union’s Horizon 2020 research and innovation programme (grant agreement No 835102) to M.S.. The authors thank C. Jackers, R. Nuyts and D. Linhares for technical support.

---

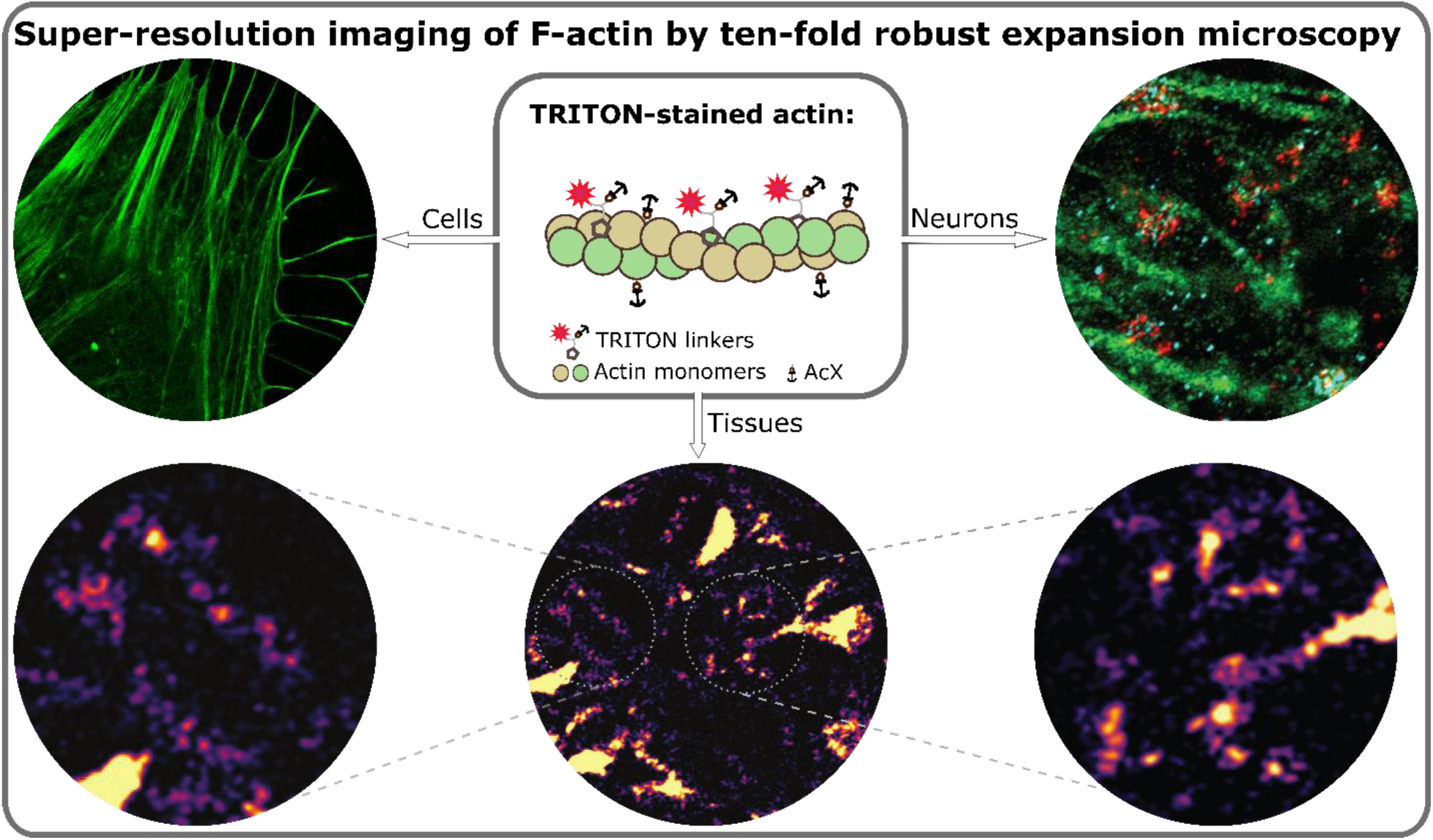
For Table of Contents only

