## Supporting information includes chemcial characteristics and raw images for "Trifunctional linkers enable improved visualization of actin by expansion microscopy"

#### 1. Synthesis and Characterization

##### 1.1 General

Commercial chemicals were obtained from Sigma-Aldrich, Fluorochem, TCI or ACROS and were used without further purification. Dry solvents were used as received from commercial sources. Thin layer chromatography (TLC) with silica gel plates (Kieselgel 60 F254 plates, Merck) was used to monitor reactions under UV light or dipping in a basic aqueous solution of potassium permanganate, followed by brief heating. 70 - 230 mesh silica 60 (E. M. Merck) was used on column chromatography. Mass spectra was achieved using a Shimadzu LC-MS 2020 Liquid Chromatograph Mass Spectrometer (ShimPack Gist C18 2 μm, 2.1x100 mm). Reverse-phase preparative HPLC was performed on a Shimadzu LC-MS 2020 Liquid Chromatography system with a Shim-pack GIST C18 2 μm column (Eluent A: 0.1% HCOOH in Milli-Q water, Eluent B: MeOH, gradient elution (20%-100% B from 0-30 min; 100% B from 30-35 min, pump flow: 2.00 mL/min). $^1$H NMR and $^{13}$C NMR spectra were recorded on a Bruker Avance 400 MHz or a Bruker Avance Ⅱ+ 600 MHz spectrometer using MeOD-*d*$_4$ or $CDCl_3$ as a solvent.

##### 1.2 Synthetic procedures

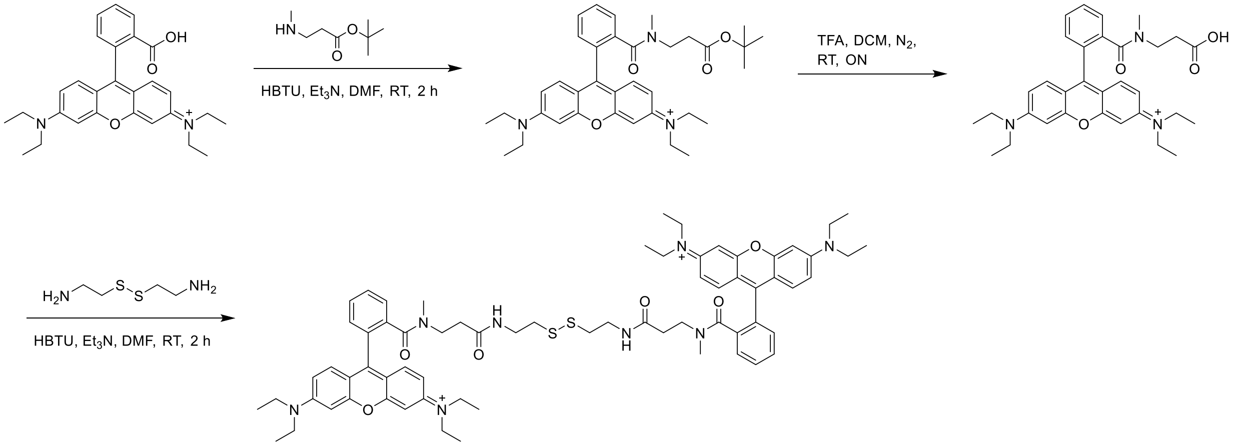

##### Synthesis of compound S1

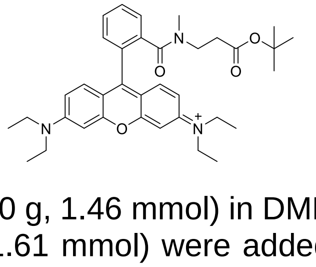

To a solution of rhodamine B (0.70 g, 1.46 mmol) in DMF (4 mL), triethylamine (0.61 mL, 4.38 mmol) and HBTU (0.61 g, 1.61 mmol) were added. After 10 minutes, *tert*-butyl 3-(methylamino)propanoate (0.28 g, 1.75 mmol synthesized from the literature[1]) was added and the reaction mixture was stirred at room temperature for 2 h. After complete reaction,

100 mL ethyl acetate was added into the reaction flask, followed by washing with water (100 mL, 2 x) and brine (100 mL). The organic layer was dried over $Na_2SO_4$ and evaporated under reduced pressure. The residue was purified by column chromatography to yield the intermediate as a red solid (85%). LC-MS (ESI$^+$): calculated for $C_{36}H_{46}N_3O_4$ $[M]^+$ m/z: 584.35; found: 584.10; $^1$H NMR (400 MHz, $CDCl_3$) δ 7.82 – 7.65 (m, 3H), 7.55 – 7.50 (m, 1H), 7.30 (d, *J* = 9.5 Hz, 2H), 7.09 (dd, *J* = 9.5, 2.4 Hz, 2H), 6.98 (d, *J* = 2.4 Hz, 2H), 3.71 (q, *J* = 7.2 Hz, 8H), 3.43 (t, *J* = 7.0 Hz, 2H), 2.92 (s, 2H), 2.83 (s, 1H), 2.60 – 1.95 (t, *J* = 7.0 Hz, 2H), 1.47 – 1.28 (m, 21H); $^{13}$C NMR (101 MHz, $CDCl_3$) δ 172.31, 170.90, 159.64, 157.50, 157.26, 137.72, 133.49, 132.17, 132.01, 131.62, 131.26, 129.05, 115.51, 115.08, 97.66, 82.19, 47.17, 45.10, 38.98, 33.88, 28.56, 13.15.

**Synthesis of compound S2**

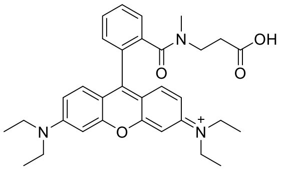

To a solution of the intermediate **S1** in DCM (1 mL), TFA (1 mL) was added and the reaction mixture was stirred overnight at room temperature. After complete reaction, all solvents were evaporated to yield the intermediate as a red solid, of sufficient purity for further use (95%).

**Synthesis of compound S3**

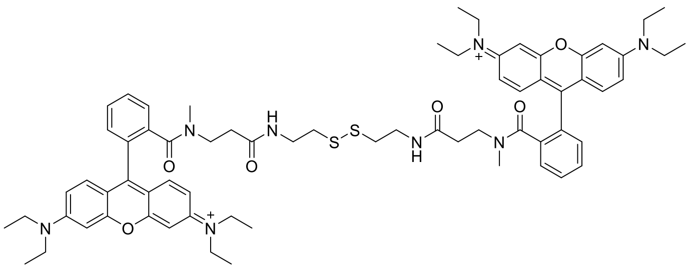

To a solution of the intermediate **S2** (0.78 g, 1.38 mmol) in DMF (5 mL), triethylamine (0.53 mL, 3.78 mmol) and HBTU (0.496 g, 1.33 mmol) were added. After 10 minutes, cystamine dihydrochloride (0.142 g, 0.629 mmol) was added and the reaction mixture was stirred at room temperature for 2 h. After complete reaction, 200 mL ethyl acetate was added into the reaction flask, followed by washing with water (200 mL, 2 x) and brine (200 mL). The organic layer was dried over $Na_2SO_4$ and evaporated under reduced pressure. The residue was purified by column chromatography to yield the intermediate as a red solid (61%). LC-MS (ESI$^+$): calculated for $C_{68}H_{84}N_8O_6S_2$ $[M]^{2+}$ m/z: 1172.59; found: 586.55; $^1$H NMR (400 MHz, $CDCl_3$) δ 7.67 – 7.54 (m, 6H), 7.25 (ddd, *J* = 27.2, 16.4, 8.1 Hz, 7H, including 1H from $CHCl_3$), 6.86 (dd, *J* = 9.6, 2.3 Hz, 4H), 6.75 (d, *J* = 14.5 Hz, 4H), 3.68 – 3.48 (m, 15H), 3.47 – 3.39 (m, 5H), 3.39 – 3.29 (m, 3H), 2.86 (s, 4H), 2.82 – 2.64 (m, 6H), 2.58 – 1.94 (m, 4H), 1.29 (t, *J* = 7.0 Hz, 24H); $^{13}$C NMR (101 MHz, MeOD-

*d*$_4$) δ 173.34, 170.86, 159.52, 157.39, 137.65, 133.49, 132.29, 131.97, 131.51, 131.26, 129.14, 128.55, 127.41, 118.96, 115.52, 115.07, 111.71, 97.64, 55.11, 47.18, 45.68, 39.77, 39.02, 38.47, 35.75, 34.42, 32.85, 13.18.

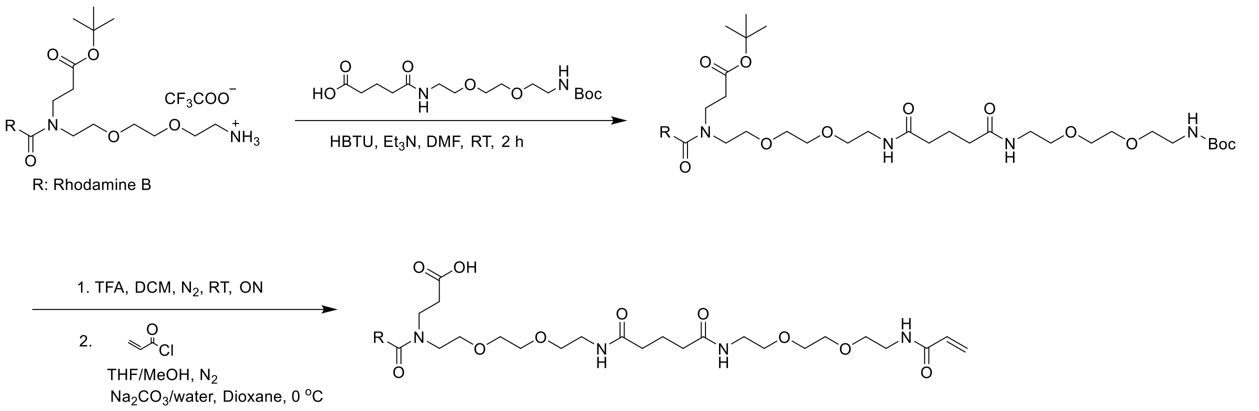

#### Synthesis of compound S4

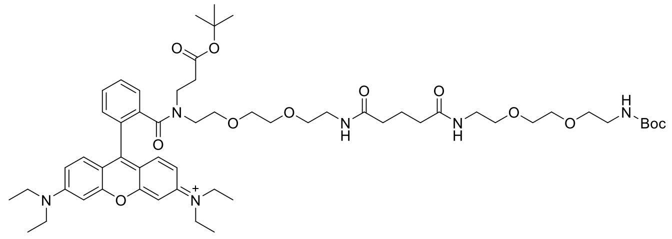

To a solution of the modified rhodamine B (110 mg, 135 µmol, synthesized from the literature[2]) in DMF (1.5 mL), trimethylamine (56 µL, 404 µmol), HBTU (56 mg, 148 µmol) and the modified PEG fragment (54 mg, 148 µmol, synthesized from the literature[3]) were added and the reaction mixture was stirred at room temperature for 2 h. After complete reaction, 40 mL ethyl acetate was added into the reaction flask, followed by washing with water (40 mL, 2 x) and brine (40 mL). The organic layer was dried over $MgSO_4$ and then evaporated under reduced pressure. The residue was purified by column chromatography to yield the intermediate as a red solid (72%). LC-MS ($ESI^+$): calculated for $C_{57}H_{85}N_6O_{12}$ $[M]^+$ m/z: 1045.62; found: 1045.45; $^1$H NMR (400 MHz, MeOD-*d*$_4$) δ 7.88 – 7.72 (m, 3H), 7.57 – 7.48 (m,1), 7.33 (dd, *J* = 9.5, 7.2 Hz, 2H), 7.15 – 7.06 (m, 2H), 7.00 (dd, *J* = 7.4, 2.3 Hz, 2H), 3.77 – 3.68 (m, 8H), 3.66 – 3.61 (m, 5H), 3.61 – 3.50 (m, 8H), 3.49 – 3.35 (m, 8H), 3.32 – 3.27 (m, 2H), 3.23 (t, *J* = 5.7 Hz, 2H), 3.03 (t, *J* = 5.2 Hz, 1H), 2.47 (t, *J* = 7.2 Hz, 1H), 2.29 – 2.15 (m, 4H), 1.91 (dd, *J* = 14.8, 7.5 Hz, 2H), 1.83 (t, *J* = 7.1 Hz, 2H), 1.45 (d, *J* = 4.6 Hz, 12H), 1.35 (dd, *J* = 12.7, 5.6 Hz, 18H).

#### Synthesis of compound S5

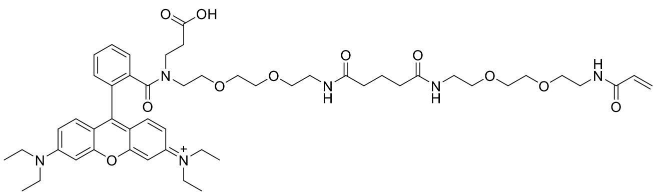

To a solution of the intermediate **S4** in DCM (0.4 mL), TFA (0.4 mL) was added and the reaction mixture was stirred overnight at room temperature. After the reaction finished, all solvents were evaporated to yield the intermediate as a red solid, which was used without further purification.

To a solution of the intermediate (96 mg, 96 μmol) in MeOH: THF (v:v = 0.2 mL: 0.2 mL), a solution of $Na_2CO_3$ (23.3 mg, 219.9 μmol) in water (0.15 mL) was added under the protection of $N_2$, followed by cooling the reaction flask to 0 °C with an ice bath. A solution of acryloyl chloride (10.8 μL, 134 μmol) in dry dioxane (0.1 mL) was added dropwise to the reaction flask, and the reaction mixture was then allowed to come to room temperature over 20 min. After complete reaction, all solvents were evaporated and the residue was purified by column chromatography to yield the product as a red solid (56%). LC-MS ($ESI^+$): calculated for $C_{51}H_{71}N_6O_{11}$ $[M]^+$ m/z: 943.52; found: 943.35; $^1H$ NMR (400 MHz, MeOD-$d_4$) δ 7.85 – 7.66 (m, 3H), 7.54 – 7.45 (m, 1H), 7.30 (dd, *J* = 9.5, 3.8 Hz, 2H), 7.06 (dt, *J* = 9.6, 2.7 Hz, 2H), 6.96 (dd, *J* = 12.4, 2.3 Hz, 2H), 6.37 – 6.15 (m, 2H), 5.64 (dd, *J* = 9.3, 2.7 Hz, 1H), 3.72 – 3.63 (m, 8H), 3.63 – 3.57 (m, 6H), 3.54 (dt, *J* = 11.1, 5.5 Hz, 6H), 3.51 – 3.40 (m, 5H), 3.40 – 3.31 (m, 6H), 3.28 – 3.17 (m, 2H), 3.04 (t, *J* = 5.1 Hz, 1H), 2.44 (t, *J* = 7.4 Hz, 1H), 2.25 – 2.15 (m, 4H), 1.87 (dd, *J* = 14.1, 7.1 Hz, 3H), 1.31 (t, *J* = 7.1 Hz, 12H); $^{13}C$ NMR (101 MHz, MeOD-$d_4$) δ 175.75, 175.14, 172.07, 171.44, 168.49, 166.23, 163.53, 159.56, 157.50, 156.96, 156.74, 137.99, 137.79, 133.78, 133.53, 132.29, 131.86, 131.79, 131.61, 131.51, 131.19, 131.03, 130.56, 129.05, 127.07, 119.97, 117.05, 115.53, 115.01, 114.86, 97.74, 71.86, 71.59, 71.88, 70.78, 69.79, 69.24, 51.48, 47.71, 47.18, 42.01, 41.44, 40.68, 40.65, 40.56, 40.17, 39.18, 36.45, 34.21, 32.23, 23.51, 13.13.

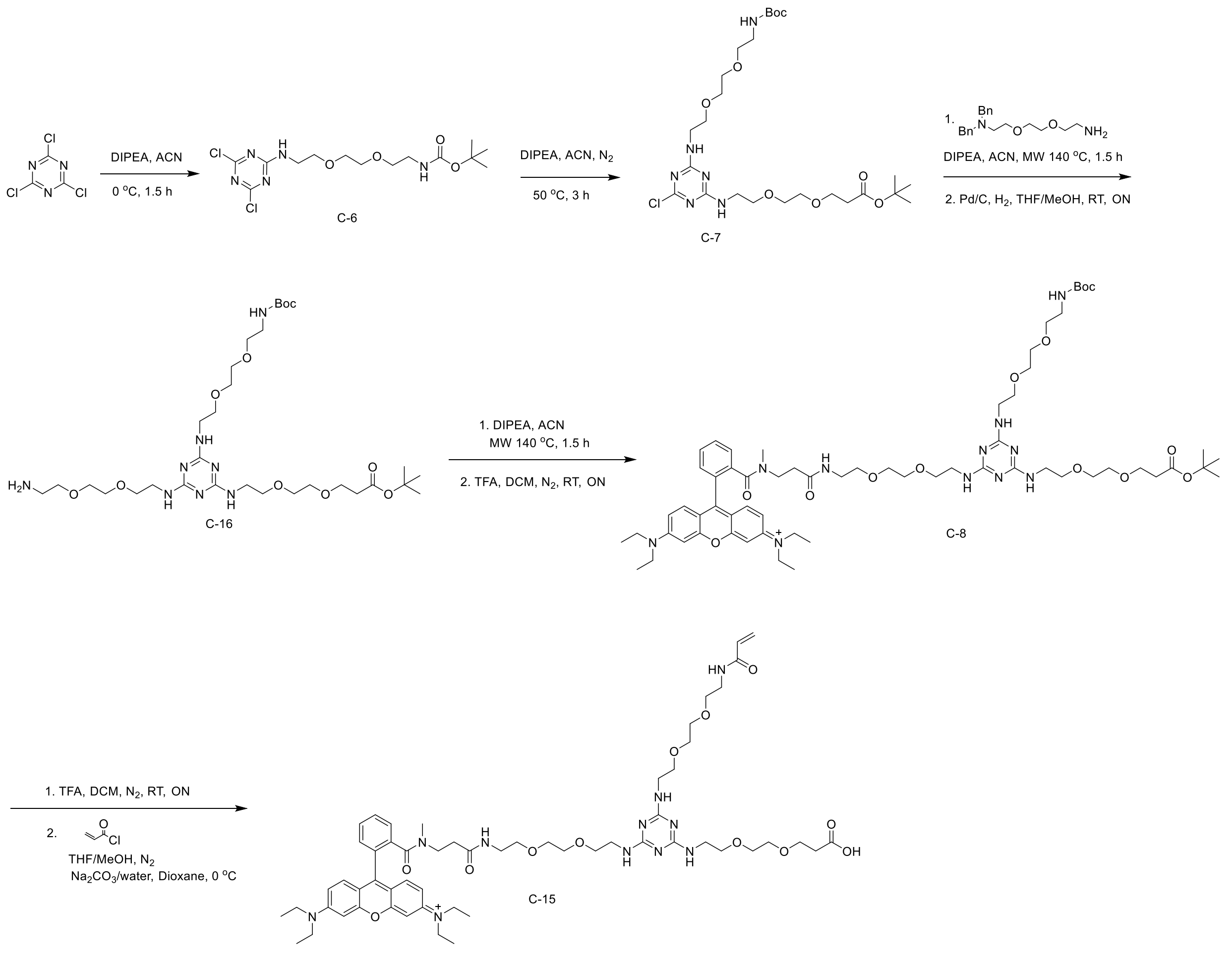

#### Synthesis of compound S6

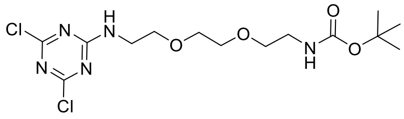

To a solution of N-Boc- 2,2'-(ethylenedioxy)diethylamine (2 g, 8.06 mmol) in anhydrous THF (30 mL), a solution of cyanuric chloride (1.34 g, 7.33 mmol) in anhydrous THF (20 mL) was added at 0 °C with an ice bath. N,N-Diisopropylethylamine (2.46 mL, 14.12 mmol) was then added dropwise to the reaction flask at 0 °C and the reaction mixture was stirred at 0 °C for another 1.5 h. After complete reaction, the white solid was removed by filtration and the filtrate was evaporated. The residue was purified by column chromatography to yield the intermediate as a yellow oil (80%). LC-MS (ESI$^+$): calculated for $C_{14}H_{24}Cl_2N_5O_4$ $[M+H]^+$ m/z: 396.12; found:395.90; $^1$H NMR (300 MHz, $CDCl_3$) $\delta$ 6.72 (brs, 1H), 5.22 (brs, 1H), 3.69 – 3.62 (m, 6H), 3.57 (t, $J$ = 5.1 Hz, 2H), 3.49 (d, $J$ = 4.6 Hz, 2H), 3.40 – 3.30 (m, 2H), 1.44 (s, 9H).

**Synthesis of compound S7**

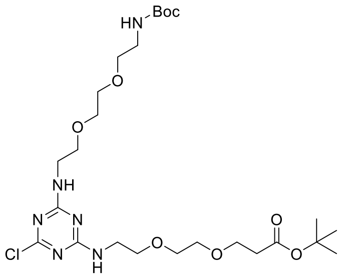

To a solution of the intermediate **S6** (0.358 g, 0.906 mmol) in ACN (22 mL), a solution of 3-(2-(2-Aminoethoxy)ethoxy)propanoic acid t-butyl ester (0.254 g, 1.09 mmol) in ACN (7 mL) was added, followed by adding DIPEA (237 μL, 1.36 mmol) in two portions with a 30 min interval at 0 °C under the protection of $N_2$. The reaction mixture was heated to 55 °C for 3 h. After complete reaction, all solvents were evaporated and the residue was purified by column chromatography to yield the intermediate as a yellow oil (59%). LC-MS (ESI$^+$): calculated for $C_{25}H_{46}ClN_6O_8$ $[M+H]^+$ m/z: 593.31; found: 593.55; $^1$H NMR (300 MHz, $CDCl_3$) δ 5.97 (brs, 1H), 5.78 (brs, 1H), 5.35 (brs, 1H), 3.72 (t, *J* = 6.3 Hz, 2H), 3.62 – 3.52 (m, 16H), 3.49 (d, *J* = 3.6 Hz, 2H), 3.37 – 3.26 (m, 2H), 2.52 (t, *J* = 6.5 Hz, 2H), 1.45 (s, 18H).

**Synthesis of compound S8**

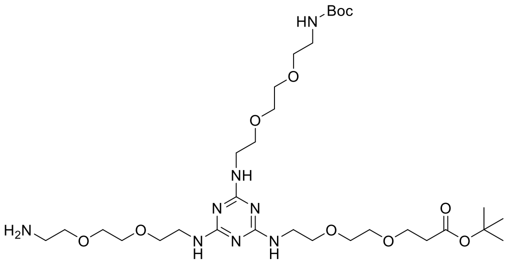

To a solution of the intermediate **S7** (300 mg, 507 μmol) in ACN (4 mL), 2-(2-(2-aminoethoxy)ethoxy)-N,N-dibenzylethan-1-amine (825 mg, 1.48 mmol) and DIPEA (614 μL, 3.55 mmol) were added. The reaction mixture was heated to 140 °C for 1.5 h under microwave. After complete reaction, all solvents were evaporated and the residue was purified by column chromatography to yield the intermediate as a yellow oil (47%).

To a solution of the intermediate (210 mg) in MeOH: THF (v:v = 0.5 mL: 0.5 mL), Pd/C (21 mg) was added and the reaction mixture was stirred overnight at room temperature under the atmosphere of $H_2$. After complete reaction, the black solid was removed by filtration on celite and the filtrate was evaporated to get the intermediate as a yellow oil, of sufficient purity for further use (90%). LC-MS (ESI$^+$): calculated for $C_{31}H_{61}N_8O_{10}$ $[M+H]^+$ m/z: 705.45; found: 705.40; $^1$H NMR (400 MHz, MeOD-*d*$_4$) δ 3.71 (t, *J* = 6.2 Hz, 2H), 3.67 – 3.59 (m, 18H), 3.58 – 3.48 (m, 10H), 3.24 (t, *J* = 5.6 Hz, 2H), 2.83 (t, *J* = 5.3 Hz, 2H), 2.49 (t, *J* = 6.2 Hz, 2H), 1.45 (d, *J* = 7.5 Hz, 18H).

#### Synthesis of compound S9

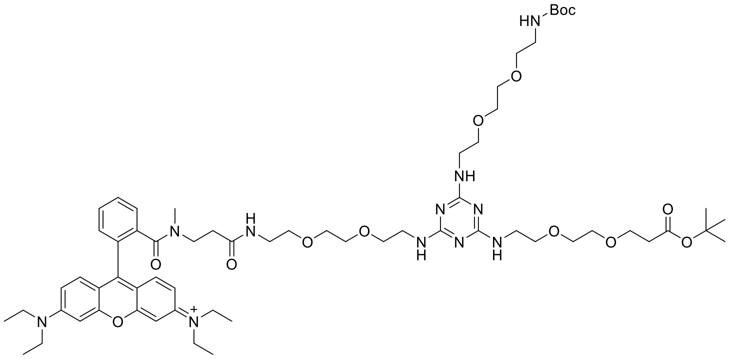

To a solution of the intermediate **S2** (37 mg, 65.5 μmol) in DMF (1.5 mL), trimethylamine (25 μL, 178.9 μmol), HBTU (27 mg, 71.5 μmol) and the intermediate **S8** (42 mg, 59.6 μmol) were added and the reaction mixture was stirred at room temperature for 2 h. After complete reaction, 40 mL ethyl acetate was added into the reaction flask, followed by washing with water (40 mL, 2 x) and brine (40 mL). The organic layer was dried over $MgSO_4$ and then evaporated under reduced pressure. The residue was purified by column chromatography to yield the intermediate as a red solid (75%). LC-MS ($ESI^+$): calculated for $C_{63}H_{96}N_{11}O_{13}$ $[M]^+$ m/z: 1214.72; found: 1214.55; $^1H$ NMR (400 MHz, MeOD-*d*$_4$) δ 7.82 – 7.65 (m, 3H), 7.57 – 7.48 (m, 1H), 7.35 – 7.25 (m, 2H), 7.08 (dd, *J* = 9.5, 2.3 Hz, 2H), 6.96 (dd, *J* = 10.5, 2.3 Hz, 2H), 3.74 – 3.68 (m, 9H), 3.65 – 3.59 (m, 20H), 3.55 – 3.45 (m, 11H), 3.26 (dt, *J* = 15.6, 5.5 Hz, 4H), 2.98 (s, 2H), 2.69 (s, 1H), 2.56 – 2.39 (m, 3H), 2.13 (t, *J* = 6.9 Hz, 1H), 1.47 (d, *J* = 2.7 Hz, 18H), 1.33 (t, *J* = 7.0 Hz, 12H).

#### Synthesis of compound S10

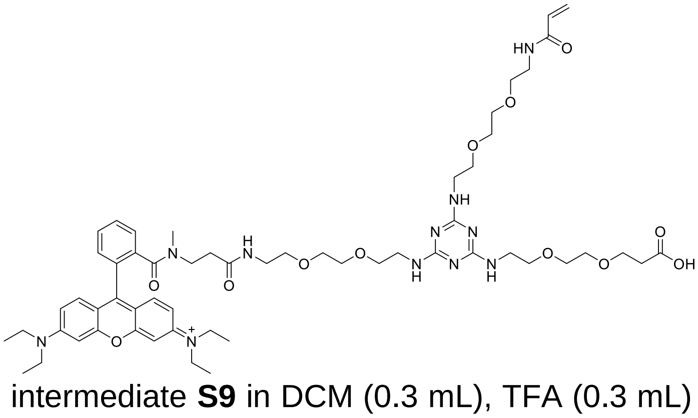

To a solution of the intermediate **S9** in DCM (0.3 mL), TFA (0.3 mL) was added and the reaction mixture was stirred overnight at room temperature. After complete reaction, all solvents were evaporated to yield the intermediate as a red solid, of sufficient purity for further use.

To a solution of the intermediate (60 mg, 51.2 μmol) in MeOH: THF (v:v = 0.3 mL: 0.3 mL), a solution of $Na_2CO_3$ (12.49 mg, 117.9 μmol) in water (0.2 mL) was added under the protection of $N_2$, followed by cooling the reaction flask to 0 °C with an ice bath. A solution of acryloyl chloride (6.6 μL, 82.0 μmol) in dry dioxane (0.2 mL) was added dropwise to the reaction flask, and the reaction mixture was then allowed to come to room temperature over 20 min. After complete reaction, all solvents were evaporated and the residue was purified by column chromatography to yield the product as a red solid (72%). LC-MS ($ESI^+$): calculated for $C_{57}H_{82}N_{11}O_{12}$ $[M]^+$ m/z: 1112.61; found: 1112.45; $^1$H NMR (400 MHz, MeOD-$d_4$) δ 7.81 – 7.63 (m, 3H), 7.54 – 7.47 (m, 1H), 7.28 (dd, *J* = 9.5, 4.9 Hz, 2H), 7.06 (dd, *J* = 9.5, 2.3 Hz, 2H), 6.95 (d, *J* = 2.3 Hz, 2H), 6.34 – 6.12 (m, 2H), 5.64 (dd, *J* = 9.0, 3.0 Hz, 1H), 3.75 – 3.65 (m, 10H), 3.63 – 3.56 (m, 22H), 3.53 – 3.36 (m, 10H), 3.25 (t, *J* = 4.5 Hz, 2H), 2.96 (s, 2H), 2.67 (s, 1H), 2.52 (brs, 2H), 2.42 (t, *J* = 7.0 Hz, 1H), 2.13 (t, *J* = 6.6 Hz, 1H), 1.31 (t, *J* = 7.1 Hz, 12H); $^{13}$C NMR (101 MHz, MeOD-$d_4$) δ 173.43, 170.89, 168.48, 159.58, 157.48, 137.63, 133.50, 132.29 , 132.02, 131.53, 131.30, 129.16, 127.06, 115.50, 115.15, 97.64, 71.65, 71.62, 71.54, 71.55, 70.79, 68.37, 47.19, 45.80, 41.87, 40.70, 38.99, 35.04, 34.46, 27.04, 26.36, 13.16.

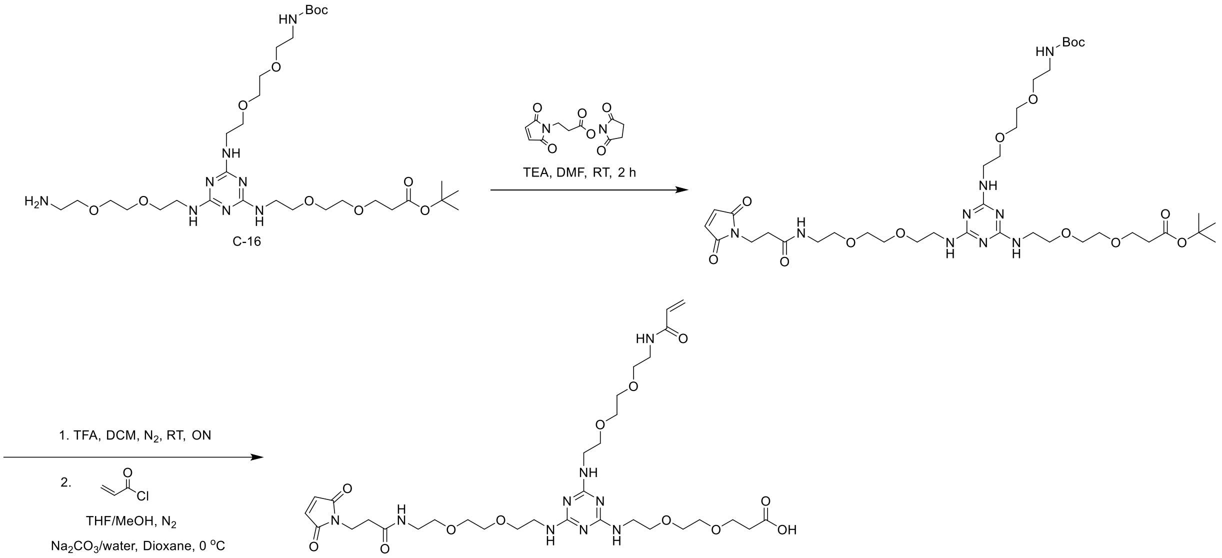

#### Synthesis of compound S11

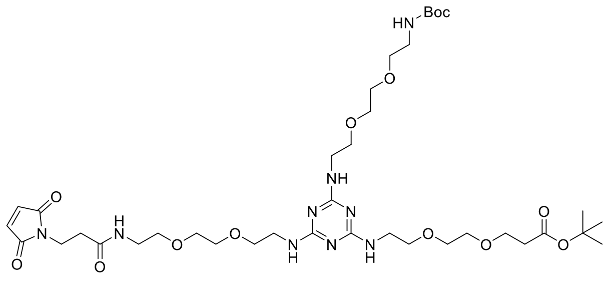

To a solution of the intermediate **S8** (60 mg, 85.2 μmol) in DMF (1.5 mL), trimethylamine (30 μL, 213 μmol), and 3-(Maleimido)propionic acid N-succinimidyl ester (27 mg, 102 μmol) were added and the reaction mixture was stirred at room temperature for 2 h. After complete reaction, 40 mL ethyl acetate was added into the reaction flask, followed by washing with water (40 mL, 2 x) and brine (40 mL). The organic layer was dried over $MgSO_4$ and then evaporated under reduced pressure. The residue was purified by column chromatography to yield the intermediate as a white solid (75%). LC-MS ($ESI^+$): calculated for $C_{38}H_{66}N_9O_{13}$ $[M+H]^+$ m/z: 856.48; found: 856.70; $^1H$ NMR (400 MHz, $CDCl_3$) δ 8.01 (s, 2H), 6.67 (s, 2H), 3.81 (t, *J* = 7.2 Hz, 2H), 3.71 (t, *J* = 6.6 Hz, 2H), 3.65 – 3.49 (m, 28H), 3.42 (dd, *J* = 10.1, 5.2 Hz, 2H), 3.31 (m, 2H), 2.57 – 2.42 (m, 4H), 1.44 (s, 18H).

**Synthesis of compound S12**

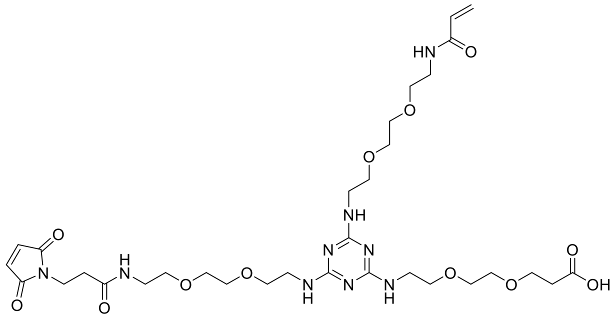

To a solution of the intermediate **S11** in DCM (0.3 mL), TFA (0.3 mL) was added and the reaction mixture was stirred overnight at room temperature. After complete reaction, all solvents were evaporated to yield the intermediate as a pale yellow oil, of sufficient purity for further use.

To a solution of the intermediate (106 mg, 130 μmol) in MeOH: THF (v:v = 0.4 mL: 0.4 mL), a solution of $Na_2CO_3$ (27.6 mg, 260 μmol) in water (0.4 mL) was added under the protection of $N_2$, followed by cooling the reaction flask to 0 °C with an ice bath. A solution of acryloyl chloride (12.5 μL, 155 μmol) in dry dioxane (0.2 mL) was added dropwise to the reaction flask, and the reaction mixture was then allowed to come to room temperature over 20 min. After complete reaction, all solvents were evaporated and the residue was purified by column chromatography to yield the product as a white solid (43%). LC-MS ($ESI^+$): calculated for $C_{32}H_{52}N_9O_{12}$ $[M+H]^+$ m/z: 754.37; found: 754.40; $^1H$ NMR (400 MHz, MeOD-$d_4$) δ 6.82 (s, 2H), 6.36 – 6.10 (m, 2H), 5.65 (dd, *J* = 9.2, 2.8 Hz, 1H), 3.76 (q, *J* = 6.6 Hz, 4H), 3.70 – 3.53 (m, 27H), 3.51 (dd, *J* = 9.3, 3.7 Hz, 3H), 3.45 (t, *J* = 5.5 Hz, 2H), 2.55 – 2.45 (m, 4H); $^{13}C$ NMR (101 MHz, MeOD-$d_4$) δ 173.41, 172.53, 168.50, 163.57, 163.23, 135.79, 132.29, 127.07, 122.83, 119.92, 117.02, 114.11, 71.58, 71.51, 71.30, 70.82, 68.86, 41.76, 40.68, 40.62, 36.02, 35.71.

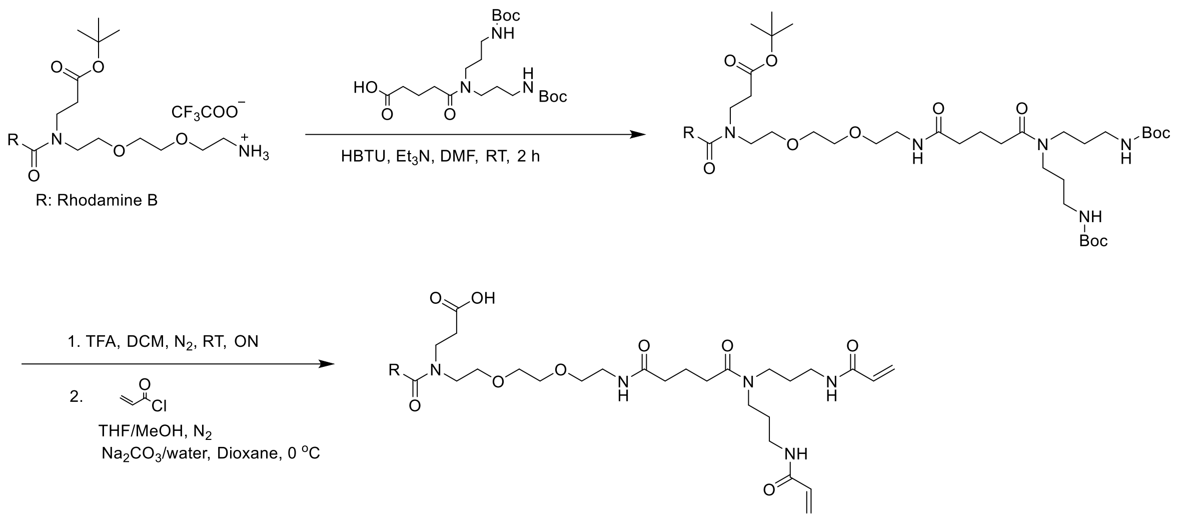

#### Synthesis of compound S13

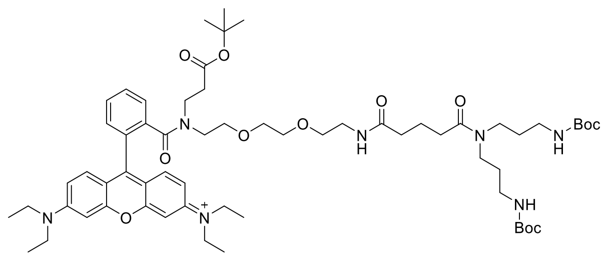

To a solution of the modified rhodamine B (100 mg, 123 μmol, synthesized from the literature[2]) in DMF (1.5 mL), trimethylamine (51 μL, 368 μmol), HBTU (51 mg, 135 μmol) and the acid containing fragment (60 mg, 135 μmol, synthesized from the literature[4]) were added and the reaction mixture was stirred at room temperature for 2 h. After complete reaction, 40 mL ethyl acetate was added into the reaction flask, followed by washing with water (40 mL, 2 x) and brine (40 mL). The organic layer was dried over $MgSO_4$ and then evaporated under reduced pressure. The residue was purified by column chromatography to yield the intermediate as a red solid (61%). LC-MS ($ESI^+$): calculated for $C_{62}H_{94}N_7O_{12}$ $[M]^+$ m/z: 1128.70; found: 1128.40; $^1$H NMR (400 MHz, MeOD-$d_4$) $\delta$ 7.89 – 7.73 (m, 3H), 7.57 – 7.49 (m, 1H), 7.38 – 7.28 (m, 2H), 7.10 (ddd, *J* = 8.8, 6.2, 2.4 Hz, 2H), 7.00 (dd, *J* = 7.6, 2.3 Hz, 2H), 3.78 – 3.68 (m, 8H), 3.68 – 3.61 (m, 2H), 3.61 – 3.52 (m, 4H), 3.50 – 3.42 (m, 3H), 3.38 (dd, *J* = 13.3, 6.4 Hz, 6H), 3.33 – 3.23 (m, 2H), 3.08 (t, *J* = 6.7 Hz, 2H), 3.03 (t, *J* = 6.5 Hz, 3H), 2.47 (t, *J* = 7.1 Hz, 1H), 2.41 (t, *J* = 7.5 Hz, 2H), 2.30 – 2.18 (m, 2H), 1.91 (dd, *J* = 14.6, 7.3 Hz, 2H), 1.84 (t, *J* = 7.2 Hz, 1H), 1.80 – 1.73 (m, 2H), 1.69 (dd, *J* = 13.8, 6.9 Hz, 2H), 1.52 – 1.43 (m, 21H), 1.35 (dd, *J* = 12.6, 5.5 Hz, 18H).

**Synthesis of compound S14**

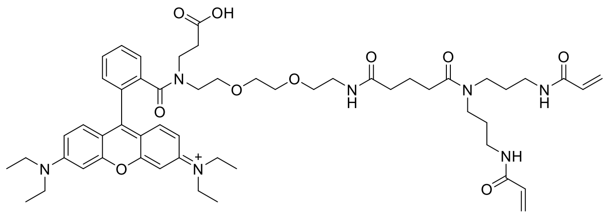

To a solution of the intermediate **S13** in DCM (0.4 mL), TFA (0.4 mL) was added and the reaction mixture was stirred overnight at room temperature. After complete reaction, all solvents were evaporated to yield the intermediate as a red solid, which was used without further purification.

To a solution of the intermediate (85 mg, 75 μmol) in MeOH: THF (v:v = 0.3 mL: 0.3 mL), a solution of $Na_2CO_3$ (31.9 mg, 301 μmol) in water (0.15 mL) was added under the protection of $N_2$, followed by cooling the reaction flask to 0 °C with an ice bath. A solution of acryloyl chloride (16 μL, 195 μmol) in dry dioxane (0.2 mL) was added dropwise to the reaction flask, and the reaction mixture was then allowed to come to room temperature over 20 min. After complete reaction, all solvents were evaporated and the residue was purified by column chromatography to yield the product as a red solid (55%). LC-MS ($ESI^+$): calculated for $C_{54}H_{74}N_7O_{10}$ $[M]^+$ m/z: 980.55; found: 980.25; $^1$H NMR (400 MHz, MeOD-$d_4$) δ 8.00 – 7.68 (m, 3H), 7.56 – 7.45 (m, 1H), 7.31 (dd, $J$ = 9.5, 3.7 Hz, 2H), 7.07 (dd, $J$ = 9.6, 2.4 Hz, 2H), 7.05 – 6.92 (m, 2H), 6.49 – 6.16 (m, 4H), 5.65 (td, $J$ = 7.6, 3.9 Hz, 2H), 3.77 – 3.67 (m, 8H), 3.67 – 3.58 (m, 2H), 3.55 (t, $J$ = 5.4 Hz, 3H), 3.52 – 3.45 (m, 2H), 3.46 – 3.33 (m, 9H), 3.25 (dt, $J$ = 13.8, 6.5 Hz, 5H), 3.06 (t, $J$ = 5.0 Hz, 1H), 2.45 (t, $J$ = 7.5 Hz, 1H), 2.39 (t, $J$ = 7.4 Hz, 2H), 2.31 – 2.19 (m, 2H), 1.93 – 1.80 (m, 5H), 1.76 (dt, $J$ = 14.1, 6.9 Hz, 2H), 1.32 (t, $J$ = 7.1 Hz, 12H); $^{13}$C NMR (101 MHz, MeOD-$d_4$) δ 175.87, 175.06, 172.09, 171.45, 168.54, 168.37, 165.69, 163.51, 159.56, 157.52, 157.00, 138.02, 133.79, 133.55,132.39, 132.24, 131.62, 131.19, 131.03, 130.60, 129.07, 127.17, 126.92, 115.50, 115.03, 114.88, 97.86, 71.88, 71.64, 70.94, 69.83, 69.25, 51.51, 47.71, 47.19, 45.26, 44.97, 42.01, 40.70,40.59, 38.31, 38.16, 36.35, 34.13, 33.45, 32.21, 30.06, 28.83, 23.08, 13.13.

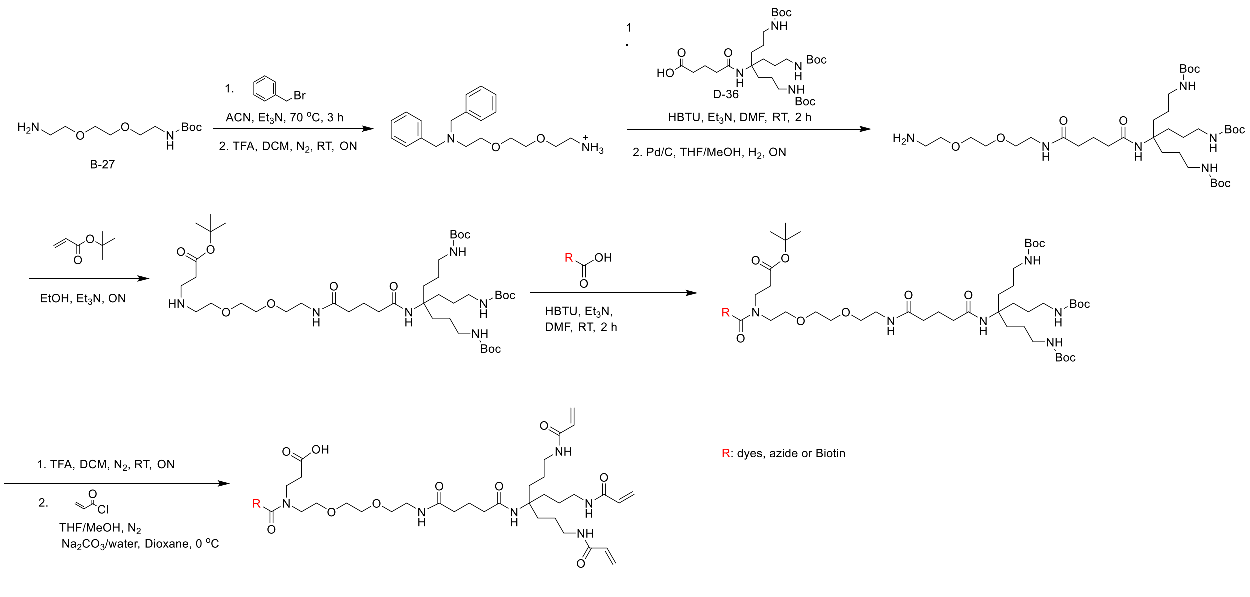

#### Synthesis of compound S15

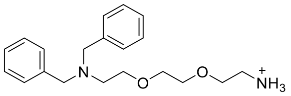

To a solution of N-Boc- 2,2'-(ethylenedioxy)diethylamine (2 g, 8.06 mmol) in anhydrous acetonitrile (10 mL), trimethylamine (2.2 mL, 16.13 mmol) and benzyl bromide (2.1 mL, 17.74 mmol) were added sequentially. The reaction mixture was heated to 70 °C for 3 h. After complete reaction, 300 mL ethyl acetate was added into the reaction flask, followed by washing with water (200 mL, 2 x) and brine (200 mL). The organic layer was dried over $MgSO_4$ and then evaporated under reduced pressure. The residue was purified by column chromatography to yield the intermediate as a colorless oil (33%). LC-MS ($ESI^+$): calculated for $C_{25}H_{37}N_2O_4$ $[M+H]^+$ m/z: 429.27; found: 429.45; $^1$H NMR (300 MHz, $CDCl_3$) $\delta$ 7.41 – 7.18 (m, 11H, including 1H of $CHCl_3$), 4.91 (brs, 1H), 3.65 (s, 4H), 3.62 – 3.43 (m, 8H), 3.28 (dd, $J$ = 10.0, 4.8 Hz, 2H), 2.70 (t, $J$ = 6.1 Hz, 2H), 1.44 (s, 9H).

To a solution of the intermediate (1 g) in DCM (1 mL), TFA (1 mL) was added and the reaction mixture was stirred overnight at room temperature. After complete reaction, all solvents were evaporated to yield the intermediate as a yellow oil, which was used without further purification.

#### Synthesis of compound S16

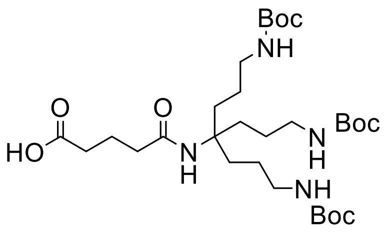

To a solution of the starting material (0.93 g, 1.85 mmol, synthesized from the literature[5]) in dry THF (90 mL) was added glutaric anhydride (0.253 g, 2.22 mmol) and triethylamine (0.52 mL, 3.71 mmol). The reaction mixture was then heated to 70 °C for 3 h. After complete reaction, all solvents were evaporated and the residue was purified by column chromatography to yield the product as a yellow solid (58%). LC-MS (ESI$^+$): calculated for $C_{30}H_{57}N_4O_9$ [M+H]$^+$ m/z: 617.41; found: 617.25; $^1$H NMR (400 MHz, $CDCl_3$) $\delta$ 4.73 (brs, 3H), 3.14 – 2.95 (d, *J* = 5.3 Hz, 6H), 2.46 – 2.34 (m, 2H), 2.32 – 2.18 (m, 2H), 2.02 – 1.94 (m, 2H), 1.80 – 1.57 (m, 6H), 1.46 – 1.34 (m, 33H).

**Synthesis of compound S17**

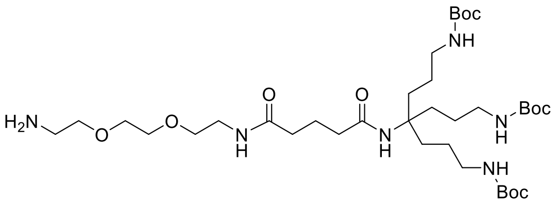

To a solution of compound **S16** (2g, 3.24 mmol) in DMF (5 mL), triethylamine (2.26 mL, 16.2 mmol), HBTU (1.35 g, 3.57 mmol) and the intermediate **S15** (2.16 g, 3.89 mmol) were added and the reaction mixture was stirred at room temperature for 2 h. After complete reaction, 300 mL ethyl acetate was added into the reaction flask, followed by washing with water (200 mL, 2 x) and brine (200 mL). The organic layer was dried over $MgSO_4$ and then evaporated under reduced pressure. The residue was purified by column chromatography to yield the intermediate as a white solid (70%).

To a solution of the intermediate (2.1 g) in MeOH: THF (v:v = 4.2 mL: 4.2 mL), Pd/C (0.2 g) was added and the reaction mixture was stirred overnight at room temperature under the atmosphere of $H_2$. After complete reaction, the black solid was removed by filtration on celite and the filtrate was evaporated to get the intermediate as a yellow oil, of sufficient purity for further use (96%). LC-MS (ESI$^+$): calculated for $C_{36}H_{71}N_6O_{10}$ [M+H]$^+$ m/z: 747.52; found: 747.35; $^1$H NMR (400 MHz, $CDCl_3$) $\delta$ 6.88 (brs, 1H), 5.74 (s, 1H), 4.85 (brs, 3H), 3.71 – 3.61 (m, 4H), 3.58 (t, *J* = 5.0 Hz, 4H), 3.44 (dd, *J* = 10.4, 5.2 Hz, 2H), 3.20 – 3.00 (m, 6H), 2.93 (brs, 2H), 2.44 (brs, 3H), 2.26 (t, *J* = 6.9 Hz, 2H), 2.19 (t, *J* = 6.9 Hz, 2H), 1.92 – 1.86 (m, 2H), 1.63 (t, *J* = 14.2 Hz, 6H), 1.54 – 1.22 (m, 33H).

**Synthesis of compound S18**

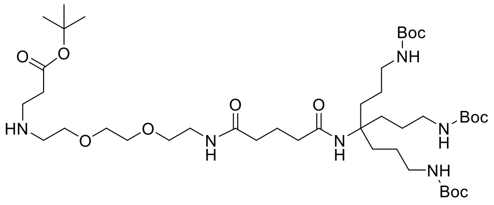

To a solution of the intermediate **S17** (1.79 g, 2.40 mmol) in EtOH (25 mL), tert-butyl acrylate (0.351 mL, 2.42 mmol) and triethylamine (0.333 mL, 2.40 mmol) were added and the reaction mixture was stirred overnight at RT. After complete reaction, all solvents were removed under reduced pressure and the residue was purified by column chromatography to obtain the intermediate as a yellow solid (62%). LC-MS (ESI$^+$): calculated for $C_{43}H_{83}N_6O_{12}$ $[M+H]^+$ m/z: 875.61; found: 875.50; $^1$H NMR (400 MHz, $CDCl_3$) δ 7.14 (brs, 1H), 5.82 (brs, 1H), 4.82 (brs, 3H), 3.85 – 3.74 (m, 2H), 3.71 – 3.64 (m, 2H), 3.64 – 3.60 (m, 2H), 3.58 (t, *J* = 5.0 Hz, 2H), 3.43 (dd, *J* = 10.3, 5.3 Hz, 2H), 3.07 (d, *J* = 6.0 Hz, 10H), 2.72 (brs, 2H), 2.29 (t, *J* = 6.9 Hz, 2H), 2.21 (t, *J* = 6.9 Hz, 2H), 2.00 – 1.88 (m, 2H), 1.78 – 1.60 (m, 6H), 1.44 (d, *J* = 8.2 Hz, 42H).

**Synthesis of compound S19**

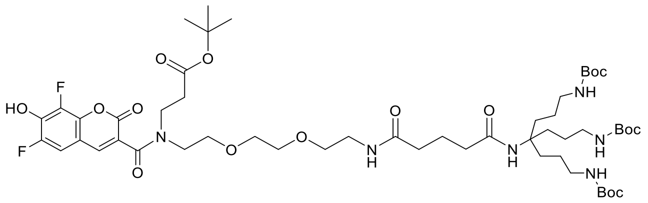

To a solution of pacific blue (30 mg, 125 μmol) in DMF (1.5 mL), trimethylamine (47 μL, 340 μmol), HBTU (43 mg, 113 μmol) and the compound **S18** (99 mg, 113 μmol) were added and the reaction mixture was stirred at room temperature for 2 h. After complete reaction, 40 mL ethyl acetate was added into the reaction flask, followed by washing with water (40 mL, 2 x) and brine (40 mL). The organic layer was dried over $MgSO_4$ and then evaporated under reduced pressure. The residue was purified by column chromatography to yield the intermediate as a pale green solid (40%). LC-MS (ESI$^+$): calculated for $C_{53}H_{85}F_2N_6O$ $[M+H]^+$ m/z: 1099.60; found: 1099.55; $^1$H NMR (400 MHz, MeOD-$d_4$) δ 7.96 (s, 1H), 7.25 (t, *J* = 9.8 Hz, 1H), 7.10 (s, 1H), 3.80 – 3.50 (m, 12H), 3.40 – 3.35 (m, 2H), 3.00 (t, *J* = 6.7 Hz, 6H), 2.74 – 2.57 (m, 2H), 2.28 – 2.12 (m, 4H), 1.95 – 1.78 (m, 2H), 1.63 (t, *J* = 14.6 Hz, 6H), 1.50 – 1.36 (m, 42H).

**Synthesis of compound S20**

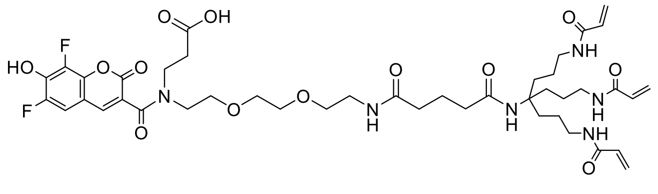

To a solution of the intermediate **S19** in DCM (0.3 mL), TFA (0.3 mL) was added and the reaction mixture was stirred overnight at room temperature. After complete reaction, all solvents were evaporated to yield the intermediate as a green solid, which was used without further purification.

To a solution of the intermediate (64 mg, 59 μmol) in MeOH: THF (v:v = 1 mL: 1 mL), a solution of $Na_2CO_3$ (43.8 mg, 413 μmol) in water (1 mL) was added under the protection

of $N_2$, followed by cooling the reaction flask to 0 °C with an ice bath. A solution of acryloyl chloride (23 μL, 283 μmol) in dry dioxane (0.5 mL) was added dropwise to the reaction flask, and the reaction mixture was then allowed to come to room temperature over 20 min. After complete reaction, all solvents were evaporated and the residue was purified by column chromatography to yield the product as a green solid (37%). LC-MS (ESI$^+$): calculated for $C_{43}H_{59}F_2N_6O_{13}$ $[M+H]^+$ m/z: 905.41; found: 905.30; $^1$H NMR (600 MHz, MeOD-*d*4) δ 7.91 (s, 1H), 7.15 – 7.09 (m, 1H), 6.30 – 6.12 (m, 6H), 5.63 (dd, *J* = 9.4, 2.6 Hz, 3H), 3.81 – 3.73 (m, 2H), 3.72 – 3.60 (m, 5H), 3.60 – 3.53 (m, 5H), 3.39 – 3.36 (m, 2H), 3.21 (t, *J* = 7.0 Hz, 6H), 2.70 (t, *J* = 7.0 Hz, 1H), 2.64 – 2.58 (m, 1H), 2.24 – 2.18 (m, 2H), 2.19 – 2.13 (m, 2H), 1.88 – 1.82 (m, 2H), 1.73 – 1.63 (m, 6H), 1.50 – 1.40 (m, 6H); $^{13}$C NMR (151 MHz, MeOD-*d*4) δ 176.60, 175.87, 175.16, 169.21, 168.46, 160.50, 145.71, 143.04, 142.98, 132.47, 126.81, 109.69, 94.28, 71.80, 71.60. 70.90, 70.35, 64.72, 59.66, 57.91, 51.16, 47.90, 46.77, 44.12, 40.97, 40.69, 37.27, 36.55, 33.98, 33.34, 31.05, 24.55, 23.69.
Other analogues were synthesized and purified based on the same method as described above, using different dyes and azide/biotin linkers.

**Synthesis of compound S21**

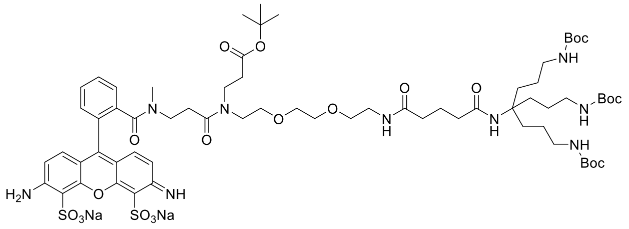

Yellow solid; Yield: 38%; Purity: 80%, quantified by liquid chromatography of LC-MS; LC-MS (ESI$^+$): calculated for $C_{67}H_{102}N_9O_{21}S_2$ $[M+H]^+$ m/z: 1432.66; found: 1432.75; Due to the small amount, no NMR data was performed.

**Synthesis of compound S22**

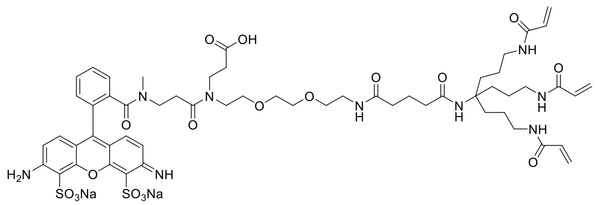

Yellow solid; Yield: 25%; Purity: 95%, quantified by liquid chromatography of LC-MS; LC-MS (ESI$^+$): calculated for $C_{57}H_{76}N_9O_{18}S_2$ $[M+H]^+$ m/z: 1238.47; found: 1238.50; Due to the small amount, no NMR data was performed.

#### Synthesis of compound S23

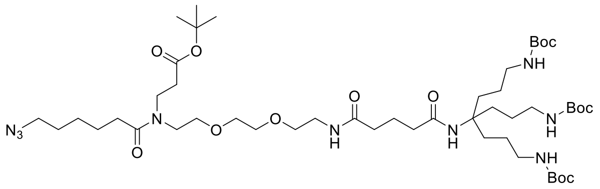

Yellow solid; Yield: 86%; LC-MS (ESI+): calculated for $C_{49}H_{92}N_9O_{13}$ $[M+H]^+$ m/z: 1014.68; found: 1014.50; $^1$H NMR (400 MHz, $CDCl_3$) δ 5.61 (s, 1H), 4.78 (brs, 2H), 3.67 – 3.48 (m, 12H), 3.43 (dt, *J* = 9.4, 4.8 Hz, 2H), 3.28 (td, *J* = 6.8, 1.6 Hz, 2H), 3.18 – 2.85 (m, 6H), 2.52 (td, *J* = 7.2, 4.0 Hz, 2H), 2.37 (td, *J* = 7.5, 4.0 Hz, 2H), 2.32 – 2.21 (m, 2H), 2.18 (t, *J* = 6.8 Hz, 2H), 1.98 – 1.84 (m, 2H), 1.81 – 1.52 (m, 10H), 1.52 – 0.87 (m, 44H).

#### Synthesis of compound S24

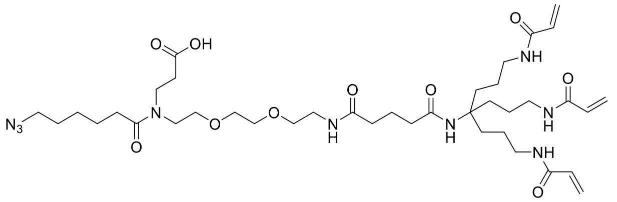

Pale yellow solid; Yield: 30%; LC-MS (ESI+): calculated for $C_{39}H_{66}N_9O_{10}$ $[M+H]^+$ m/z: 820.49; found: 820.30; $^1$H NMR (400 MHz, MeOD-*d*4) δ 6.31 – 6.12 (m, 6H), 5.64 (dd, *J* = 8.6, 3.4 Hz, 3H), 3.72 – 3.58 (m, 10H), 3.57 – 3.50 (m, 3H), 3.40 – 3.35 (m, 3H), 3.29 – 3.17 (m, 6H), 2.69 – 2.55 (m, 2H), 2.50 – 2.41 (m, 2H), 2.30 – 2.14 (m, 4H), 1.90 – 1.81 (m, 2H), 1.74 – 1.66 (m, 6H), 1.66 – 1.57 (m, 4H), 1.53 – 1.37 (m, 8H); $^{13}$C NMR (151 MHz, MeOD-*d*4) δ 175.85, 175.13, 168.43, 163.51, 163.28, 132.45, 126.83, 72.04, 71.67, 70.97, 70.69, 70.37, 59.66, 52.63, 50.15, 47.36, 44.52, 40.96, 40.63, 37.25, 36.53, 34.20, 34.03, 33.31, 30.05, 27.80, 26.35, 26.23, 24.54, 23.68.

#### Synthesis of compound S25

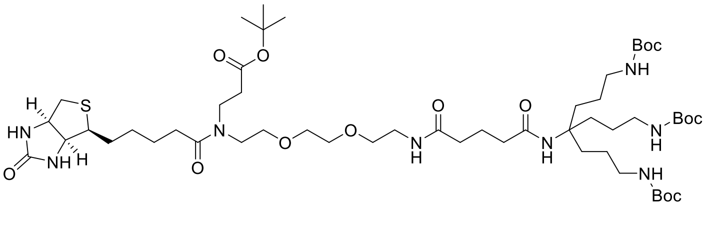

Pale yellow solid; Yield: 62%; LC-MS (ESI+): calculated for $C_{53}H_{97}N_8O_{14}S$ $[M+H]^+$ m/z: 1101.68; found: 1101.55; $^1$H NMR (400 MHz, $CDCl_3$) δ 6.86 (brs, 1H), 4.92 (brs, 2H), 4.59 – 4.49 (m, 1H), 4.40 – 4.30 (m, 1H), 3.70 – 3.46 (m, 13H), 3.46 – 3.35 (m, 2H), 3.20 (dd, *J* = 11.8, 7.1 Hz, 1H), 3.16 – 3.00 (d, *J* = 5.0 Hz, 6H), 2.94 (dd, *J* = 12.8, 4.9 Hz, 1H), 2.79 (d, *J* = 12.8 Hz, 1H), 2.60 – 2.48 (m, 2H), 2.48 – 2.35 (m, 2H), 2.32 – 2.17 (m, 4H), 2.00 – 1.86 (m, 3H), 1.74 – 1.60 (m, 8H), 1.51 – 1.34 (m, 44H).

**Synthesis of compound S26**

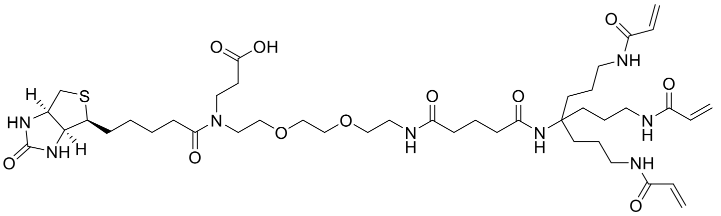

White solid; Yield: 25%; LC-MS (ESI$^+$): calculated for $C_{43}H_{71}N_8O_{11}S$ $[M+H]^+$ m/z: 907.50; found: 907.25; $^1H$ NMR (400 MHz, MeOD-*d*$_4$) $\delta$ 6.28 – 6.15 (m, 6H), 5.64 (dd, *J* = 8.2, 3.8 Hz, 3H), 4.50 (dd, *J* = 7.8, 4.8 Hz, 1H), 4.32 (dd, *J* = 7.8, 4.4 Hz, 1H), 3.77 – 3.70 (m, 1H), 3.69 – 3.57 (m, 9H), 3.54 (t, *J* = 5.6 Hz, 3H), 3.40 – 3.34 (m, 3H), 3.22 (t, *J* = 6.9 Hz, 7H), 2.93 (dd, *J* = 12.7, 5.0 Hz, 1H), 2.75 – 2.56 (m, 3H), 2.47 (dt, *J* = 12.0, 7.4 Hz, 2H), 2.20 (dt, *J* = 18.3, 7.4 Hz, 4H), 1.90 – 1.79 (m, 2H), 1.74 – 1.63 (m, 8H), 1.53 – 1.41 (m, 8H); $^{13}C$ NMR (101 MHz, MeOD-*d*$_4$) $\delta$ 174.69, 174.14, 173.41, 166.73, 130.74, 125.17, 70.38, 70.02, 69.27, 68.98, 61.93, 60.24, 57.95, 55.64, 42.72, 39.67, 39.27 38.97, 35.57, 34.86, 32.40, 32.20, 31.96, 31.61, 28.50, 28.19, 24.91, 22.02.

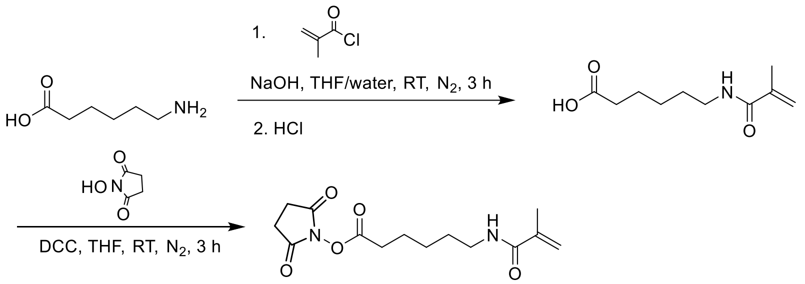

**Synthesis of compound S27**

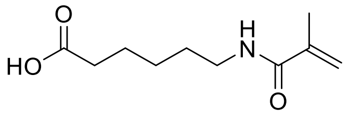

To a solution of 6-aminohexanoic acid (2.62 g, 20 mmol) in 1,4-dioxane/water (v:v =100 mL:100 mL), sodium hydroxide (0.8 g, 20 mmol) was added, followed by cooling to 0 °C with an ice bath. A solution of methacryloyl chloride (3.14 g, 30 mmol) in 1,4-dioxane (30 mL) was added dropwise and the resulting mixture was stirred at room temperature for 12 h. After complete reaction, the pH was adjusted to 3 by addition of HCl (1 M), followed by extraction with dichloromethane (200 mL). Afterwards, the organic phase was then washed with water (100 mL) and brine (100 mL). The organic layer was dried over $MgSO_4$ and then evaporated under reduced pressure. The residue was purified by column chromatography to yield the intermediate as a white oil (50%). LC-MS (ESI$^+$): calculated for $C_{10}H_{18}NO_3$ $[M+H]^+$ m/z: 200.13; found: 199.80; $^1H$ NMR (400 MHz, $CDCl_3$) $\delta$ 5.93 (brs,

1H), 5.67 (s, 1H), 5.34 – 5.30 (m, 1H), 3.31 (dd, *J* = 13.1, 7.1 Hz, 2H), 2.35 (t, *J* = 7.3 Hz, 2H), 1.95 (s, 3H), 1.70 – 1.62 (m, 2H), 1.56 (dt, *J* = 14.8, 7.4 Hz, 2H), 1.43 – 1.34 (m, 2H).

**Synthesis of compound 7**

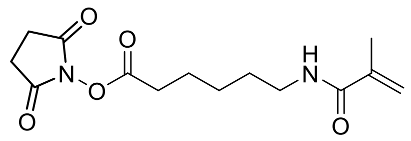

To a solution of the intermediate **S27** (570 mg, 2.86 mmol) in THF (40 mL), N-Hydroxysuccinimide (362 mg, 3.15 mmol) was added, followed by cooling to 0 °C with an ice bath. A solution of DCC (2.36 g, 11.45 mmol) in THF (20 mL) was added dropwise and the resulting mixture was stirred at room temperature for 2 h. After complete reaction, the mixture was filtered to remove the white precipitate and the filtrate was evaporated under reduced pressure. The residue was further triturated with EtOAc (6 mL) in an ice bath. The mixture was filtered again to remove the precipitate. The filtrate was evaporated under reduced pressure and the residue was purified by column chromatography to yield the product as a white solid (63%). LC-MS (ESI$^+$): calculated for $C_{14}H_{21}N_2O_5$ $[M+H]^+$ m/z: 297.14; found: 296.80; $^1$H NMR (400 MHz, $CDCl_3$) $\delta$ 6.02 (brs, 1H), 5.65 (s, 1H), 5.29 – 5.26 (m, 1H), 3.29 (dd, *J* = 12.9, 6.9 Hz, 2H), 2.81 (s, 4H), 2.59 (t, *J* = 7.2 Hz, 2H), 1.96 – 1.87 (m, 3H), 1.83 – 1.71 (m, 2H), 1.62 – 1.51 (m, 2H), 1.50 – 1.39 (m, 2H); $^{13}$C NMR (101 MHz, $CDCl_3$) $\delta$ 169.34, 168.62, 168.60, 140.24, 119.35, 39.35, 30.94, 29.02, 26.01, 24.32, 19.13, 18.80.

**Synthesis of compound 8**

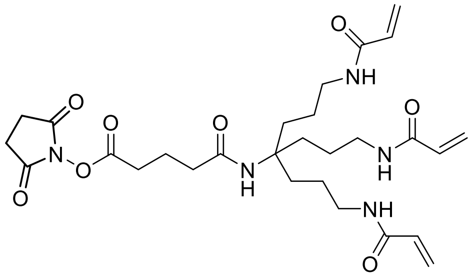

To a solution of the intermediate (61 mg, 0.128 mmol, synthesized from the literature[2]) in THF (6 mL), N-Hydroxysuccinimide (22 mg, 0.191 mmol) was added, followed by cooling to 0 °C with an ice bath. A solution of DCC (132 mg, 0.638 mmol) in THF (5 mL) was added dropwise and the resulting mixture was stirred at room temperature for 2 h. After complete reaction, the mixture was filtered to remove the white precipitate and the filtrate was evaporated under reduced pressure. The residue was further triturated with EtOAc (3 mL) in an ice bath. The mixture was filtered again to remove the precipitate and the filtrate was evaporated under reduced pressure. This procedure was repeated three times and the residue was used for coupling without further purification. LC-MS (ESI$^+$): calculated for $C_{28}H_{42}N_5O_8$ $[M+H]^+$ m/z: 576.30; found: 576.00.

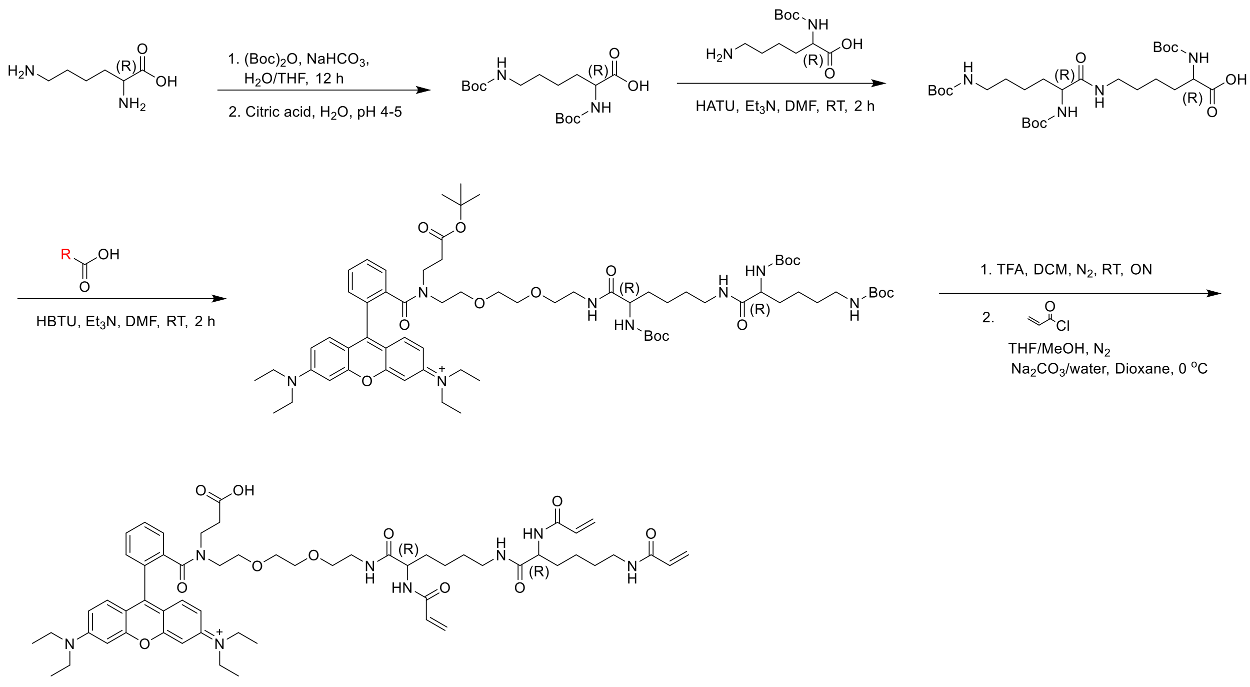

#### Synthesis of compound S28

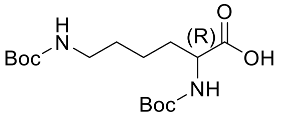

To a solution of (R)-2,6-Diaminocaproic acid (2.5 g, 17 mmol) in water (31 mL), $NaHCO_3$ (4.31 g, 51.3 mmol) was added under the protection of $N_2$, followed by cooling the reaction flask to 0 ℃ with an ice bath. A solution of $(Boc)_2O$ (4.48 g, 20.5 mmol) in THF (31 mL) was then added dropwise to the reaction flask and the reaction mixture was stirred at room temperature for 2 h. And the same amount of $(Boc)_2O$ (4.48 g, 20.5 mmol) was added and the reaction mixture was stirred at room temperature for additional 2 h. After complete reaction, THF was evaporated and the aqueous layer was acidified to pH 4-5 with a solution of 10% citric acid in water, followed by extraction with 300 mL DCM. The organic layer was washed with brine, dried over $MgSO_4$ and evaporated under reduced pressure. The residue was purified by column chromatography to yield the intermediate as a white solid (73%). LC-MS (ESI$^+$): calculated for $C_{16}H_{29}N_2O_6$ [M-H]$^-$ m/z: 345.20; found: 344.90; $^1$H NMR (400 MHz, DMSO-$d_4$) δ 12.39 (brs, 1H), 6.99 (d, *J* = 7.9 Hz, 1H), 6.76 (t, *J* = 5.2 Hz, 1H), 3.81 (td, *J* = 9.0, 4.8 Hz, 1H), 2.92 – 2.85 (m, 2H), 1.63 – 1.48 (m, 2H), 1.42 – 1.31 (m, 22H).

**Synthesis of compound S29**

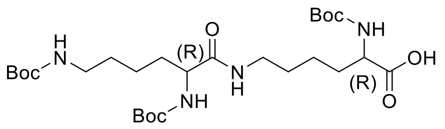

To a solution of the intermediate **S28** (736 mg, 2.13 mmol) in DMF (2 mL), trimethylamine (0.89 mL, 6.38 mmol) and HATU (849 mg, 2.23 mmol) were added and the reaction mixture was stirred at room temperature for 30 min. A solution of $N^{\alpha}$-Boc-D-lysine (550 mg, 2.23 mmol) in DMF (1 mL) was added to the reaction flask and the resulting mixture was stirred at room temperature for 1 h. After complete reaction, all solvents were evaporated and the residue was dissolved in 70 mL water. And the aqueous solution was adjusted to pH 5 by using citric acid, followed by extraction with DCM (150 mL). The organic layer was then washed with brine (100 mL, 2 x), dried over $MgSO_4$ and evaporated under reduced pressure. The residue was purified by column chromatography to yield the intermediate as a white solid (36%). LC-MS (ESI$^+$): calculated for $C_{27}H_{49}N_4O_9$ $[M-H]^-$ m/z: 573.35; found: 573.05; $^1$H NMR (400 MHz, DMSO-$d_6$) $\delta$ 12.46 (brs, 1H), 7.73 (t, *J* = 5.5 Hz, 1H), 6.99 (d, *J* = 7.9 Hz, 1H), 6.80 – 6.60 (m, 2H), 3.87 – 3.63 (m, 2H), 3.08 – 2.93 (m, 2H), 2.91 – 2.78 (m, 2H), 1.69 – 1.40 (m, 4H), 1.39 – 1.15 (m, 35H).

**Synthesis of compound S30**

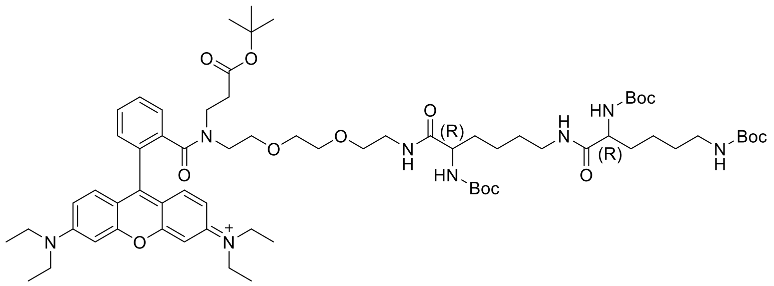

To a solution of the modified rhodamine B (145.3 mg, 178 μmol, synthesized from the literature[2]) in DMF (1 mL), trimethylamine (68 μL, 486 μmol), HBTU (68 mg, 178 μmol) and the modified D-lysine intermediate **S29** (93 mg, 162 μmol) were added and the reaction mixture was stirred at room temperature for 2 h. After complete reaction, 50 mL ethyl acetate was added into the reaction flask, followed by washing with water (50 mL, 2 x) and brine (40 mL). The organic layer was dried over $MgSO_4$ and then evaporated under reduced pressure. The residue was purified by column chromatography to yield the intermediate as a red solid (85%). LC-MS (ESI$^+$): calculated for $C_{68}H_{105}N_8O_{14}$ $[M]^+$ m/z: 1257.77; found: 1257.55; $^1$H NMR (400 MHz, MeOD-$d_4$) $\delta$ 7.86 – 7.66 (m, 3H), 7.55 – 7.44 (m, 1H), 7.31 (dd, *J* = 9.5, 3.4 Hz, 2H), 7.14 – 7.04 (m, 2H), 6.98 (t, *J* = 3.8 Hz, 2H), 4.05 – 3.80 (m, 2H), 3.78 – 3.64 (m, 8H), 3.62 (dd, *J* = 5.8, 2.9 Hz, 1H), 3.60 – 3.48 (m, 4H), 3.48 – 3.33 (m, 6H), 3.29 – 3.23 (m, 2H), 3.23 – 3.08 (m, 2H), 3.08 – 2.95 (m, 3H),

2.45 (t, *J* = 7.2 Hz, 1H), 1.81 (t, *J* = 7.1 Hz, 1H), 1.78 – 1.63 (m, 2H), 1.62 – 1.54 (m, 2H), 1.53 – 1.35 (m, 38H), 1.33 (dd, *J* = 12.6, 5.5 Hz, 18H).

**Synthesis of compound S31**

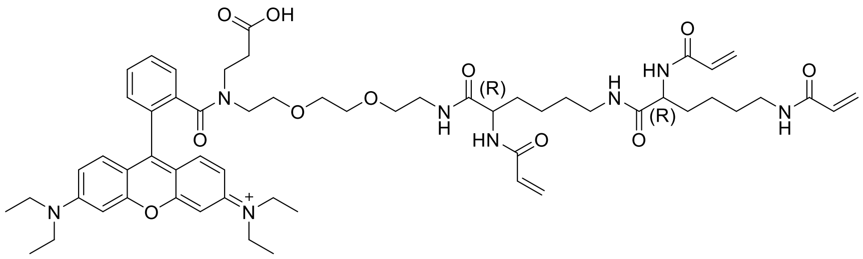

To a solution of the intermediate **S30** in DCM (0.3 mL), TFA (0.3 mL) was added and the reaction mixture was stirred overnight at room temperature. After complete reaction, all solvents were evaporated to yield the intermediate as a red solid, which was used without further purification.

To a solution of the intermediate (180 mg, 145 μmol) in MeOH: THF (v:v = 0.8 mL: 0.8 mL), a solution of $Na_2CO_3$ (115 mg, 1.086 mmol) in water (0.7 mL) was added under the protection of $N_2$, followed by cooling the reaction flask to 0 ℃ with an ice bath. A solution of acryloyl chloride (52.7 μL, 651 μmol) in dry dioxane (0.5 mL) was added dropwise to the reaction flask, and the reaction mixture was then allowed to come to room temperature over 20 min. After complete reaction, all solvents were evaporated and the residue was purified by column chromatography to yield the product as a red solid (52%). LC-MS ($ESI^+$): calculated for $C_{58}H_{79}N_8O_{11}$ $[M]^+$ m/z: 1063.59; found: 1063.35; $^1$H NMR (400 MHz, MeOD-*d*$_4$) δ 7.85 – 7.67 (m, 3H), 7.53 – 7.46 (m, 1H), 7.32 (d, *J* = 9.5 Hz, 2H), 7.13 – 7.03 (m, 2H), 6.97 (dd, *J* = 11.6, 2.2 Hz, 2H), 6.84 – 6.16 (m, 6H), 5.90 – 5.52 (m, 3H), 4.45 – 4.08 (m, 2H), 3.78 – 3.66 (m, 8H), 3.66 – 3.58 (m, 2H), 3.58 – 3.51 (m, 3H), 3.49 (t, *J* = 4.8 Hz, 1H), 3.47 – 3.32 (m, 6H), 3.30 – 3.18 (m, 4H), 3.10 (dt, *J* = 10.1, 5.9 Hz, 2H), 2.42 (t, *J* = 7.4 Hz, 1H), 1.88 (t, *J* = 6.7 Hz, 1H), 1.84 – 1.74 (m, 2H), 1.73 – 1.62 (m, 2H), 1.59 – 1.46 (m, 4H), 1.43 – 1.25 (m, 16H); $^{13}$C NMR (101 MHz, MeOD-*d*$_4$) δ 175.29, 174.46, 174.30, 171.90, 171.19, 168.23, 167.20, 159.36, 157.31, 156.70, 137.88, 137.67, 133.62, 133.36, 132.22, 131.92, 131.84, 131.65, 131.56, 131.43, 131.34, 131.27, 130.94, 130.81, 130.43, 128.82, 127.31, 126.63, 115.36, 114.85, 114.68, 97.66, 97.40, 71.70, 71.50, 71.39, 70.65, 69.54, 68.98, 55.14, 54.98, 51.28, 46.99, 44.99, 41.91, 40.56, 40.56, 40.14, 34.96, 32.96, 32.36, 30.10, 30.03, 24.43, 24.32, 24.27, 12.95.

Other analogues were synthesized and purified based on the same method as described above, using different D-lysine fragments.

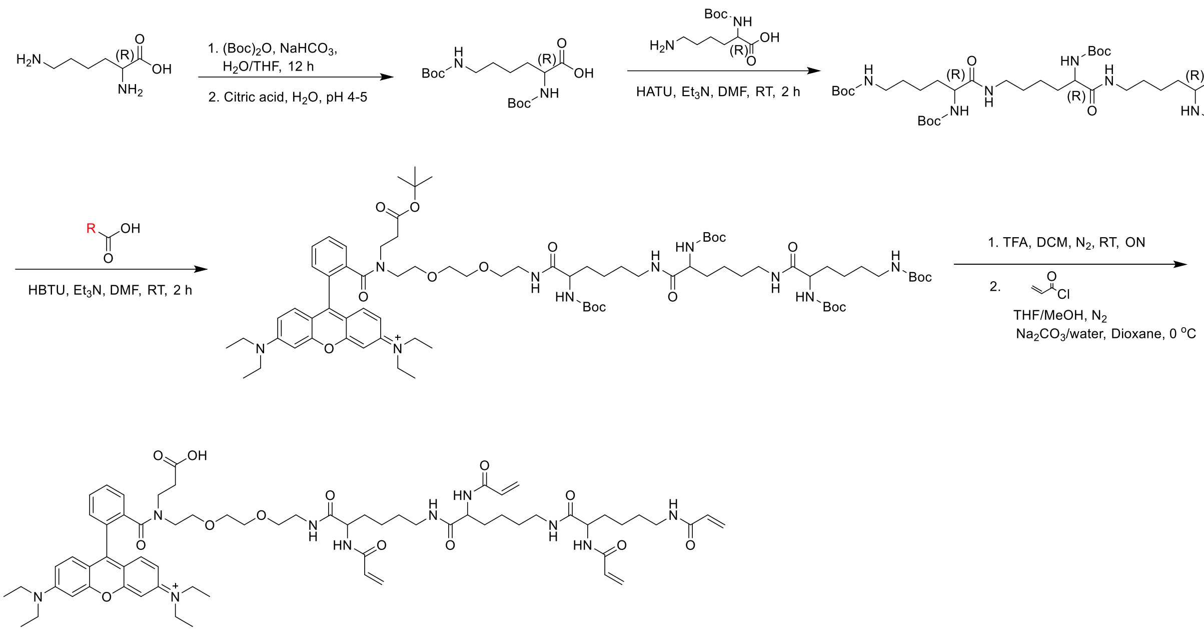

#### Synthesis of compound S32

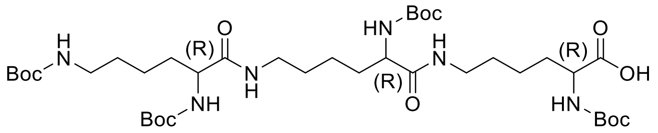

To a solution of the intermediate **S28** (865 mg, 2.50 mmol) in DMF (2 mL), trimethylamine (2.43 mL, 17.49 mmol) and HATU (1.99 g, 5.25 mmol) were added and the reaction mixture was stirred at room temperature for 30 min. A solution of $N^{\alpha}$-Boc-D-lysine (1.29 g, 5.25 mmol) in DMF (2 mL) was added to the reaction flask and the resulting mixture was stirred at room temperature for 1 h. After complete reaction, all solvents were evaporated and the residue was dissolved in 70 mL water. And the aqueous solution was adjusted to pH 5 by using citric acid, followed by extraction with DCM (170 mL). The organic layer was then washed with brine (100 mL, 2 x), dried over $MgSO_4$ and evaporated under reduced pressure. The residue was purified by column chromatography to yield the intermediate as a white solid (23%). LC-MS (ESI$^+$): calculated for $C_{38}H_{69}N_6O_{12}$ [M-H]$^-$ m/z: 801.50; found: 801.15; $^1$H NMR (400 MHz, DMSO-$d_6$) $\delta$ 12.40 (brs, 1H), 7.81 – 7.63 (m, 2H), 6.95 (d, $J$ = 7.6 Hz, 1H), 6.83 – 6.51 (m, 3H), 3.90 – 3.65 (m, 3H), 3.07 – 2.92 (m, 4H), 2.87 – 2.76 (m, 2H), 1.64 – 1.20 (m, 54H).

**Synthesis of compound S33**

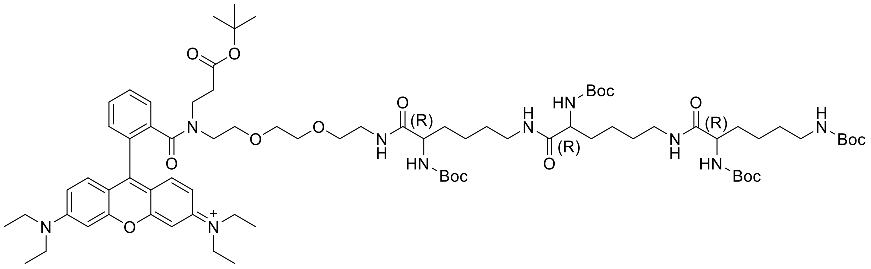

To a solution of the modified rhodamine B (60 mg, 74.1 μmol, synthesized from the literature[2]) in DMF (1 mL), trimethylamine (28.1 μL, 202 μmol), HBTU (28.1 mg, 74.1 μmol) and the modified D-lysine intermediate **S32** (54 mg, 67.3 μmol) were added and the reaction mixture was stirred at room temperature for 2 h. After complete reaction, 40 mL ethyl acetate was added into the reaction flask, followed by washing with water (40 mL, 2 x) and brine (40 mL). The organic layer was dried over $MgSO_4$ and then evaporated under reduced pressure. The residue was purified by column chromatography to yield the intermediate as a red solid (66%). LC-MS ($ESI^+$): calculated for $C_{79}H_{125}N_{10}O_{17}$ $[M]^+$ m/z: 1485.92; found: 1485.75; $^1H$ NMR (400 MHz, MeOD-$d_4$) δ 7.86 – 7.68 (m, 3H), 7.54 – 7.46 (m, 1H), 7.32 (dd, *J* = 9.5, 3.1 Hz, 2H), 7.11 – 7.03 (m, 2H), 6.98 (t, *J* = 3.5 Hz, 2H), 4.04 – 3.84 (m, 3H), 3.76 – 3.65 (m, 8H), 3.62 (dd, *J* = 5.8, 2.8 Hz, 1H), 3.58 – 3.49 (m, 4H), 3.47 – 3.37 (m, 4H), 3.37 – 3.33 (m,2H), 3.29 – 3.23 (m, 2H), 3.23 – 3.08 (m, 4H), 3.02 (t, *J* = 6.7 Hz, 3H), 2.45 (t, *J* = 7.2 Hz, 1H), 1.81 (t, *J* = 7.1 Hz, 1H), 1.76 – 1.64 (m, 3H), 1.63 – 1.55 (m, 3H), 1.55 – 1.35 (m, 51H), 1.33 (dd, *J* = 12.5, 5.3 Hz, 18H).

**Synthesis of compound S34**

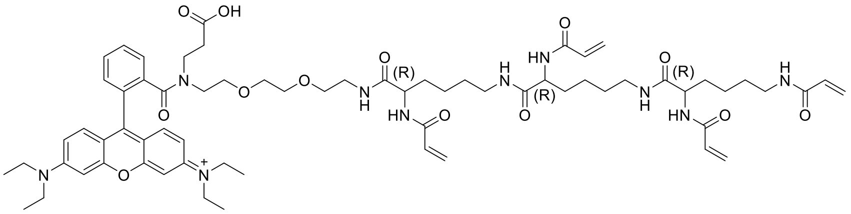

To a solution of the intermediate **S33** in DCM (0.2 mL), TFA (0.2 mL) was added and the reaction mixture was stirred overnight at room temperature. After complete reaction, all solvents were evaporated to yield the intermediate as a red solid, which was used without further purification.

To a solution of the intermediate (78 mg, 52.5 μmol) in MeOH: THF (v:v = 0.4 mL: 0.4 mL), a solution of $Na_2CO_3$ (56 mg, 525 μmol) in water (0.4 mL) was added under the protection of $N_2$, followed by cooling the reaction flask to 0 ℃ with an ice bath. A solution of acryloyl chloride (30 μL, 368 μmol) in dry dioxane (0.3 mL) was added dropwise to the reaction flask, and the reaction mixture was then allowed to come to room temperature over 20 min. After complete reaction, all solvents were evaporated and the residue was purified by column chromatography to yield the product as a red solid (43%). LC-MS (ESI$^+$): calculated for $C_{67}H_{93}N_{10}O_{13}$ [M]$^+$ m/z: 1245.69; found: 1245.45; $^1$H NMR (400 MHz, MeOD-$d_4$) δ 7.94 – 7.58 (m, 3H), 7.49 (dd, $J$ = 7.8, 4.7 Hz, 1H), 7.35 – 7.28 (m, 2H), 7.10 – 7.04 (m, 2H), 6.99 – 6.93 (m, 2H), 6.45 – 6.10 (m, 8H), 5.74 – 5.55 (m, 4H), 4.40 – 4.28 (m, 3H), 3.78 – 3.65 (m, 8H), 3.62 – 3.58 (m, 1H), 3.58 – 3.49 (m, 3H), 3.46 – 3.36 (m, 6H), 3.28 – 3.22 (m, 4H), 3.21 – 3.12 (m, 5H), 3.04 (t, $J$ = 5.3 Hz, 1H), 2.35 (t, $J$ = 7.7 Hz, 1H), 1.87 – 1.74 (m, 4H), 1.72 – 1.63 (m, 3H), 1.58 – 1.47 (m, 6H), 1.45 – 1.34 (m, 6H), 1.31 (t, $J$ = 7.1 Hz, 12H); $^{13}$C NMR (101 MHz, MeOD-$d_4$) δ 173.03, 172.85, 169.62, 168.57, 166.71, 166.55, 166.48, 136.44, 136.28, 132.19, 130.73, 130.47, 130.10, 129.84, 129.37, 129.24, 128.82, 127.23, 125.81, 125.14, 113.95, 113.79, 96.14, 95.86, 70.25, 69.87, 69.16, 67.88, 67.47, 53.66, 43.38, 40.94, 39.09, 38.97, 38.66, 38.61, 35.52, 31.50, 29.36, 28.61, 28.53, 22.95, 22.80, 11.51.

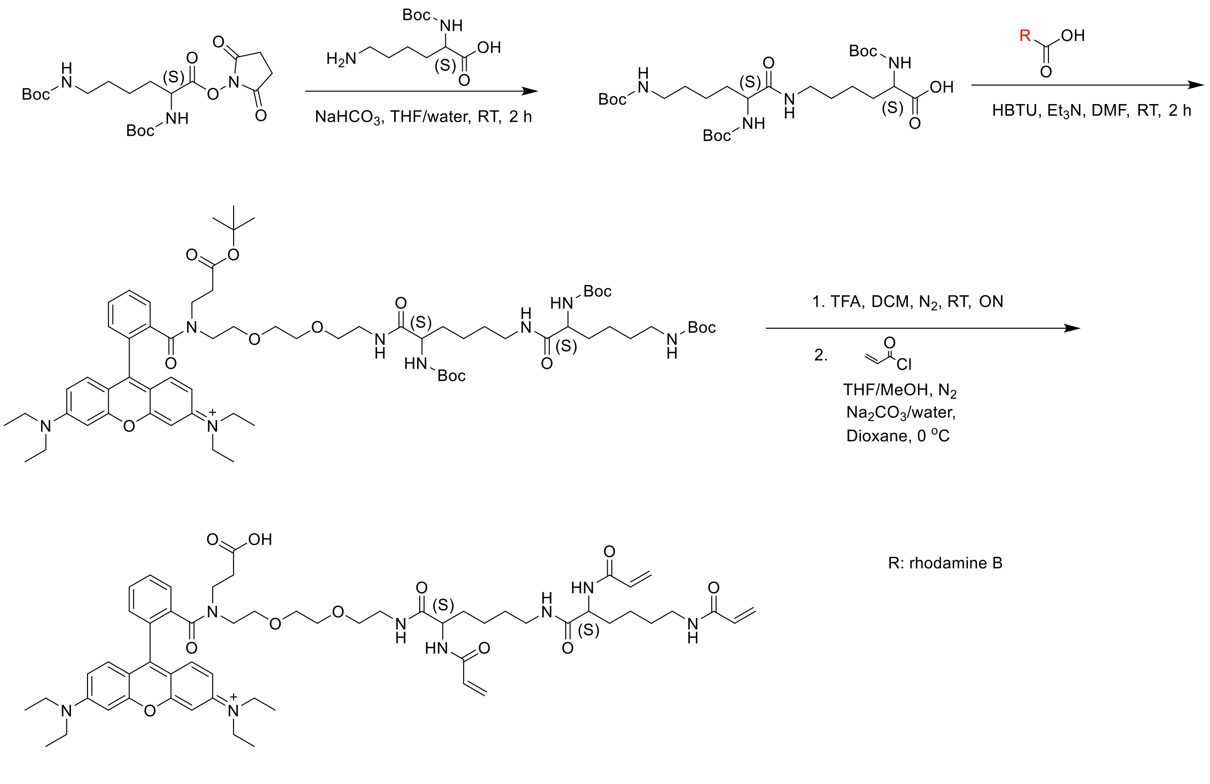

**Synthesis of compound S35**

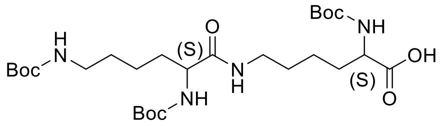

To a solution of Boc-Lys(Boc)-OSu (781 mg, 1.76 mmol) in $H_2O$/THF (v:v = 3 mL: 30 mL), $NaHCO_3$ (163 mg, 1.94 mmol) and Boc-Lys-OH (477 mg, 1.94 mmol) were added and the reaction mixture was stirred at room temperature for 2 h. After complete reaction, THF was evaporated, followed by the addition of 70 mL water. And the aqueous solution was adjusted to pH 5 by using citric acid, followed by extraction with DCM (100 mL, 3 x). The organic layer was then washed with brine (100 mL, 1 x), dried over $MgSO_4$ and evaporated under reduced pressure. The residue was purified by column chromatography to yield the intermediate as a white solid (65%). LC-MS ($ESI^+$): calculated for $C_{27}H_{49}N_4O_9$ $[M-H]^-$ m/z: 573.35; found: 573.05; $^1H$ NMR (400 MHz, DMSO-*d*4) δ 12.36 (brs, 1H), 7.74 (t, *J* = 5.5 Hz, 1H), 6.99 (d, *J* = 7.9 Hz, 1H), 6.78 – 6.61 (m, 2H), 3.86 – 3.75 (m, 2H), 3.07 – 2.94 (m, 2H), 2.90 – 2.82 (m, 2H), 1.67 – 1.43 (m, 4H), 1.38 – 1.19 (m, 35H).

**Synthesis of compound S36**

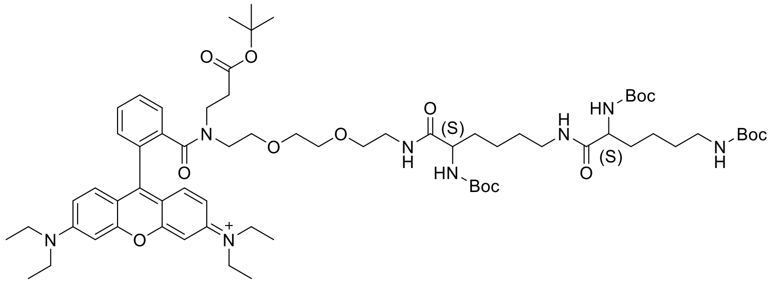

To a solution of the modified rhodamine B (71 mg, 87 μmol, synthesized from the literature[2]) in DMF (1 mL), trimethylamine (36 μL, 261 μmol), HBTU (34 mg, 90 μmol) and the modified L-lysine intermediate **S35** (47 mg, 82 μmol) were added and the reaction mixture was stirred at room temperature for 2 h. After complete reaction, 50 mL ethyl acetate was added into the reaction flask, followed by washing with water (50 mL, 2 x) and brine (40 mL). The organic layer was dried over $MgSO_4$ and then evaporated under reduced pressure. The residue was purified by column chromatography to yield the intermediate as a red solid (78%). LC-MS ($ESI^+$): calculated for $C_{68}H_{105}N_8O_{14}$ $[M]^+$ m/z: 1257.77; found: 1257.60; $^1H$ NMR (400 MHz, MeOD-*d*4) δ 7.86 – 7.66 (m, 3H), 7.54 – 7.47 (m, 1H), 7.31 (dt, *J* = 7.8, 3.7 Hz, 2H), 7.11 – 7.03 (m, 2H), 6.98 (dd, *J* = 5.7, 1.9 Hz, 2H), 4.08 – 3.78 (m, 2H), 3.76 – 3.66 (m, 8H), 3.62 (dd, *J* = 5.9, 2.9 Hz, 1H), 3.59 – 3.51 (m, 4H), 3.49 – 3.39 (m, 4H), 3.39 – 3.32 (m, 3H), 3.29 – 3.23 (d, *J* = 3.8 Hz, 2H), 3.23 – 3.05 (m, 2H), 3.02 (t, *J* = 6.7 Hz, 2H), 2.45 (t, *J* = 7.2 Hz, 1H), 1.81 (t, *J* = 7.1 Hz, 1H), 1.77 – 1.63 (m, 2H), 1.63 – 1.54 (m, 2H), 1.54 – 1.37 (m, 34H), 1.38 – 1.02 (m, 22H).

#### Synthesis of compound S37

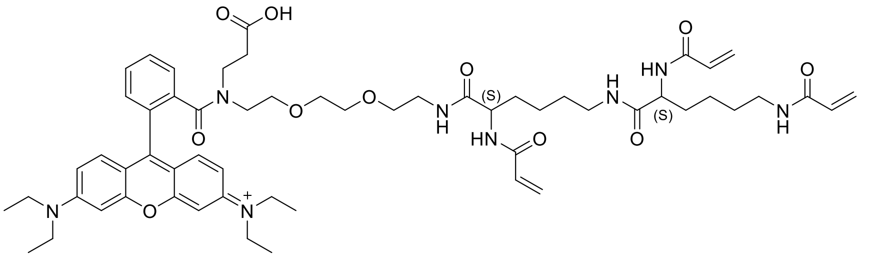

To a solution of the intermediate **S36** in DCM (0.3 mL), TFA (0.3 mL) was added and the reaction mixture was stirred overnight at room temperature. After complete reaction, all solvents were evaporated to yield the intermediate as a red solid, which was used without further purification.

To a solution of the intermediate (80 mg, 64 μmol) in MeOH: THF (v:v = 0.35 mL: 0.35 mL), a solution of $Na_2CO_3$ (51 mg, 0.482 mmol) in water (0.35 mL) was added under the protection of $N_2$, followed by cooling the reaction flask to 0 ℃ with an ice bath. A solution of acryloyl chloride (23.4 μL, 289 μmol) in dry dioxane (0.25 mL) was added dropwise to the reaction flask, and the reaction mixture was then allowed to come to room temperature over 20 min. After complete reaction, all solvents were evaporated and the residue was purified by column chromatography to yield the product as a red solid (48%). LC-MS ($ESI^+$): calculated for $C_{58}H_{79}N_8O_{11}$ $[M]^+$ m/z: 1063.59; found: 1063.35; $^1H$ NMR (400 MHz, MeOD-*d*$_4$) δ 7.86 – 7.67 (m, 3H), 7.55 – 7.47 (m, 1H), 7.32 (d, *J* = 9.5 Hz, 2H), 7.07 (ddd, *J* = 9.5, 5.2, 2.3 Hz, 2H), 7.01 – 6.92 (m, 2H), 6.54 – 6.10 (m, 6H), 5.75 – 5.58 (m, 3H), 4.43 – 4.27 (m, 2H), 3.76 – 3.65 (m, 8H), 3.66 – 3.57 (m, 2H), 3.57 – 3.51 (m, 3H), 3.48 (t, *J* = 4.8 Hz, 1H), 3.46 – 3.33 (m, 6H), 3.30 – 3.19 (m, 4H), 3.19 – 3.02 (m, 2H), 2.46 (t, *J* = 7.3 Hz, 1H), 1.89 (t, *J* = 6.6 Hz, 1H), 1.85 – 1.74 (m, 2H), 1.73 – 1.62 (m, 2H), 1.60 – 1.47 (m, 4H), 1.46 – 1.35 (m, 4H), 1.31 (t, *J* = 7.0 Hz, 12H); $^{13}C$ NMR (101 MHz, MeOD-*d*$_4$) δ 173.42, 172.97, 172.82, 170.48, 169.76, 166.72, 166.57, 166.50, 163.82, 157.87, 155.82, 155.27, 155.03, 136.36, 136.12, 132.10, 131.87, 130.69, 130.41, 130.09, 129.96, 129.88, 129.76, 129.35, 128.96, 127.38, 125.90, 125.20, 115.36, 113.89, 113.83, 113.32, 113.17, 96.18, 95.94, 70.20, 70.01, 69.88, 69.15, 68.13, 67.51, 53.66, 53.49, 49.86, 45.97, 45.49, 43.56, 40.32, 39.08, 38.94, 38.64, 38.60, 31.48, 31.43, 30.52, 28.59, 28.52, 22.91, 22.75, 11.45.

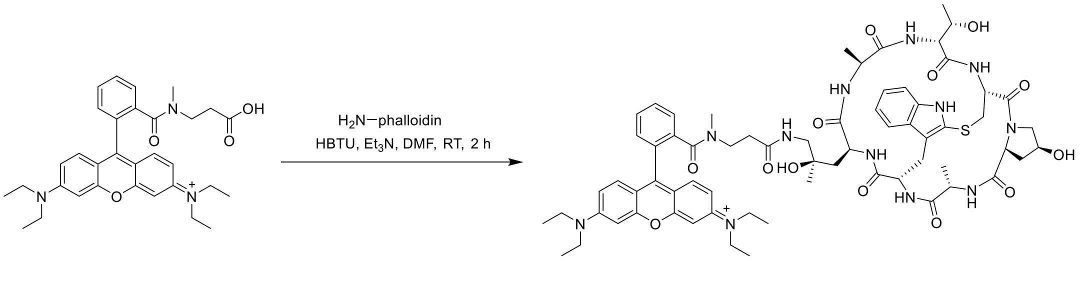

#### Synthesis of compound 1

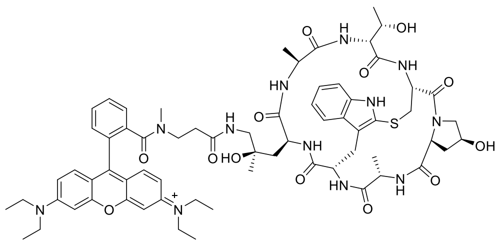

To a solution of the intermediate **S2** (5.8 μL, 34 mM in DMSO), triethylamine (4.86 μL, 72 mM in DMSO), HBTU (2.86 μL, 70 mM in DMSO) and phalloidin amine (tosylate) (10 μL, 10 mM solution in DMF) were added. Upon indication of complete reaction, the crude mixture was purified using preparative HPLC, and the desired products was used as such. Preparative HPLC: Shim-pack GIST C18 2 μm column, Eluent A: 0.1% HCOOH in Milli-Q water, Eluent B: MeOH, gradient elution (20%-100% B from 0-30 min; 100% B from 30-35 min), pump flow: 2.00 mL/min, $T_R$ = 19.73 min. Red solid; Yield: 20%; LC-MS (ESI$^+$): calculated for $C_{67}H_{85}N_{12}O_{13}S$ $[M]^+$ m/z: 1297.61; found: 1297.40.
Other dye-phalloidin analogues were synthesized and purified based on the same method, unless stated.

#### Synthesis of compound 2

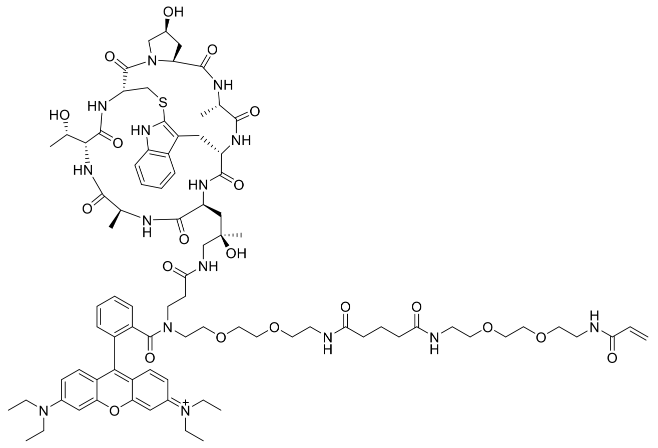

Preparative HPLC: Shim-pack GIST C18 2 μm column, Eluent A: 0.1% HCOOH in Milli-Q water, Eluent B: MeOH, gradient elution (20%-100% B from 0-30 min; 100% B from 30-35 min), pump flow: 2.00 mL/min, $T_R$ = 19.26 min. Red solid; Yield: 15%; LC-MS (ESI$^+$): calculated for $C_{86}H_{118}N_{15}O_{20}S$ $[M]^+$ m/z: 1712.84; found: 1712.95.

**Synthesis of compound 3**

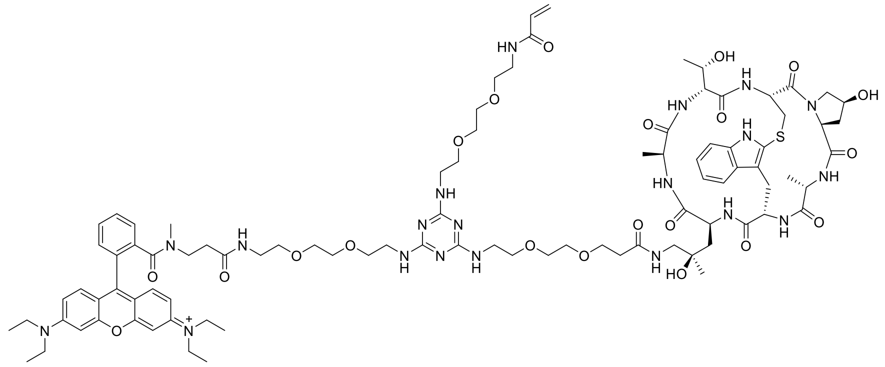

Preparative HPLC: Shim-pack GIST C18 2 μm column, Eluent A: 0.1% HCOOH in Milli-Q water, Eluent B: MeOH, gradient elution (20%-100% B from 0-30 min; 100% B from 30-35 min), pump flow: 2.00 mL/min, $T_R$ = 17.87 min. Red solid; Yield: 8%; LC-MS (ESI$^+$): calculated for $C_{92}H_{129}N_{20}O_{21}S$ $[M]^+$ m/z: 1881.94; found: 1882.05.

**Synthesis of compound 4**

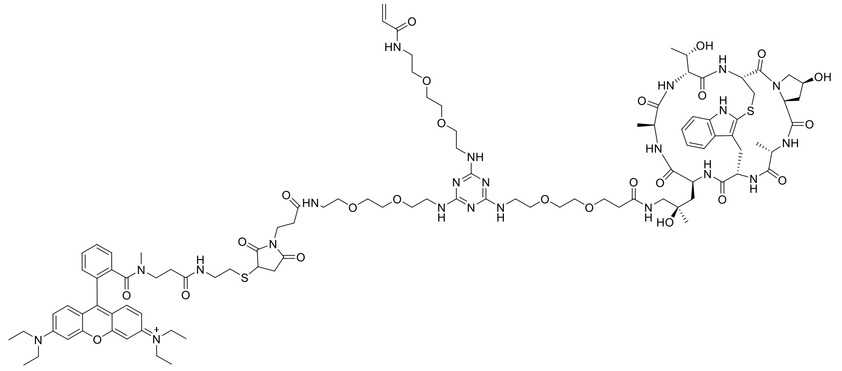

To a solution of **S12** (3.2 μL, 48 mM in DMSO), triethylamine (4.3 μL, 72 mM in DMSO), HBTU (1.8 μL, 105 mM in DMSO) and phalloidin amine (tosylate) (10 μL, 10 mM solution in DMF) were added and the reaction mixture was stirred at room temperature for 1.5 h. Upon indication of complete reaction, the crude mixture was used in the next step without further purification.

To a solution of **S3** (1.47 μL, 34 mM in DMSO), tris(2-carboxyethyl)phosphine (TCEP, 4.5 μL, 16.7 mM in DMSO) was added and the reaction mixture was stirred at room temperature for 0.5 h. Upon indication of complete reaction, the crude mixture synthesized above was added, followed by adding triethylamine (3.5 μL, 72 mM in DMSO). The reaction mixture was then stirred for another 0.5 h. After complete reaction, the crude mixture was purified using preparative HPLC. Preparative HPLC: Shim-pack GIST C18 2 μm column, Eluent A: 0.1% HCOOH in Milli-Q water, Eluent B: MeOH, gradient elution (20%-100% B from 0-30 min; 100% B from 30-35 min), pump flow: 2.00

mL/min, $T_R$ = 18.00 min. Red solid; Yield: 9%; LC-MS (ESI$^+$): calculated for $C_{101}H_{142}N_{22}O_{24}S_2$ $[M+H]^{2+}$ m/z: 2111.00; found: 1055.60.

**Synthesis of compound 5**

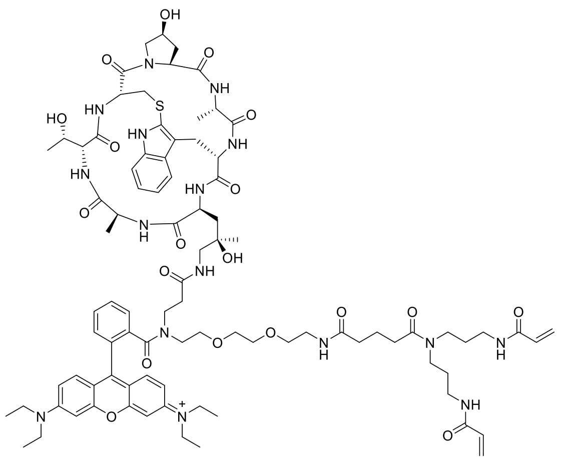

Preparative HPLC: Shim-pack GIST C18 2 μm column, Eluent A: 0.1% HCOOH in Milli-Q water, Eluent B: MeOH, gradient elution (20%-100% B from 0-30 min; 100% B from 30-35 min), pump flow: 2.00 mL/min, $T_R$ = 19.40 min. Red solid; Yield: 16%; LC-MS (ESI$^+$): calculated for $C_{89}H_{121}N_{16}O_{19}S$ $[M]^+$ m/z: 1749.87; found: 1749.95.

**Synthesis of compound 6**

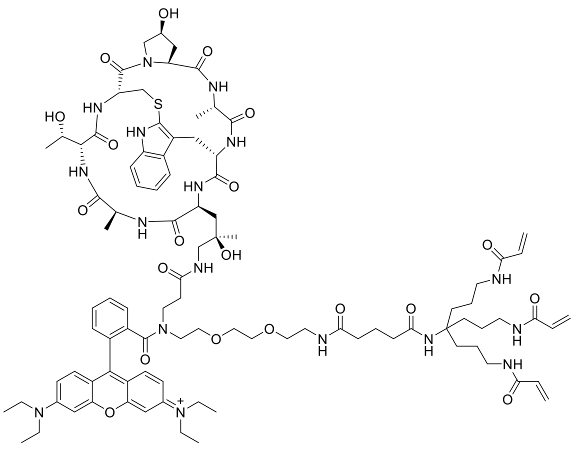

Preparative HPLC: Shim-pack GIST C18 2 μm column, Eluent A: 0.1% HCOOH in Milli-Q water, Eluent B: MeOH, gradient elution (20%-100% B from 0-30 min; 100% B from 30-35 min), pump flow: 2.00 mL/min, $T_R$ = 19.68 min. Red solid; Yield: 20%; LC-MS (ESI$^+$): calculated for $C_{96}H_{133}N_{17}O_{20}S$ $[M+H]^{2+}$ m/z: 1875.96; found: 938.40.

**Synthesis of compound S38**

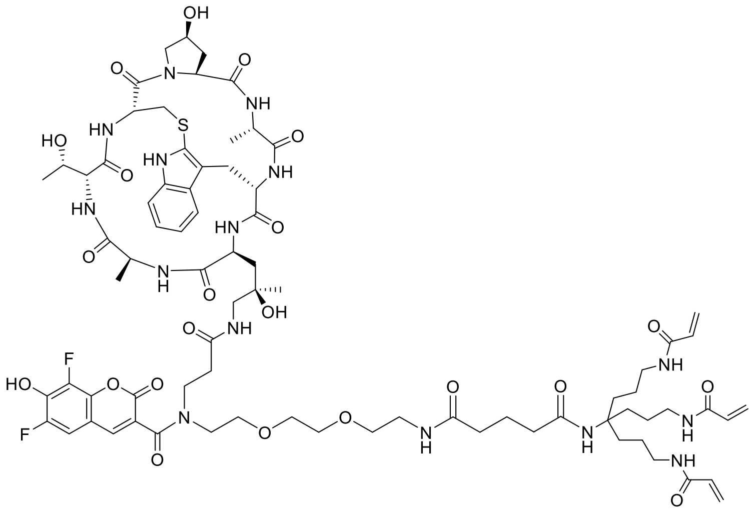

Preparative HPLC: Shim-pack GIST C18 2 μm column, Eluent A: 0.1% HCOOH in Milli-Q water, Eluent B: MeOH, gradient elution (20%-100% B from 0-30 min; 100% B from 30-35 min), pump flow: 2.00 mL/min, $T_R$ = 15.34 min. Green solid; Yield: 14%; LC-MS ($ESI^+$): calculated for $C_{78}H_{106}F_2N_{15}O_{22}$ $[M+H]^+$ m/z: 1674.73; found: 1674.85.

**Synthesis of compound S39**

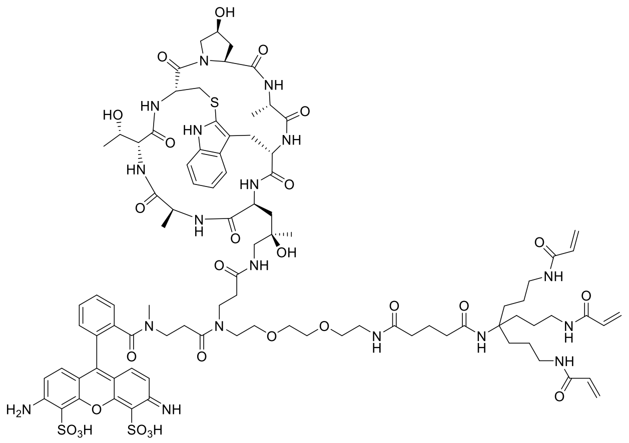

Preparative HPLC: Shim-pack GIST C18 2 μm column, Eluent A: 0.1% HCOOH in Milli-Q water, Eluent B: MeOH, gradient elution (20%-100% B from 0-30 min; 100% B from 30-35 min), pump flow: 2.00 mL/min, $T_R$ = 13.03 min. Orange solid; Yield: 10%; LC-MS ($ESI^+$): calculated for $C_{92}H_{124}N_{18}O_{27}S_3$ $[M+2H]^{2+}$ m/z: 2008.80; found: 1004.55.

#### Synthesis of compound S40

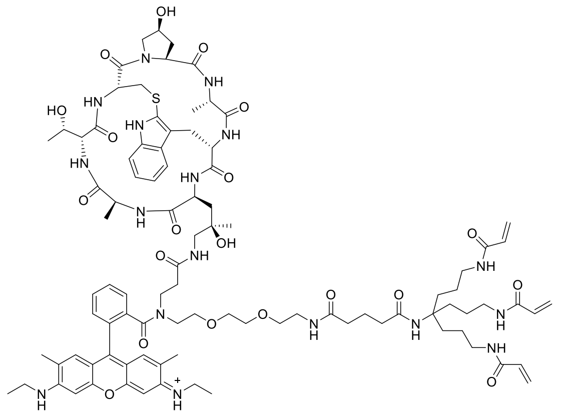

The starting material was synthesized from the literature[2]. Preparative HPLC: Shim-pack GIST C18 2 μm column, Eluent A: 0.1% HCOOH in Milli-Q water, Eluent B: MeOH, gradient elution (20%-100% B from 0-30 min; 100% B from 30-35 min), pump flow: 2.00 mL/min, $T_R$ = 18.57 min. Orange solid; Yield: 14%; LC-MS ($ESI^+$): calculated for $C_{94}H_{128}N_{17}O_{20}S$ $[M]^+$ m/z: 1846.92; found: 1847.15.

#### Synthesis of compound 12

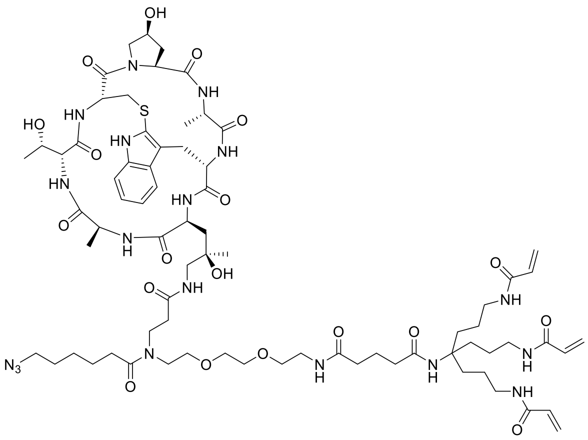

The crude product was used without further purification. LC-MS ($ESI^+$): calculated for $C_{74}H_{113}N_{18}O_{19}S$ $[M+H]^+$ m/z: 1589.81; found: 1589.60.

**Synthesis of compound S41**

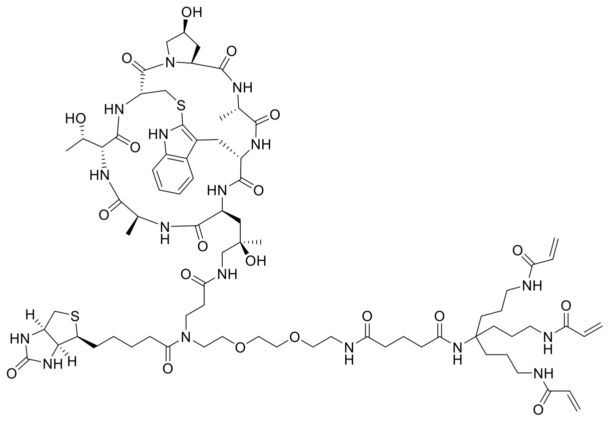

The crude product was used without further purification. LC-MS (ESI+): calculated for $C_{78}H_{120}N_{17}O_{20}S_2$ $[M+3H]^{3+}$ m/z: 1678.83; found: 559.60.

**Synthesis of compound 9**

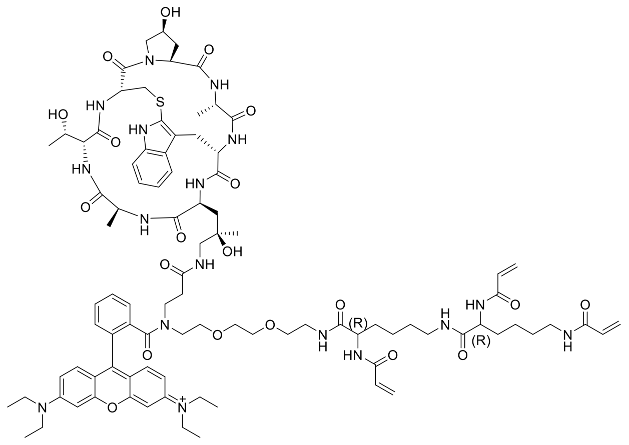

Preparative HPLC: Shim-pack GIST C18 2 μm column, Eluent A: 0.1% HCOOH in Milli-Q water, Eluent B: MeOH, gradient elution (20%-100% B from 0-30 min; 100% B from 30-35 min), pump flow: 2.00 mL/min, $T_R$ = 19.55 min. Red solid; Yield: 28%; LC-MS (ESI+): calculated for $C_{93}H_{127}N_{17}O_{20}S$ $[M+H]^{2+}$ m/z: 916.96; found: 917.10.

#### Synthesis of compound 10

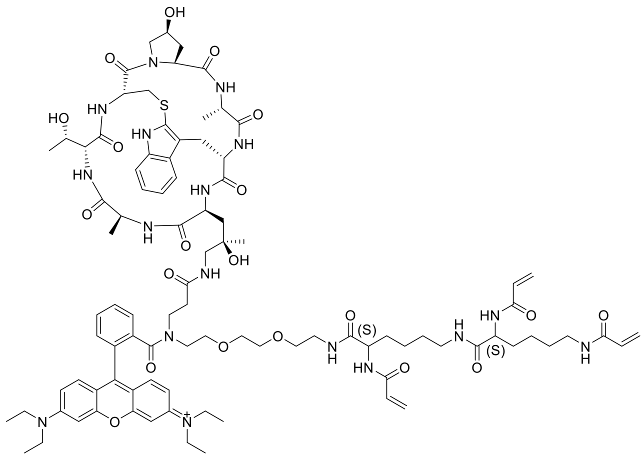

Preparative HPLC: Shim-pack GIST C18 2 μm column, Eluent A: 0.1% HCOOH in Milli-Q water, Eluent B: MeOH, gradient elution (20%-100% B from 0-30 min; 100% B from 30-35 min), pump flow: 2.00 mL/min, $T_R$ = 19.68 min. Red solid; Yield: 28%; LC-MS (ESI$^+$): calculated for $C_{93}H_{127}N_{17}O_{20}S$ $[M+H]^{2+}$ m/z: 916.96; found: 917.15.

#### Synthesis of compound 11

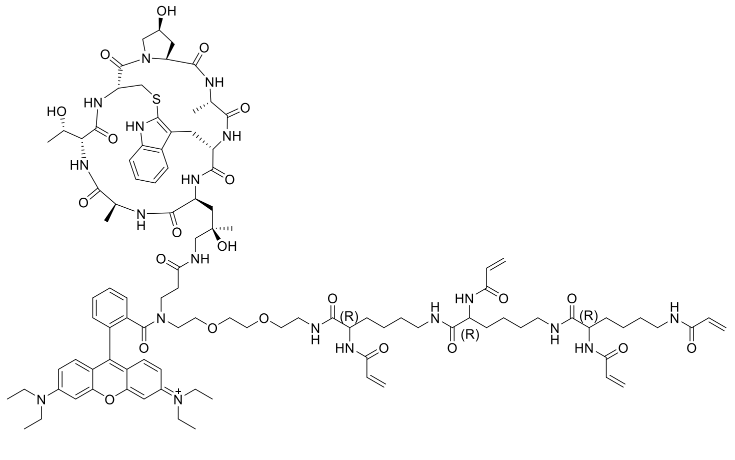

Preparative HPLC: Shim-pack GIST C18 2 μm column, Eluent A: 0.1% HCOOH in Milli-Q water, Eluent B: MeOH, gradient elution (20%-100% B from 0-30 min; 100% B from 30-35 min), pump flow: 2.00 mL/min, $T_R$ = 19.66 min. Red solid; Yield: 25%; LC-MS (ESI$^+$): calculated for $C_{102}H_{141}N_{19}O_{22}S$ $[M+H]^{2+}$ m/z: 1008.01; found: 1008.25.

#### Supplementary Figures

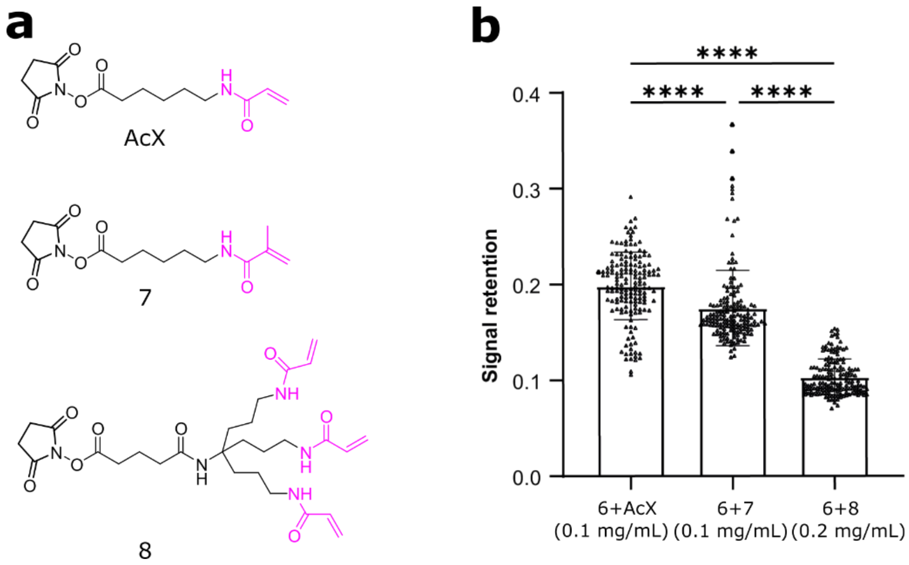

**Figure S1. Quantification of the influence of different anchoring agents on post-digestion signal retention of actin filaments**. (a) Molecular structures of anchoring agents. (b) Comparison of post-digestion signal retention of actin filaments stained with compound **6** and different alternative anchoring agents. Bars represent the mean value and error bars represent the standard deviation. Statistical significance is assessed by one-way ANOVA test. **** $p < 0.0001$. From left to right, mean values are 0.199 ± 0.028, 0.175 ± 0.039, 0.104 ± 0.019 (n= 162 from three independent samples), respectively.

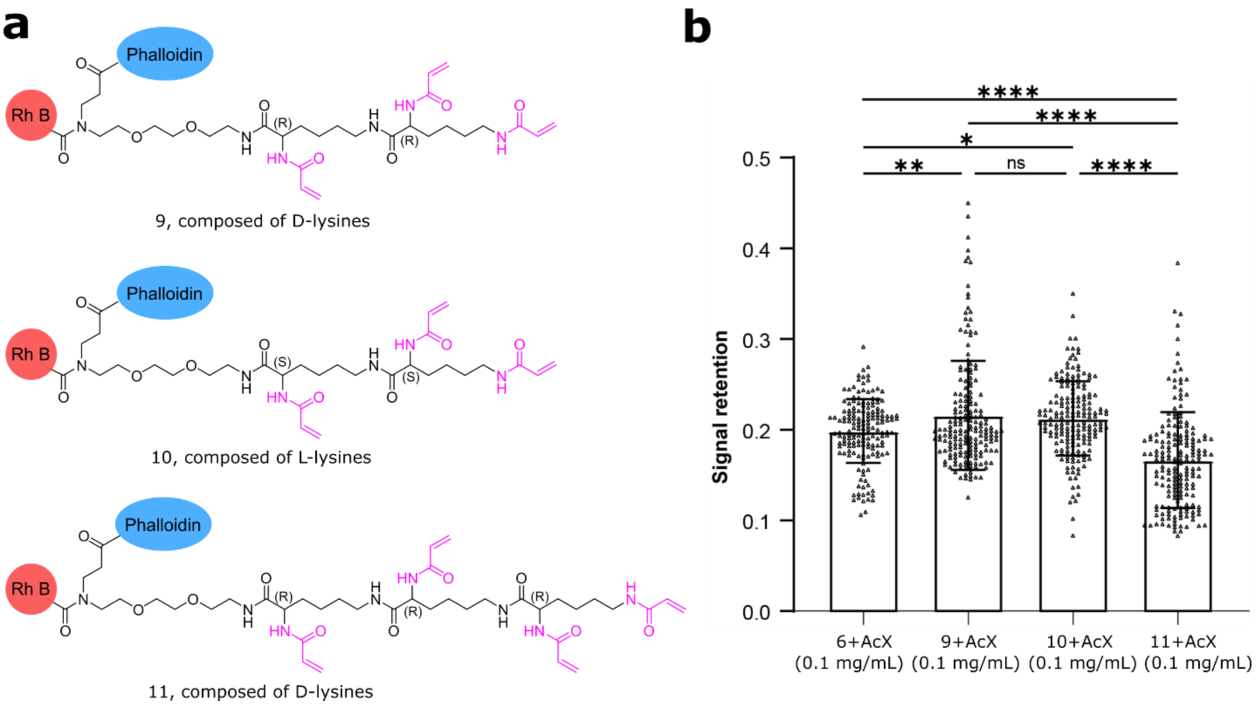

**Figure S2. Quantification of the post-digestion signal retention of actin filaments stained with TRITON linkers containing linear multiple acryloyl units.** (a) Structures of branched multiple acryloyl-containing TRITON linkers. (b) Comparison of post-digestion signal retention of actin filaments stained with different TRITON linkers. Bars represent the mean value and error bars represent the standard deviation. Statistical significance is assessed by one-way ANOVA test. * $p < 0.05$; ** $p < 0.01$; **** $p < 0.0001$; ns (non-significant)= 0.9117. From left to right, mean values are 0.199 ± 0.035 (n= 162 from three independent samples), 0.216 ± 0.060 (n= 184), 0.212 ± 0.041 (n= 184) and 0.167 ± 0.053 (n= 184), respectively.

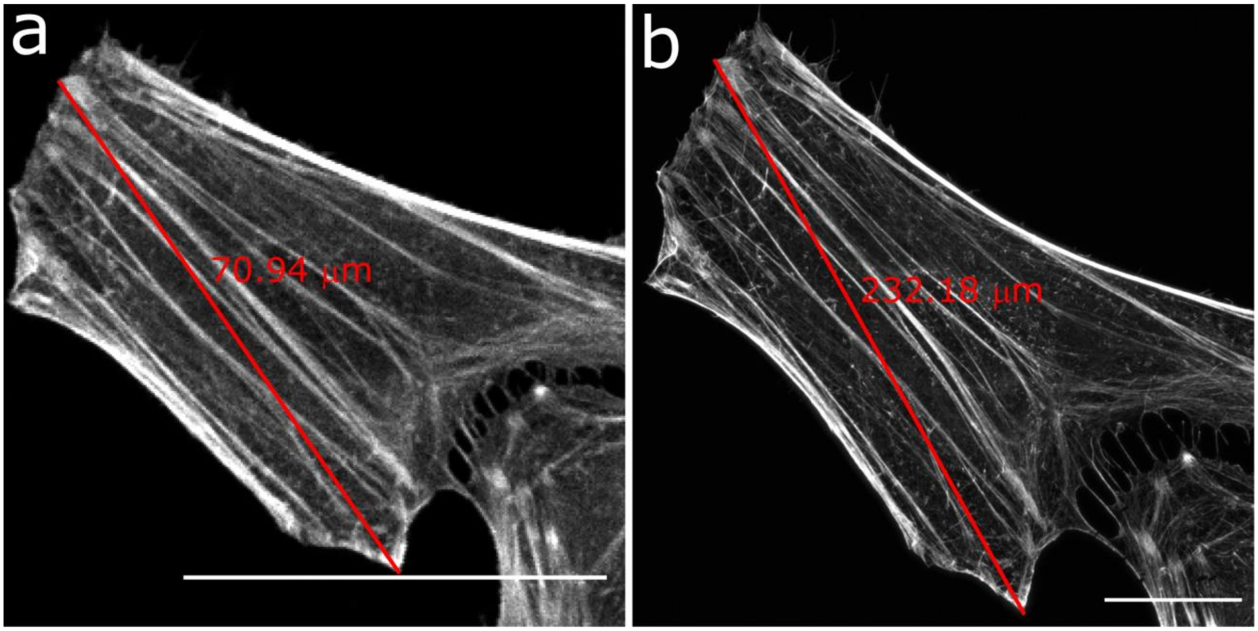

**Figure S3. Expansion factor calculated after expansion.** (a) Pre-expansion image of actin filaments; (b) Post-expansion image of actin filaments in the same cells. Scale bars, 50 μm (a-b).

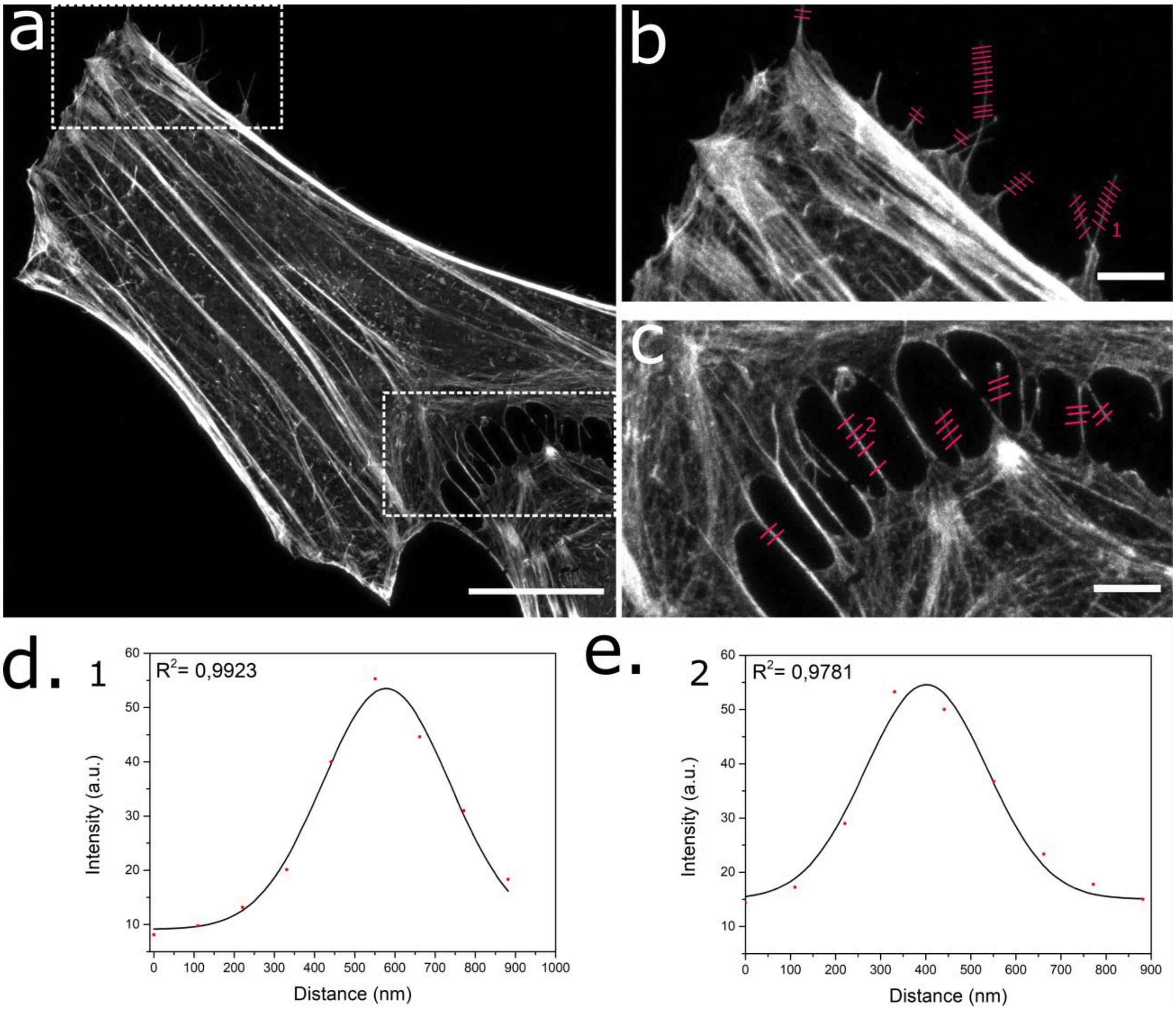

**Figure S4. Quantification of resolution achieved in 4x ExM.** (a) ExM imaging of actin filaments stained with our TRITON. (b-c) FWHM line profiles of actin filaments used for resolution determination were shown in red (n= 53). (d-e) Two representative cross-sectional intensity profiles of actin filament (dots) with Gaussian fitting (solid line), one line in b, another line in c. Scale bars, 50 μm (a), 10 μm (b, c).

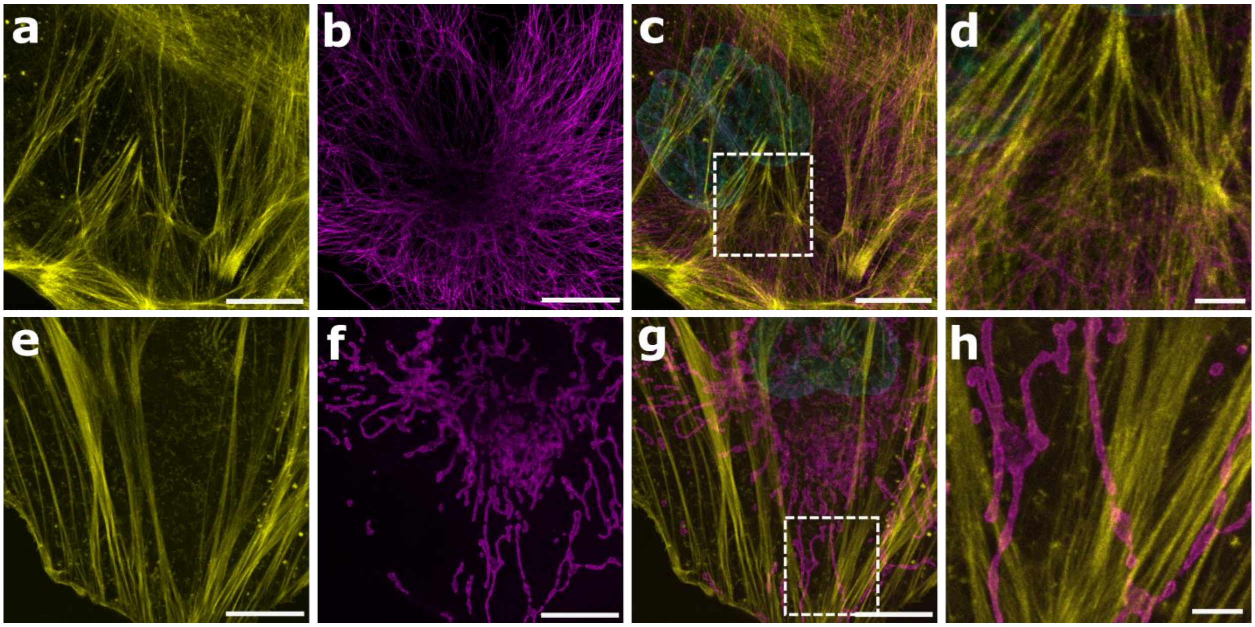

**Figure S5.** Super-resolution imaging of actin filaments and microtubules or mitochondria in Hela cells. (a-d) Simultaneous staining of actin filaments and microtubules in 4x ExM. Actin filaments (yellow) were stained with TRITON (rhodamine 6G, acrylates, and phalloidin) and microtubules (magenta) were stained with TRITON (ATTO 647N, acrylates and TFP)-labeled secondary antibodies. (a) Post-expansion image of actin filaments. (b) Post-expansion image of microtubules. (c) Merged image of actin filaments and microtubules, in which nuclei were stained with 4′,6-diamidino-2-phenylindole (DAPI). (d) Zoom-in of the boxed region in panel c. (e-h) Simultaneous staining of actin filaments and mitochondria in 4x ExM. Actin filaments (yellow) were stained with TRITON (rhodamine 6G, acrylates, and phalloidin) and mitochondria (magenta) were stained with TRITON (ATTO 647N, acrylates and TFP)-labeled secondary antibodies. (e) Post-expansion image of actin filaments. (f) Post-expansion image of mitochondria. (g) Merged image of actin filaments and mitochondria, in which nuclei were stained with DAPI. (h) Zoom-in of the boxed region in panel g. Scale bars, 50 μm (a-c, e-g) and 10 μm (d, h).

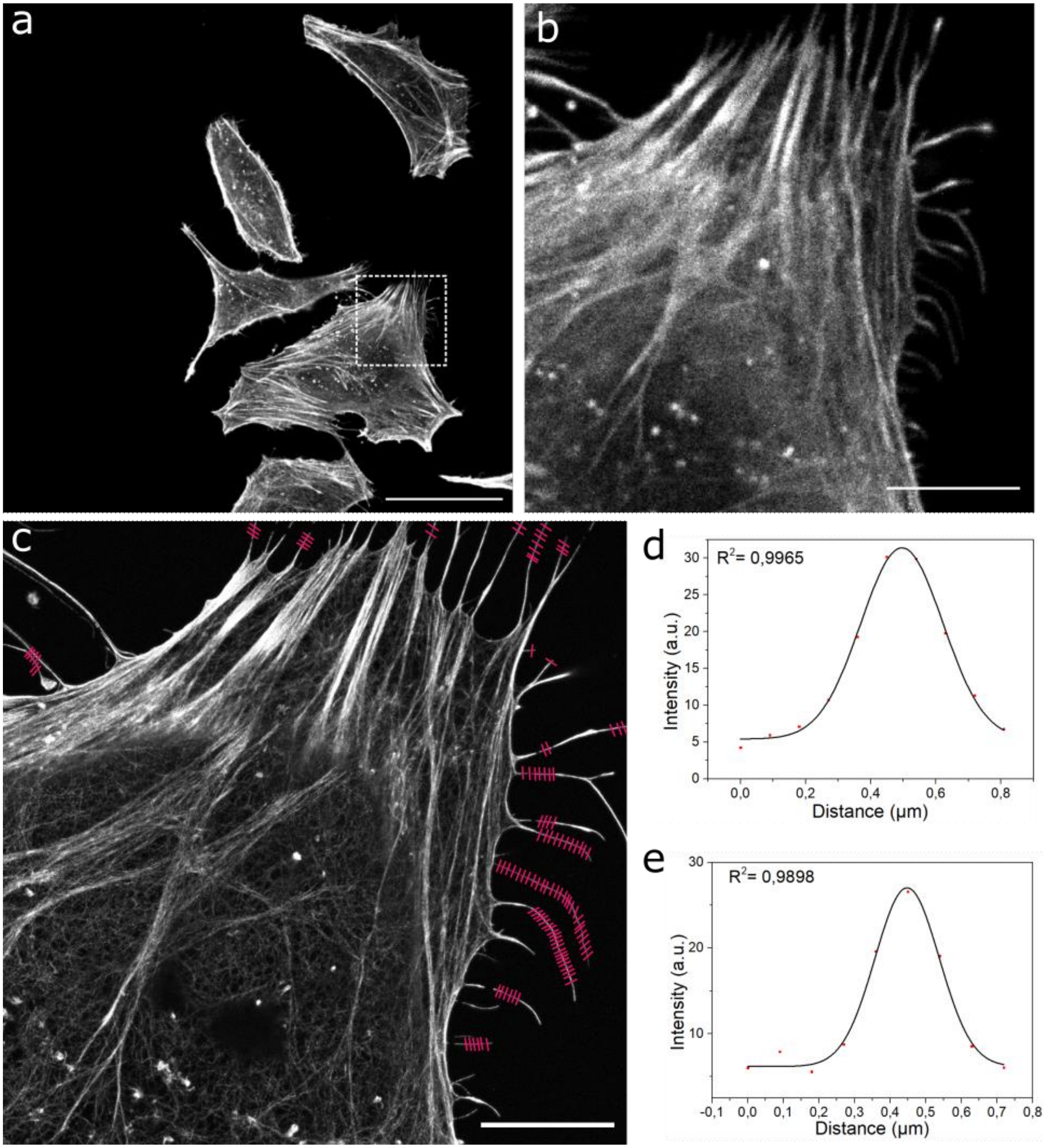

**Figure S6. Quantification of resolution achieved in TREx.** (a-b) Pre-expansion image of actin filaments. (c) FWHM line profiles of expanded actin filaments used for resolution determination were shown in red. (d-e) Two representative cross-sectional intensity profiles of actin filament (dots) with Gaussian fitting (solid line). Scale bars, 50 μm (a, c), 10 μm (b).

### Copies of $^1H$ and $^{13}C$ NMR spectra

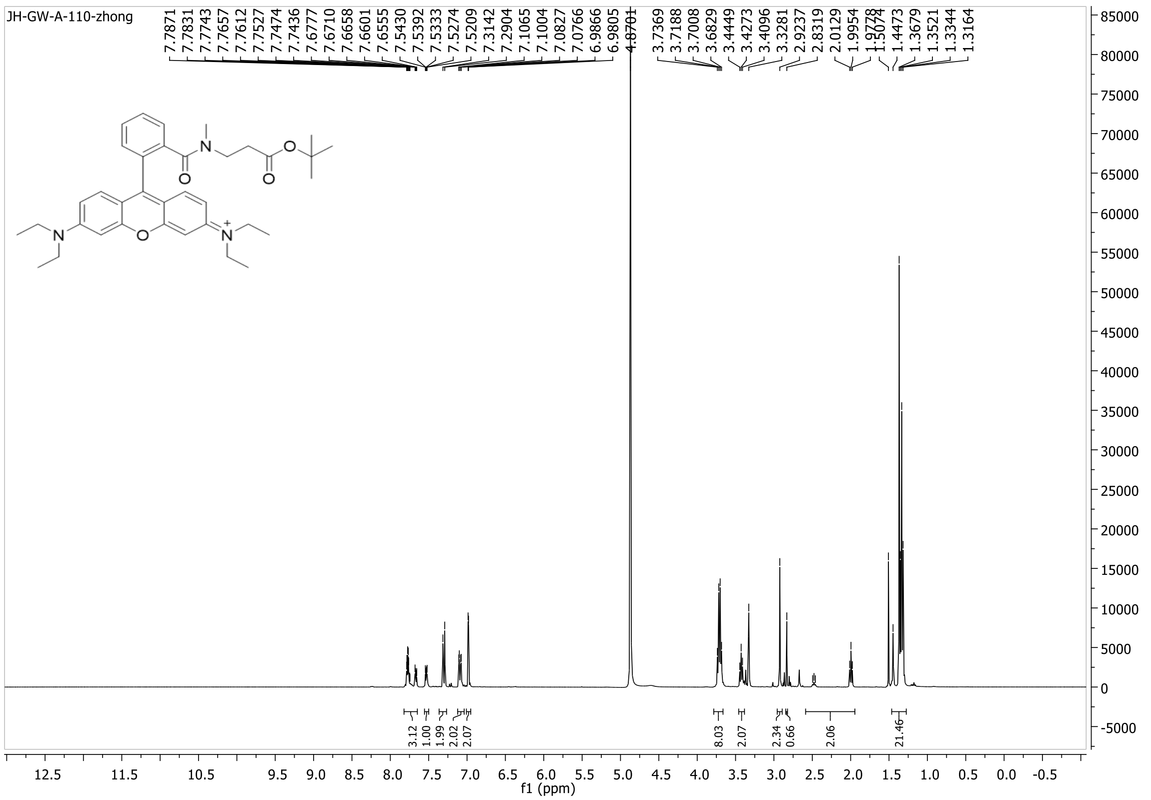

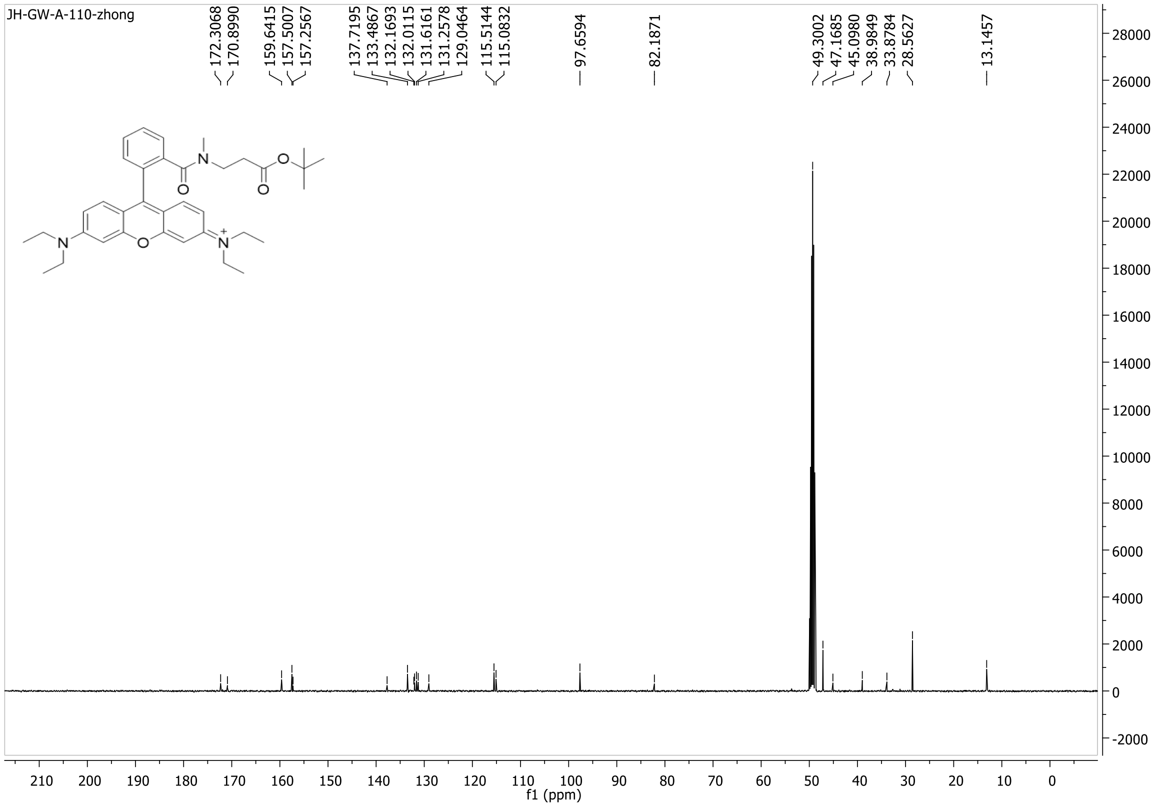

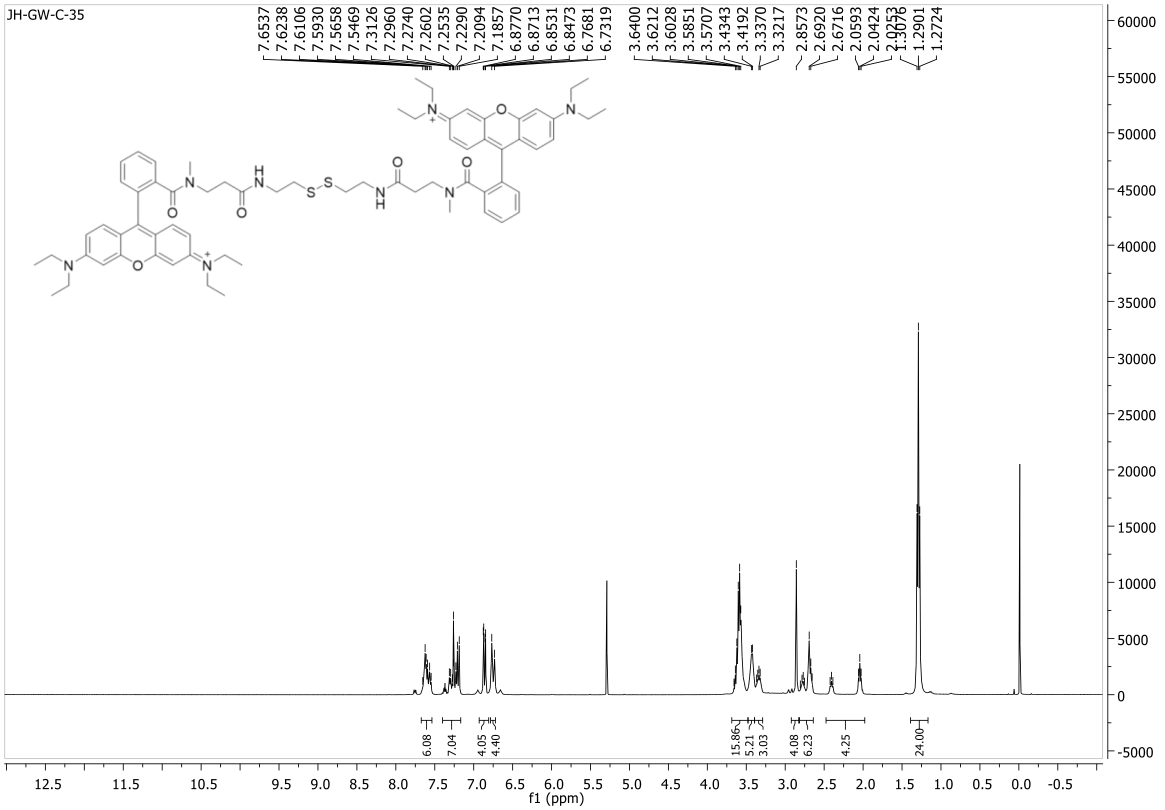
JH-GW-C-35
f1 (ppm)

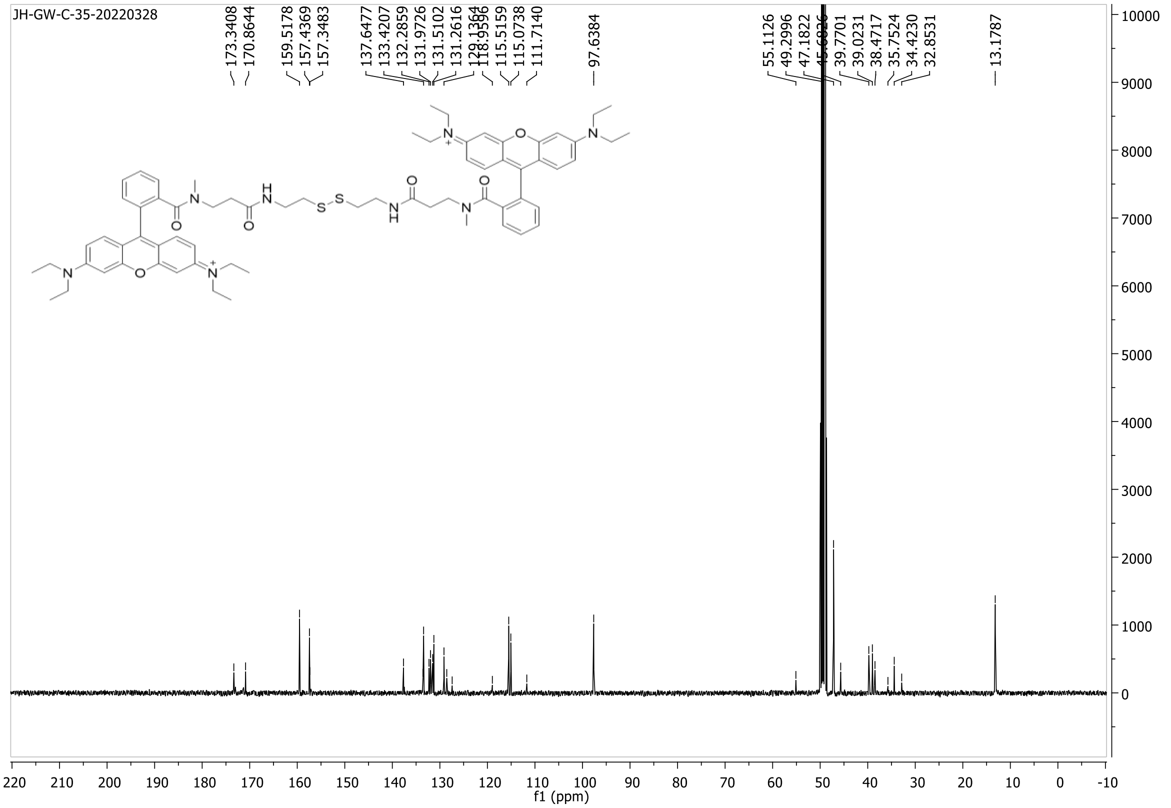
JH-GW-C-35-20220328
f1 (ppm)

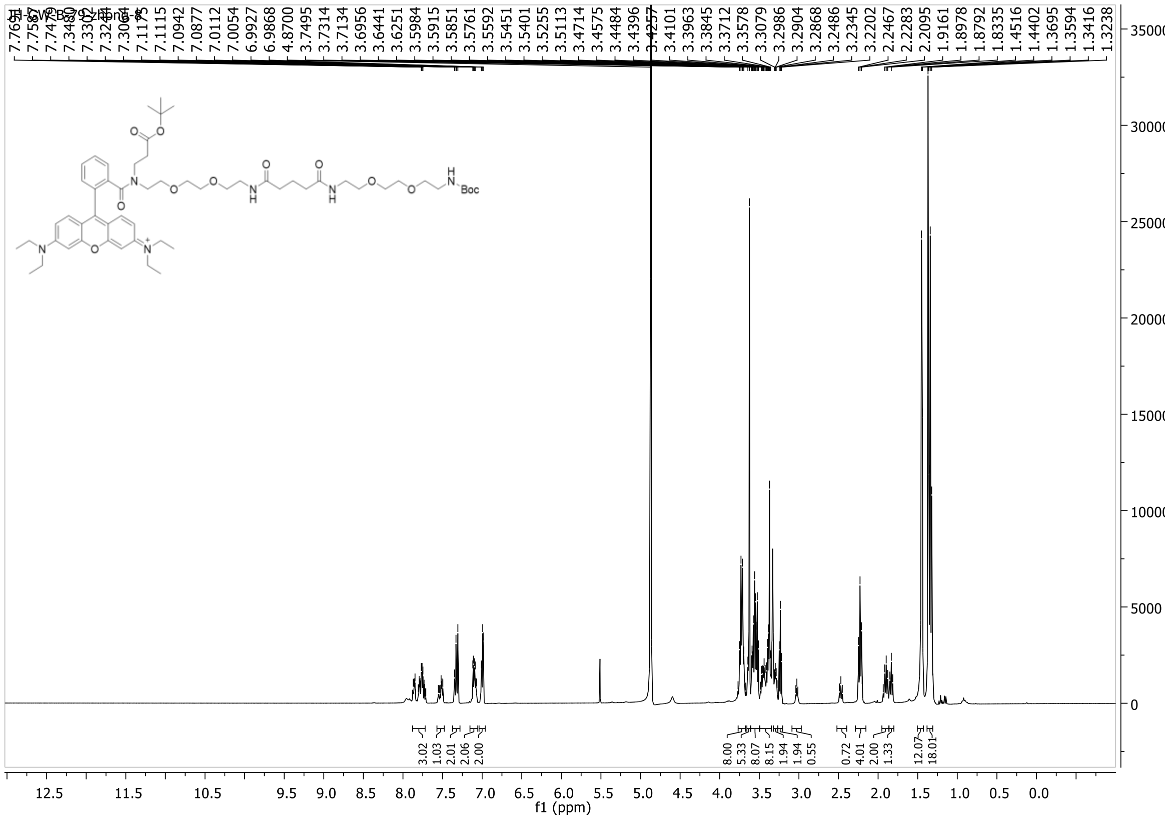

7.1175
7.1115
7.0942
7.0877
7.0112
7.0054
6.9927
6.9868
4.8700
3.7495
3.7314
3.7134
3.6956
3.6441
3.6251
3.5984
3.5915
3.5851
3.5761
3.5592
3.5451
3.5401
3.5255
3.5113
3.4714
3.4575
3.4484
3.4396
3.4257
3.4101
3.3963
3.3845
3.3712
3.3578
3.3079
3.2986
3.2904
3.2868
3.2486
3.2345
3.2202
2.2467
2.2283
2.2095
1.9161
1.8978
1.8792
1.8335
1.4516
1.4402
1.3695
1.3594
1.3416
1.3238
Boc
3.02
1.03
2.01
2.06
2.00
8.00
5.33
8.07
8.15
1.94
1.94
0.55
0.72
4.01
2.00
1.33
12.07
18.01
f1 (ppm)

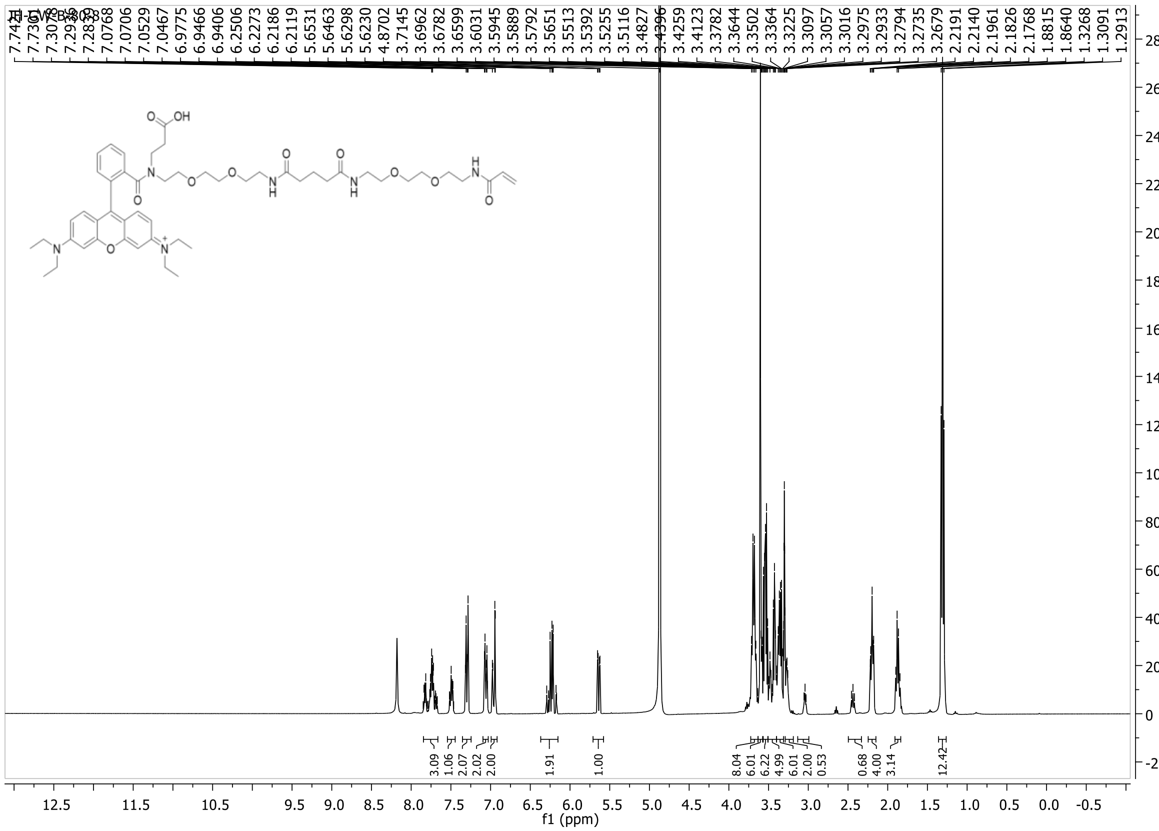

7.0768
7.0706
7.0529
7.0467
6.9775
6.9466
6.9406
6.2506
6.2273
6.2186
6.2119
5.6531
5.6463
5.6298
5.6230
4.8702
3.7145
3.6962
3.6782
3.6599
3.6031
3.5945
3.5889
3.5792
3.5651
3.5513
3.5392
3.5255
3.5116
3.4827
3.4396
3.4259
3.4123
3.3782
3.3644
3.3502
3.3364
3.3225
3.3097
3.3057
3.3016
3.2975
3.2933
3.2794
3.2735
3.2679
2.2191
2.2140
2.1961
2.1826
2.1768
1.8815
1.8640
1.3268
1.3091
1.2913
3.09
1.06
2.07
2.02
2.00
1.91
1.00
8.04
6.01
6.22
4.99
6.01
2.00
0.53
0.68
4.00
3.14
12.42
f1 (ppm)

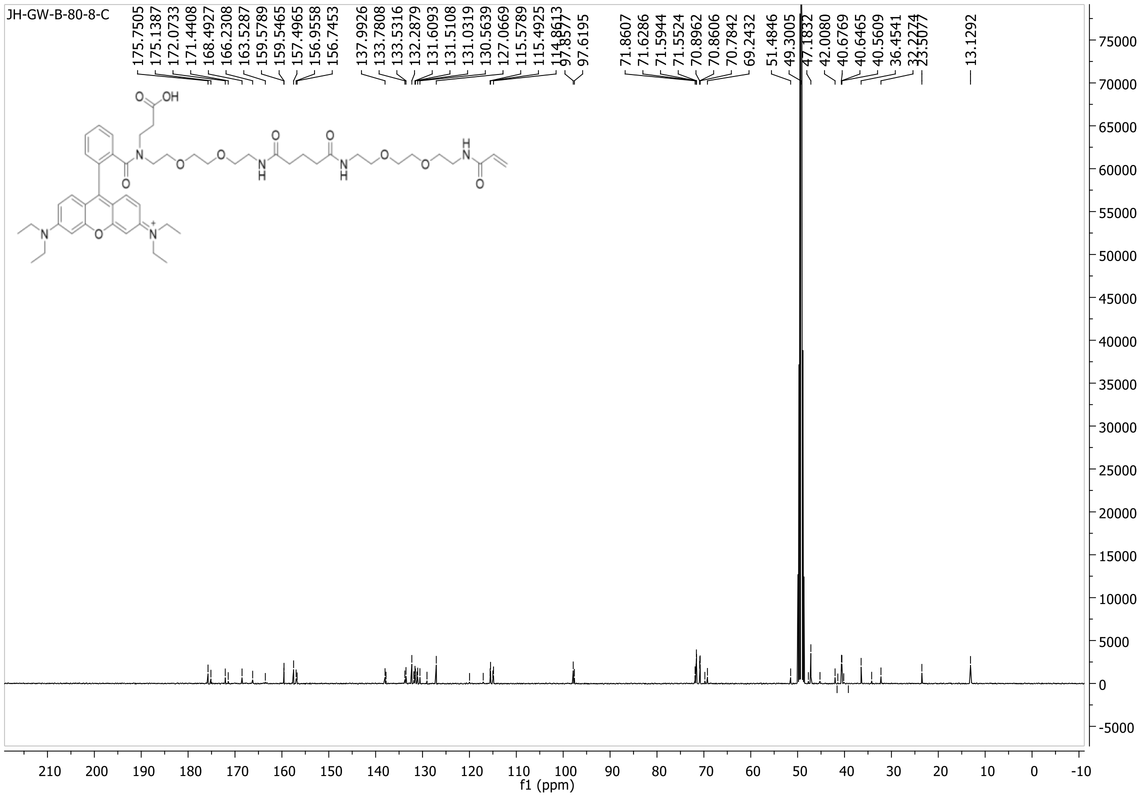
JH-GW-B-80-8-C
175.7505
175.1387
172.0733
171.4408
168.4927
166.2308
163.5287
159.5789
159.5465
157.4965
156.9558
156.7453
137.9926
133.7808
133.5316
132.2879
131.6093
131.5108
131.0319
130.5639
127.0669
115.5789
115.4925
114.8613
97.8577
97.6195
71.8607
71.6286
71.5944
71.5524
70.8962
70.8606
70.7842
69.2432
51.4846
49.3005
47.1832
42.0080
40.6769
40.6465
40.5609
36.4541
32.2274
23.5077
13.1292
f1 (ppm)

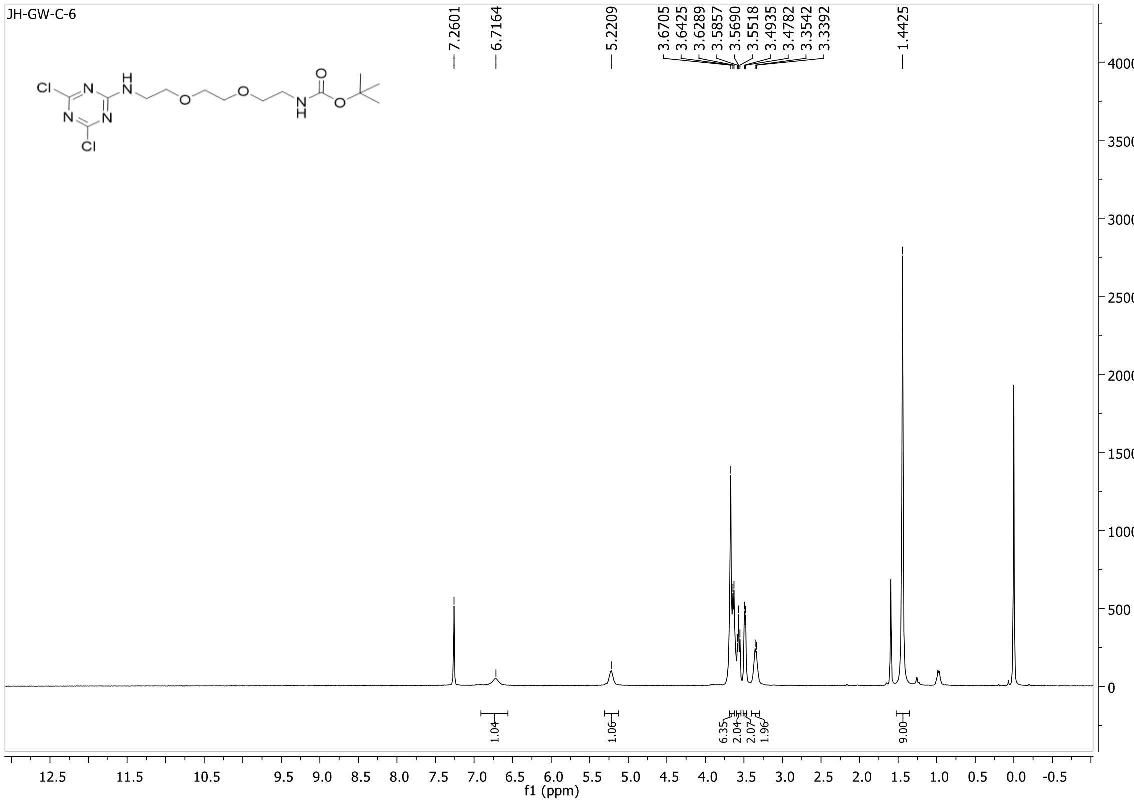
JH-GW-C-6
7.2601
6.7164
5.2209
3.6705
3.6425
3.6289
3.5857
3.5690
3.5518
3.4935
3.4782
3.3542
3.3392
1.4425
1.04
1.06
6.35
2.04
2.07
1.96
9.00
f1 (ppm)

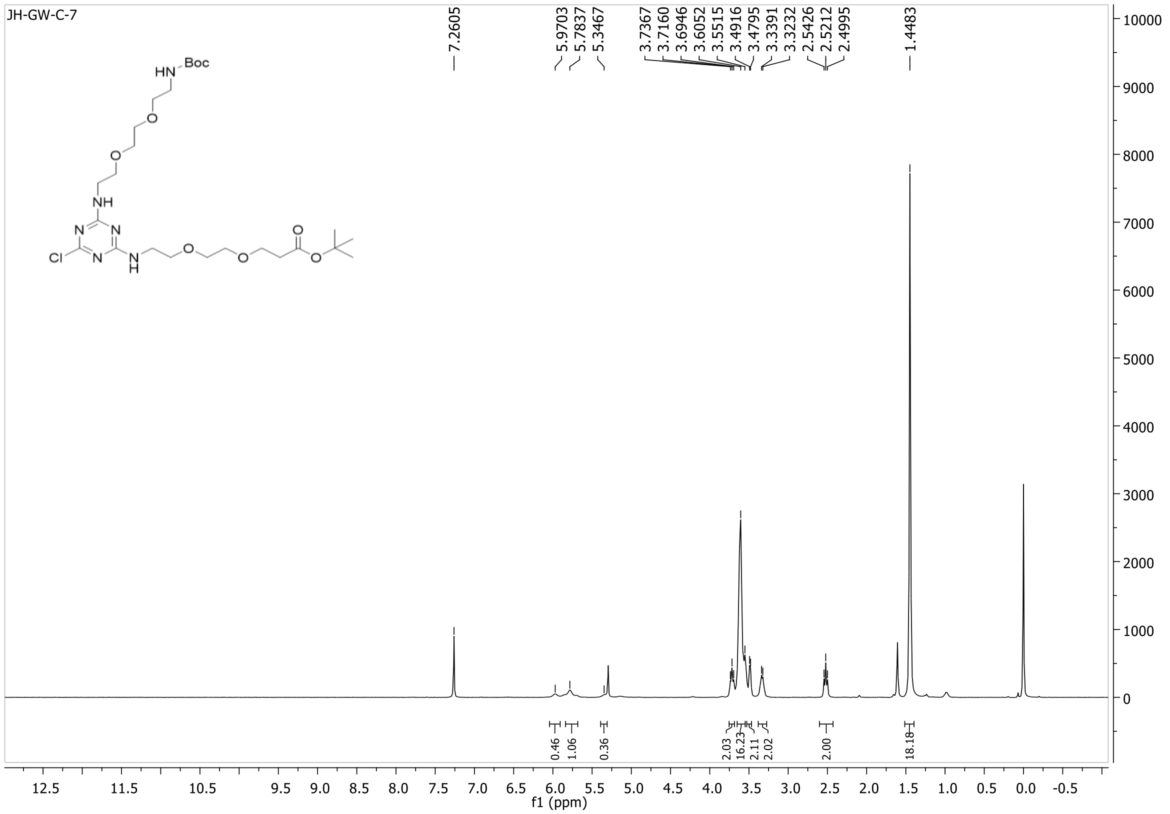
JH-GW-C-7
7.2605
5.9703
5.7837
5.3467
3.7367
3.7160
3.6946
3.6052
3.5515
3.4916
3.4795
3.3391
3.3232
2.5426
2.5212
2.4995
1.4483
0.46
1.06
0.36
2.03
16.23
2.11
2.02
2.00
18.18
f1 (ppm)

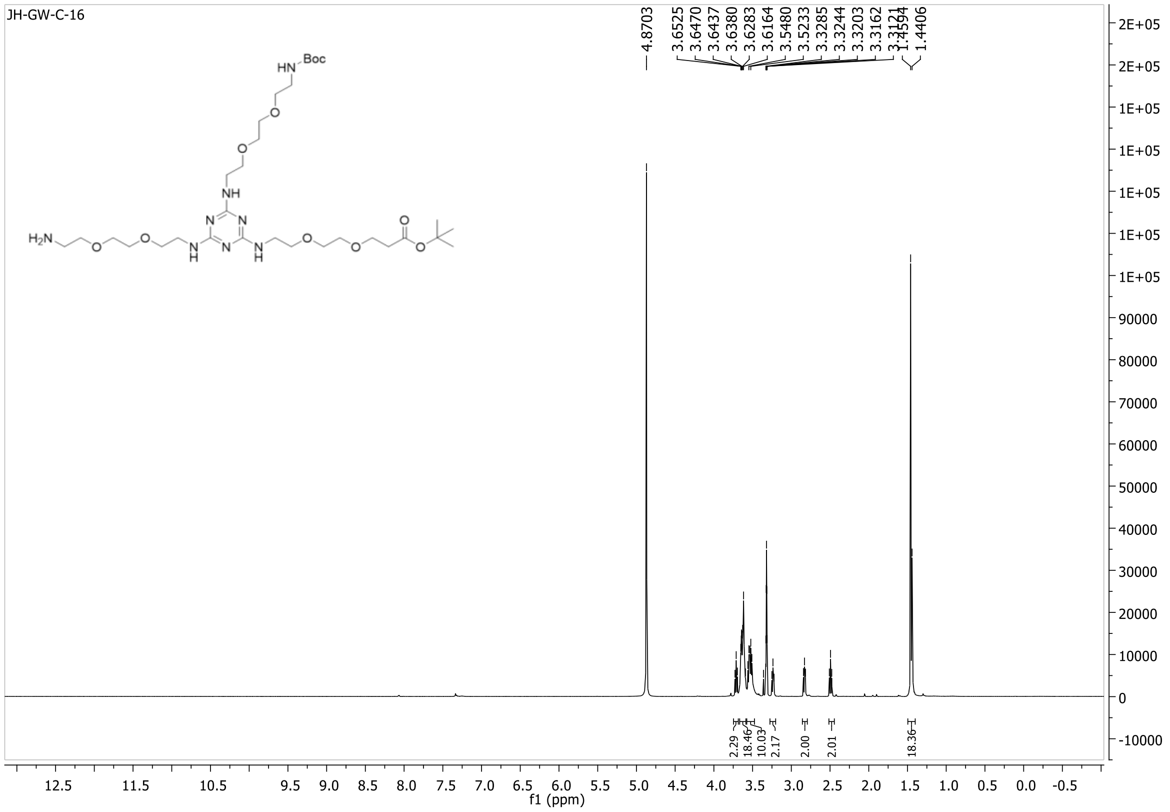
JH-GW-C-16
4.8703
3.6525
3.6470
3.6437
3.6380
3.6283
3.6164
3.5480
3.5233
3.3285
3.3244
3.3203
3.3162
3.3121
1.4594
1.4406
2.29
18.46
10.03
2.17
2.00
2.01
18.36
f1 (ppm)

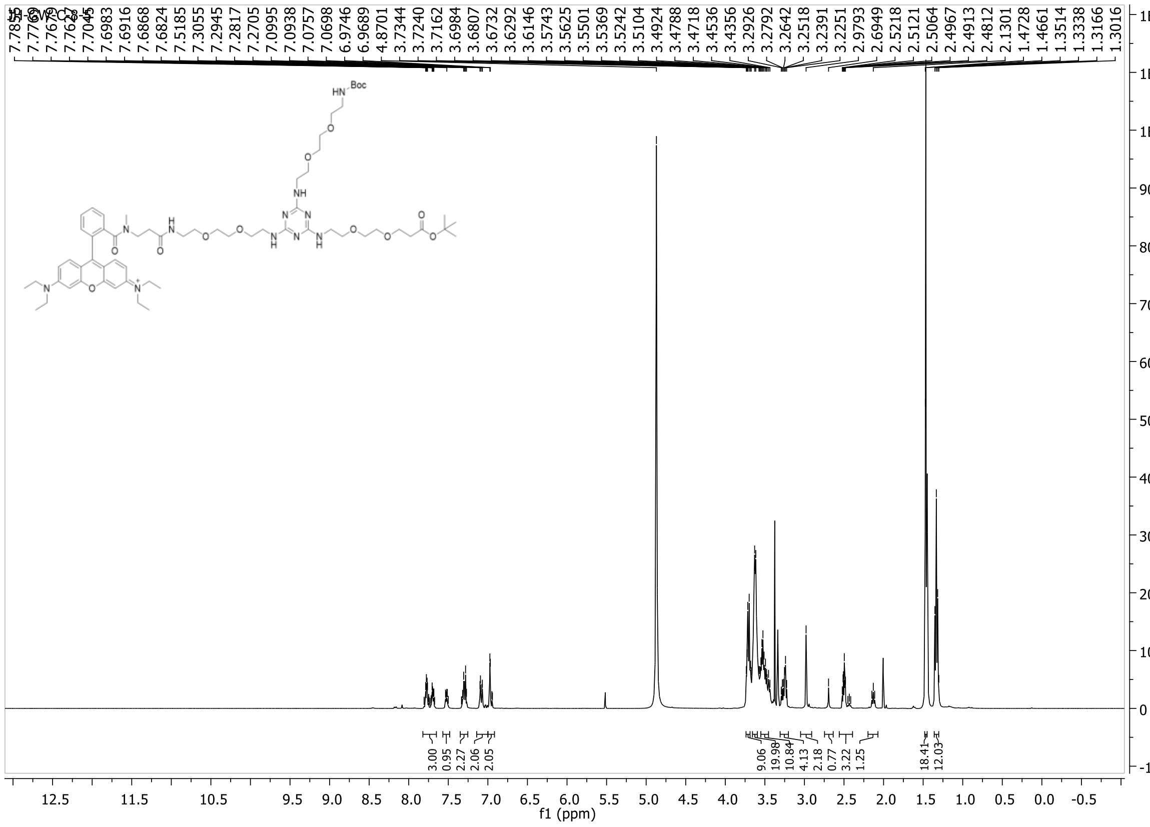
7.7045
7.6983
7.6916
7.6868
7.6824
7.5185
7.3055
7.2945
7.2817
7.2705
7.0995
7.0938
7.0757
7.0698
6.9746
6.9689
4.8701
3.7344
3.7240
3.7162
3.6984
3.6807
3.6732
3.6292
3.6146
3.5743
3.5625
3.5501
3.5369
3.5242
3.5104
3.4924
3.4788
3.4718
3.4536
3.4356
3.2926
3.2792
3.2642
3.2518
3.2391
3.2251
2.9793
2.6949
2.5218
2.5121
2.5064
2.4967
2.4913
2.4812
2.1301
1.4728
1.4661
1.3514
1.3338
1.3166
1.3016
HN Boc
O
NH
N
H
N
O
O
N
H
N
N
N
N
H
O
O
O
O
O
N
N+
3.00
0.95
2.27
2.06
2.05
9.06
19.98
10.84
4.13
2.18
0.77
3.22
1.25
18.41
12.03
f1 (ppm)

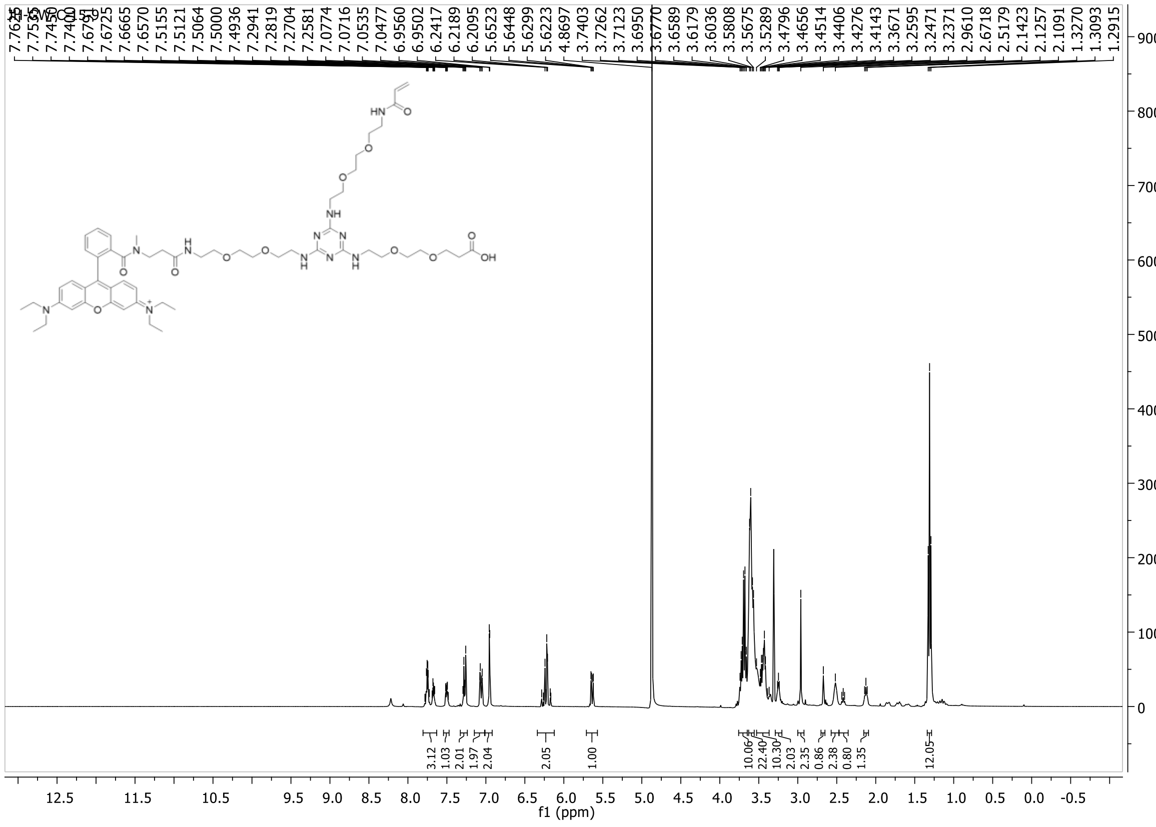
7.7400
7.6791
7.6725
7.6665
7.6570
7.5155
7.5121
7.5064
7.5000
7.4936
7.2941
7.2819
7.2704
7.2581
7.0774
7.0716
7.0535
7.0477
6.9560
6.9502
6.2417
6.2189
6.2095
5.6523
5.6448
5.6299
5.6223
4.8697
3.7403
3.7262
3.7123
3.6950
3.6770
3.6589
3.6179
3.6036
3.5808
3.5675
3.5289
3.4796
3.4656
3.4514
3.4406
3.4276
3.4143
3.3671
3.2595
3.2471
3.2371
2.9610
2.6718
2.5179
2.1423
2.1257
2.1091
1.3270
1.3093
1.2915
HN
O
OH
3.12
1.03
2.01
1.97
2.04
2.05
1.00
10.06
22.40
10.30
2.03
2.35
0.86
2.38
0.80
1.35
12.05
f1 (ppm)

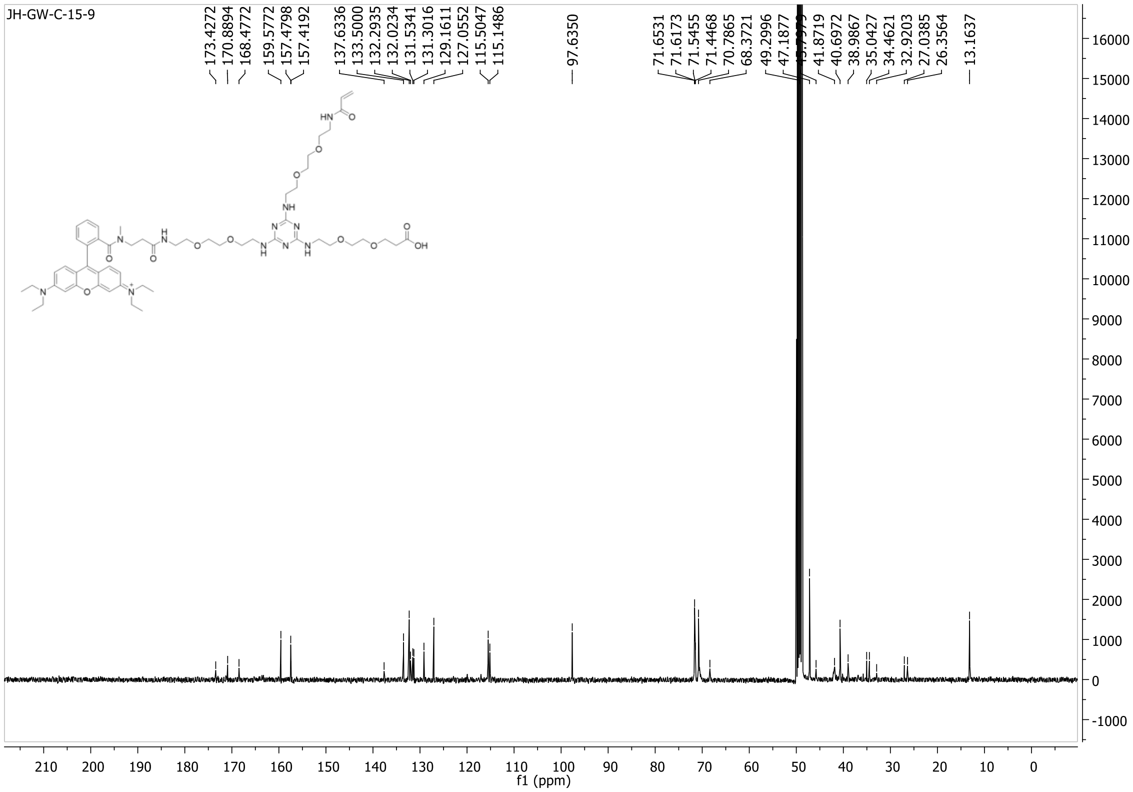
JH-GW-C-15-9
173.4272
170.8894
168.4772
159.5772
157.4798
157.4192
137.6336
133.5000
132.2935
132.0234
131.5341
131.3016
129.1611
127.0552
115.5047
115.1486
97.6350
71.6531
71.6173
71.5455
71.4468
70.7865
68.3721
49.2996
47.1877
41.8719
40.6972
38.9867
35.0427
34.4621
32.9203
27.0385
26.3564
13.1637
f1 (ppm)

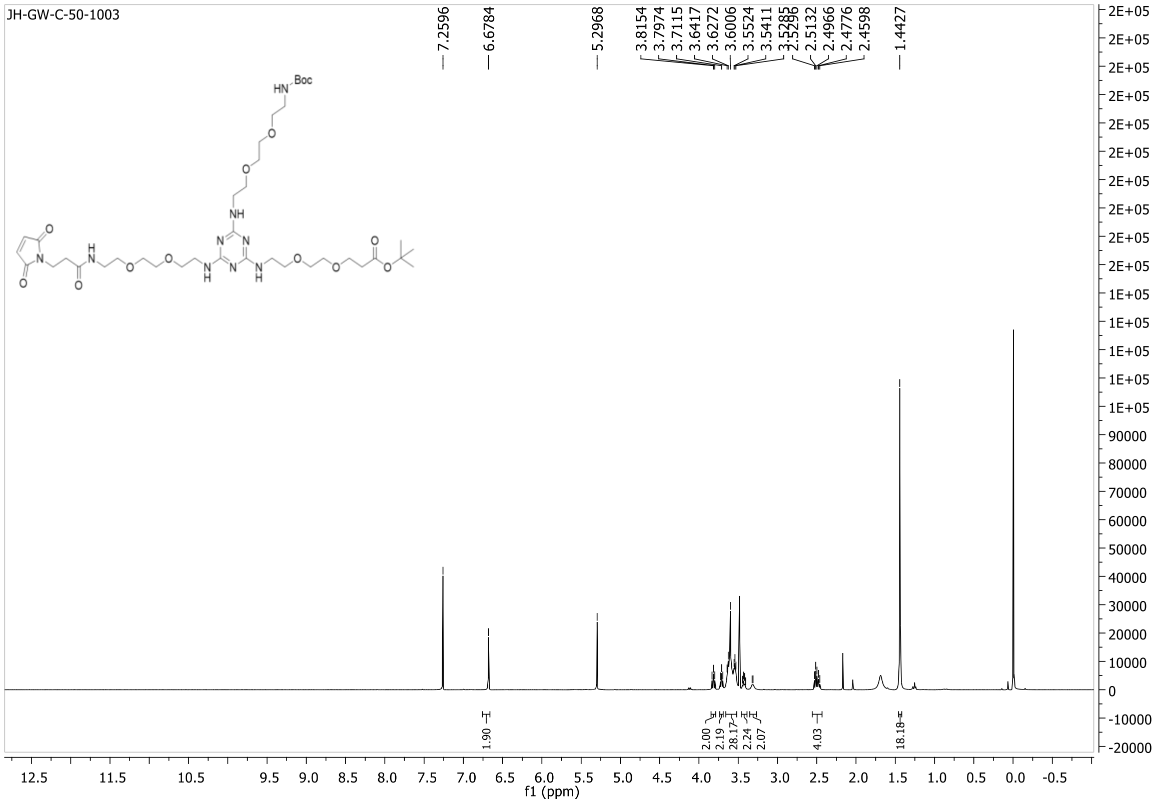
JH-GW-C-50-1003
7.2596
6.6784
5.2968
3.8154
3.7974
3.7115
3.6417
3.6272
3.6006
3.5524
3.5411
3.5285
3.5296
2.5132
2.4966
2.4776
2.4598
1.4427
1.90
2.00
2.19
28.17
2.24
2.07
4.03
18.18
f1 (ppm)

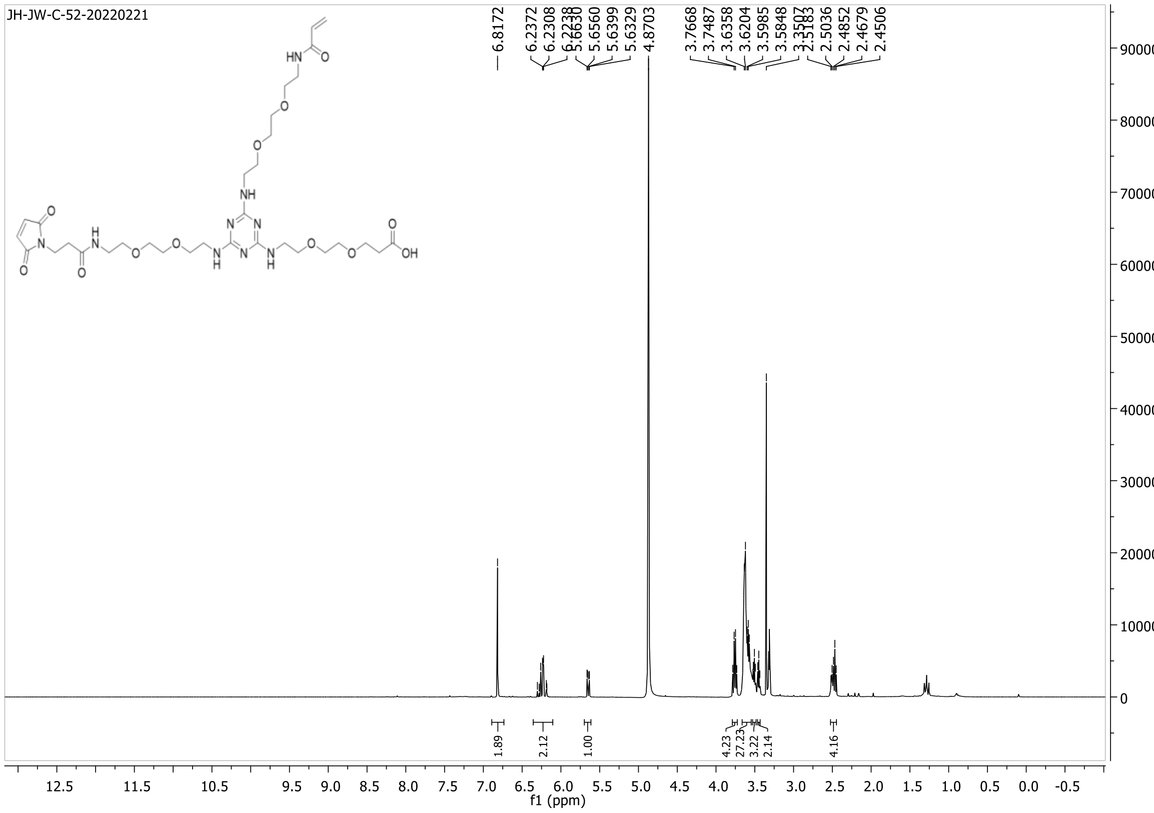
JH-JW-C-52-20220221
6.8172
6.2372
6.2308
6.2238
5.6630
5.6560
5.6399
5.6329
4.8703
3.7668
3.7487
3.6358
3.6204
3.5985
3.5848
3.3507
2.5183
2.5036
2.4852
2.4679
2.4506
1.89
2.12
1.00
4.23
27.23
3.22
2.14
4.16
f1 (ppm)

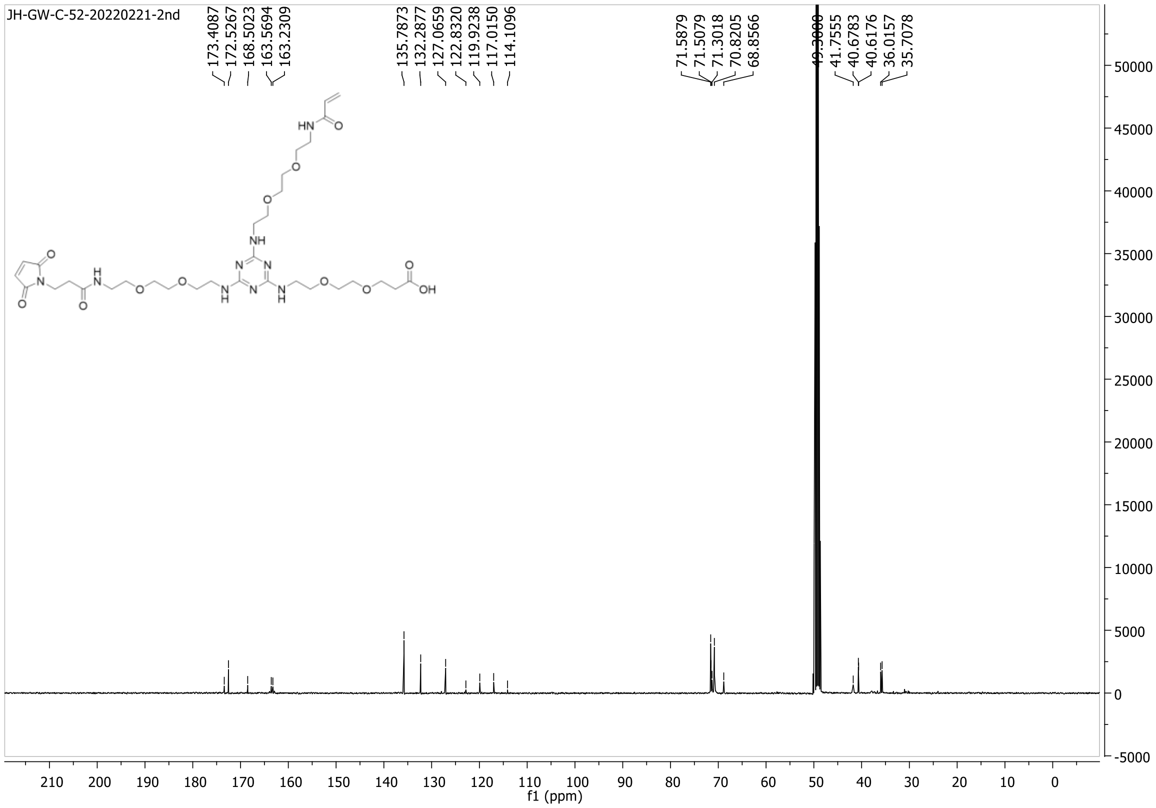
JH-GW-C-52-20220221-2nd
173.4087
172.5267
168.5023
163.5694
163.2309
135.7873
132.2877
127.0659
122.8320
119.9238
117.0150
114.1096
71.5879
71.5079
71.3018
70.8205
68.8566
41.7555
40.6783
40.6176
36.0157
35.7078
f1 (ppm)

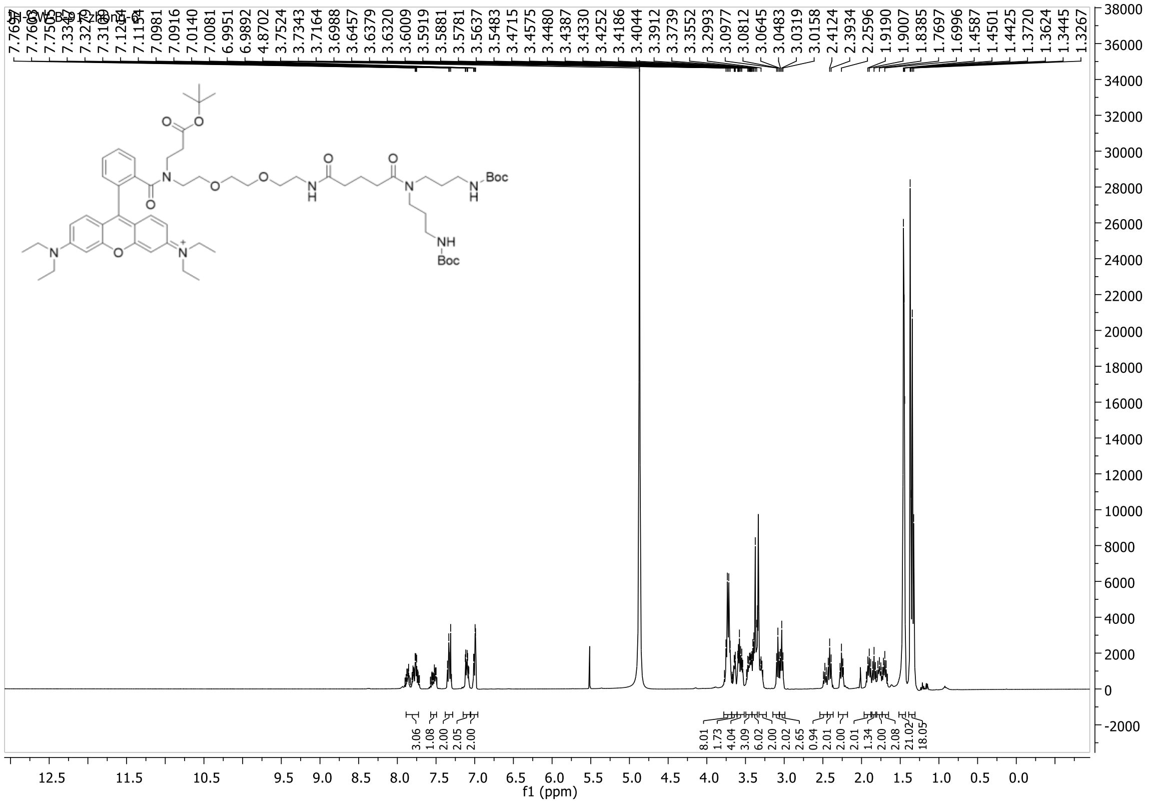

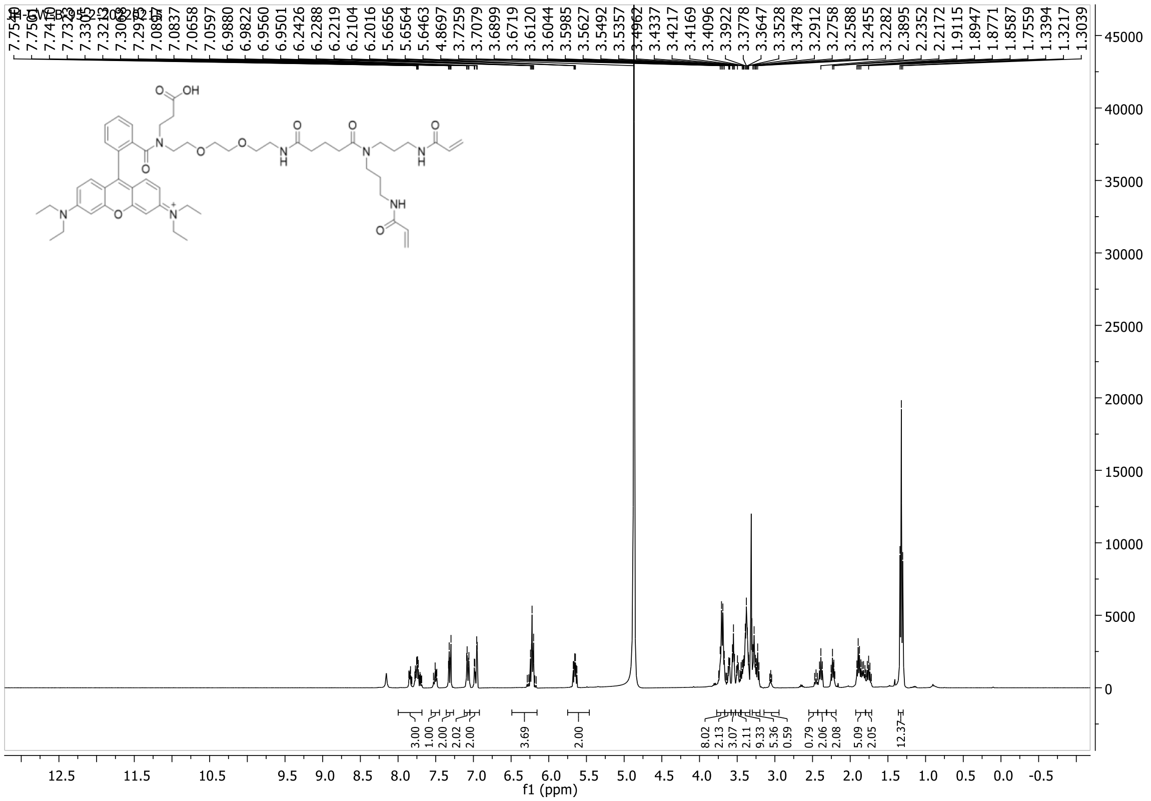

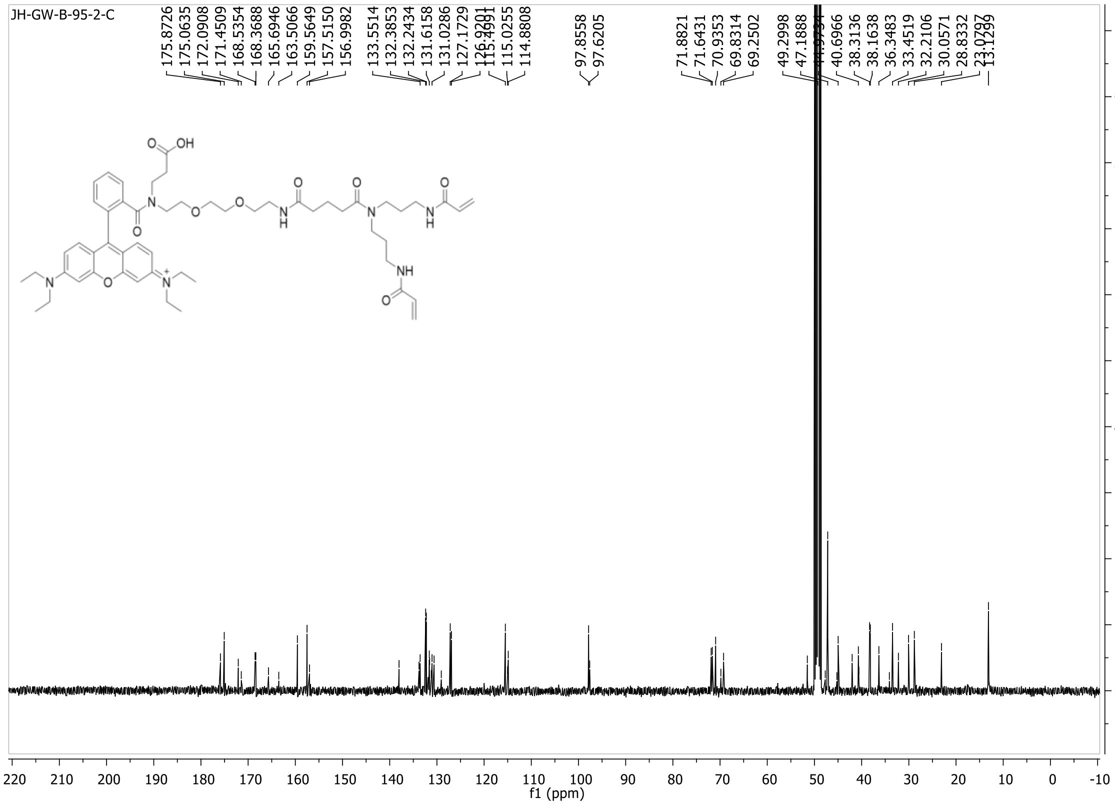
JH-GW-B-95-2-C
175.8726
175.0635
172.0908
171.4509
168.5354
168.3688
165.6946
163.5066
159.5649
157.5150
156.9982
133.5514
132.3853
132.2434
131.6158
131.0286
127.1729
115.0255
114.8808
97.8558
97.6205
71.8821
71.6431
70.9353
69.8314
69.2502
49.2998
47.1888
40.6966
38.3136
38.1638
36.3483
33.4519
32.2106
30.0571
28.8332
13.1299
f1 (ppm)

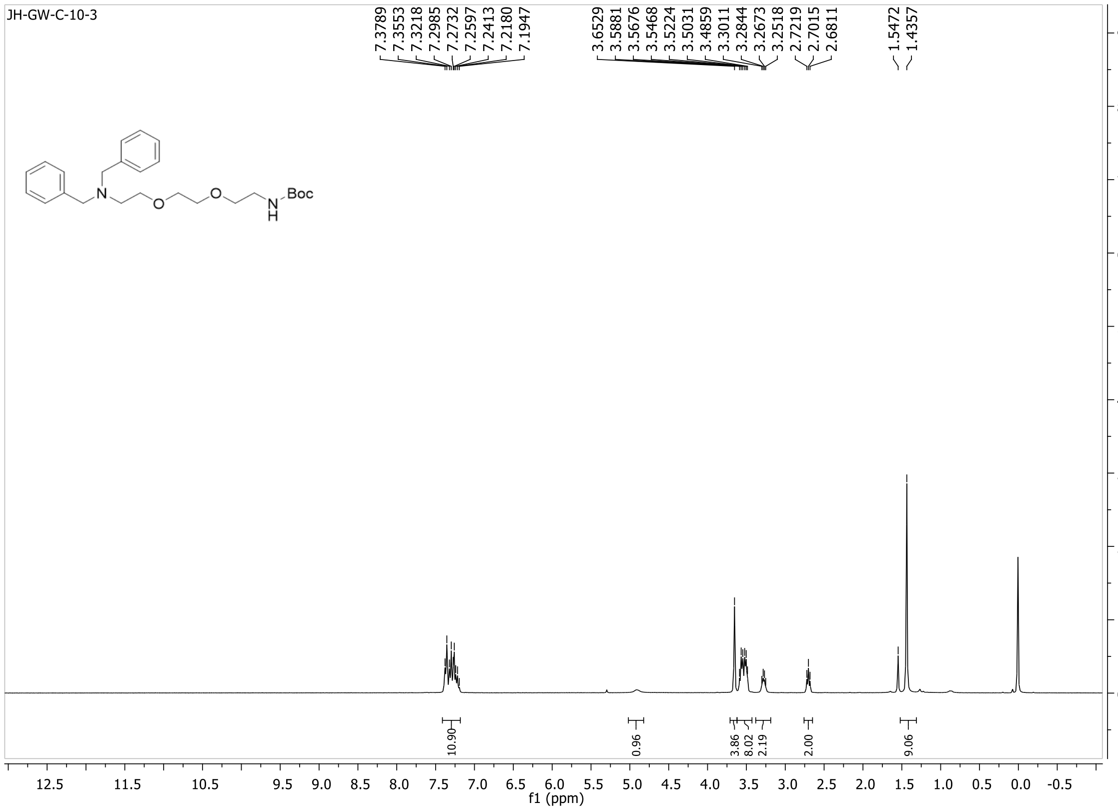
JH-GW-C-10-3
7.3789
7.3553
7.3218
7.2985
7.2732
7.2597
7.2413
7.2180
7.1947
3.6529
3.5881
3.5676
3.5468
3.5224
3.5031
3.4859
3.3011
3.2844
3.2673
3.2518
2.7219
2.7015
2.6811
1.5472
1.4357
Boc
10.90
0.96
3.86
8.02
2.19
2.00
9.06
f1 (ppm)

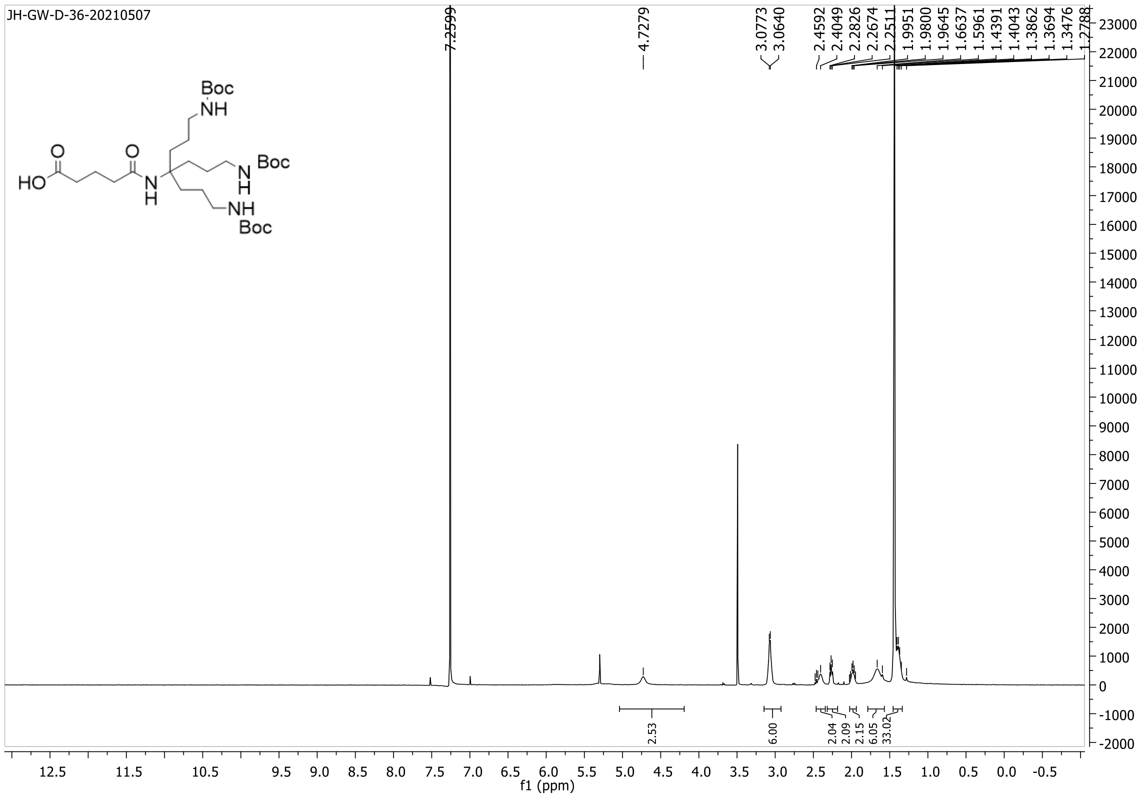
JH-GW-D-36-20210507
Boc
NH
HO
O
O
N
H
N
H
Boc
NH
Boc
f1 (ppm)

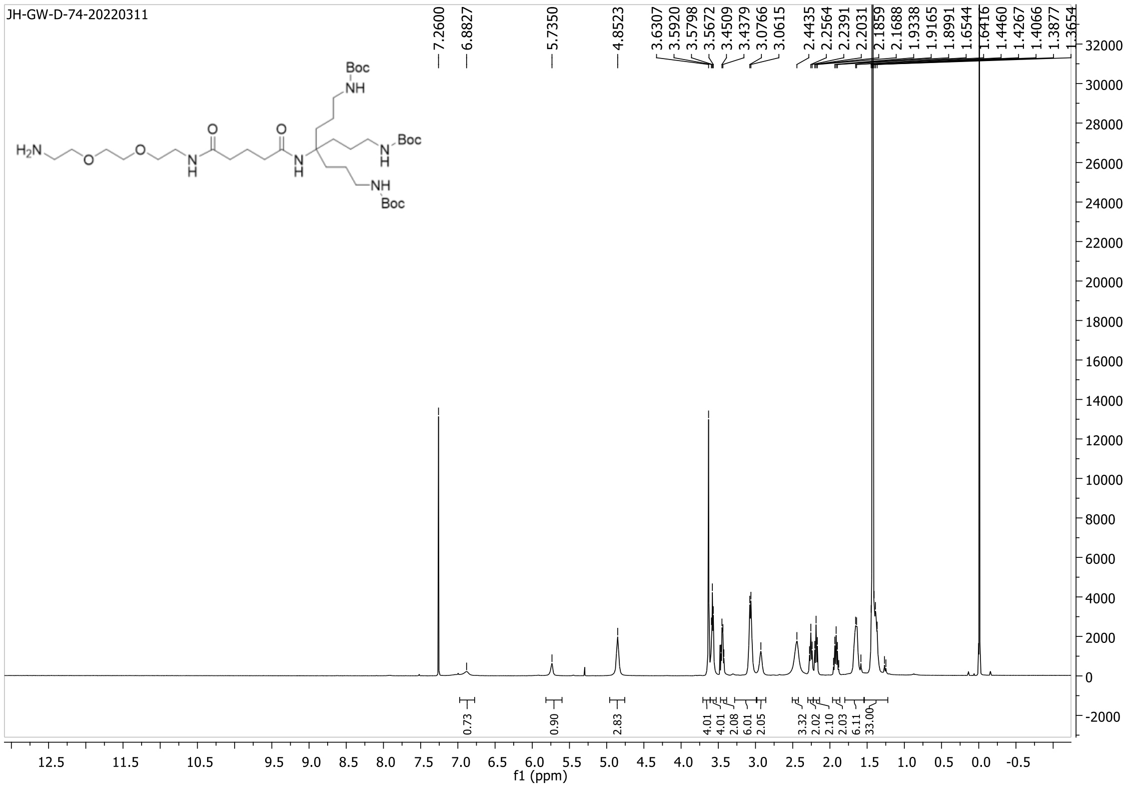
JH-GW-D-74-20220311
Boc
NH
$H_2N$
O
O
N
H
Boc
f1 (ppm)

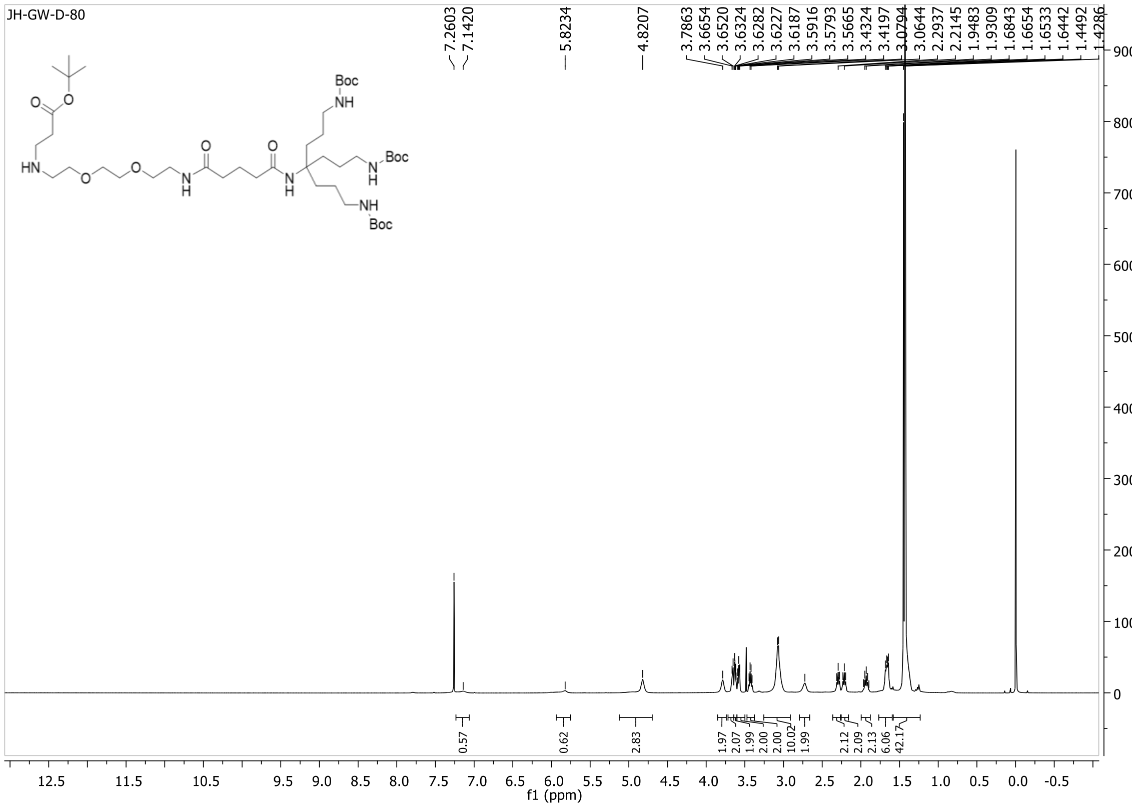
JH-GW-D-80
f1 (ppm)

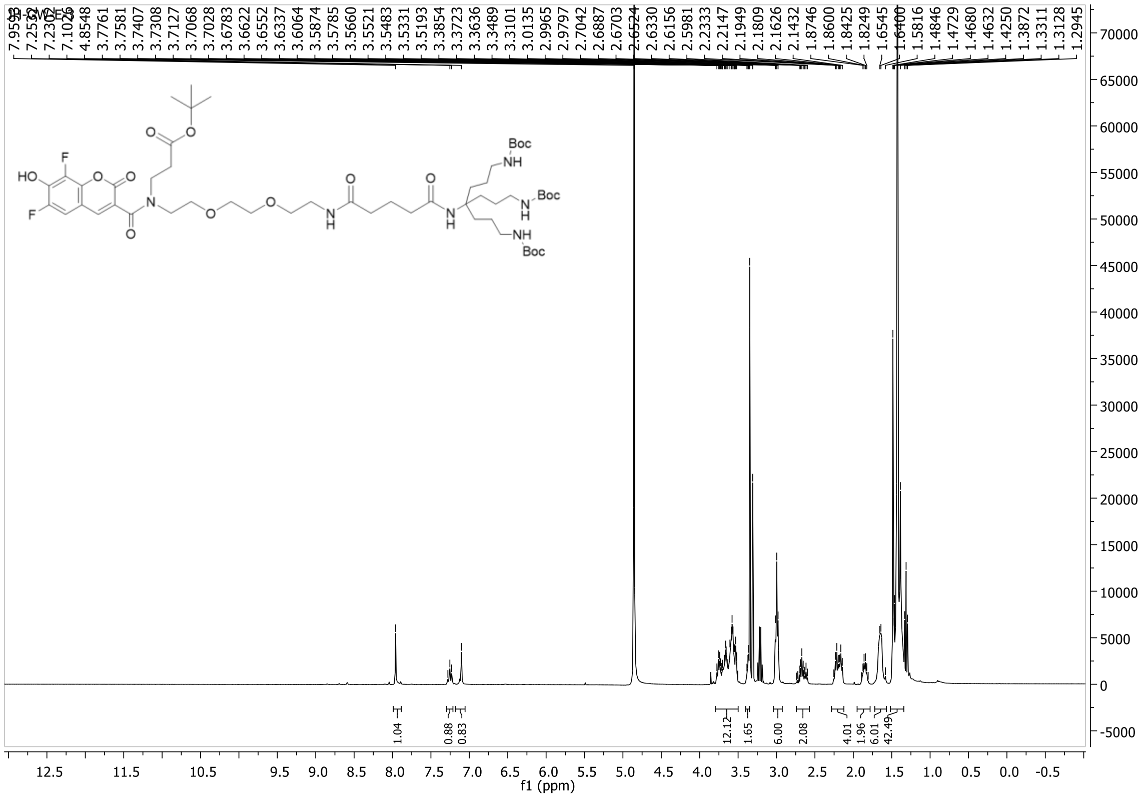
f1 (ppm)

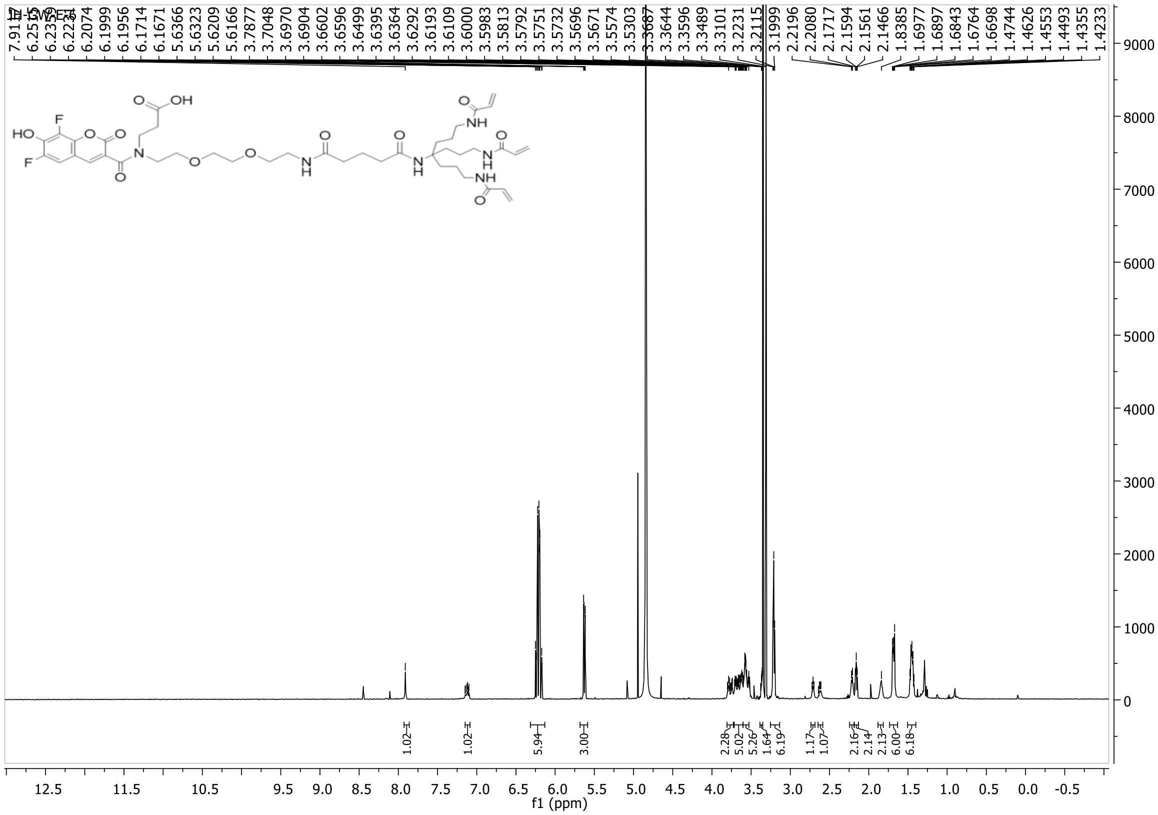
f1 (ppm)

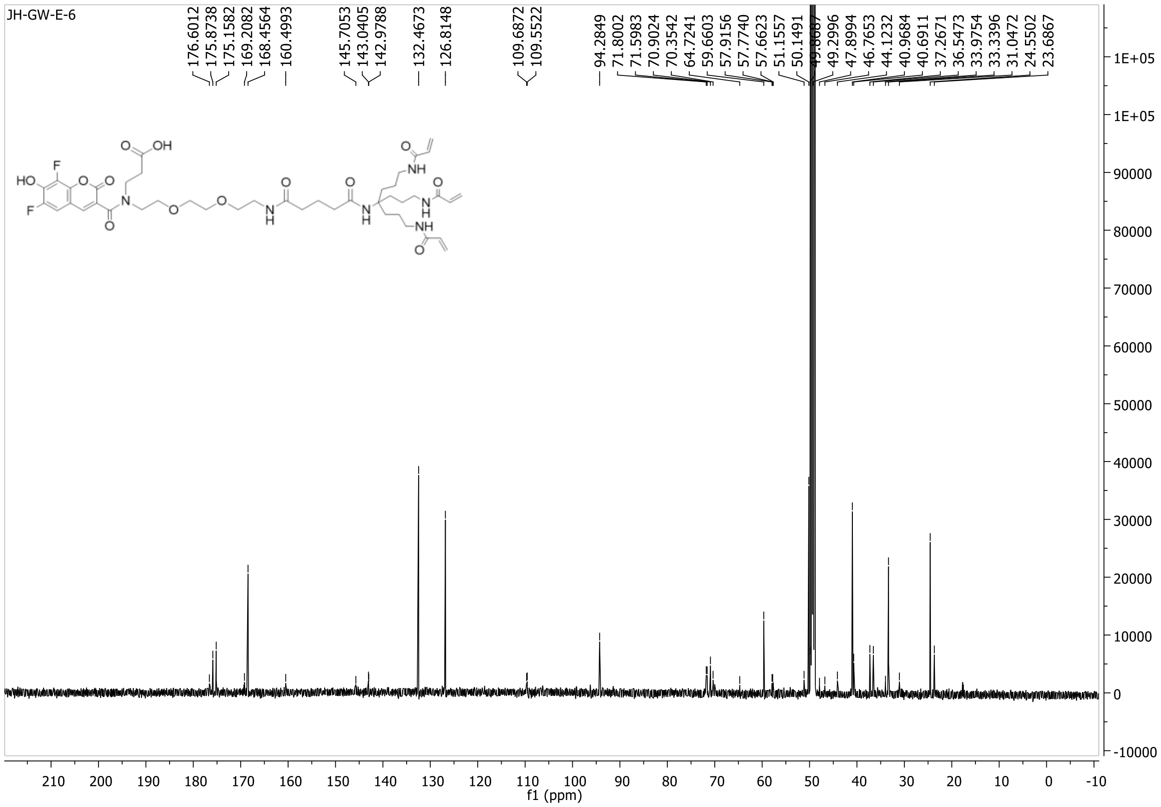
JH-GW-E-6
f1 (ppm)

**LC of compound S17**

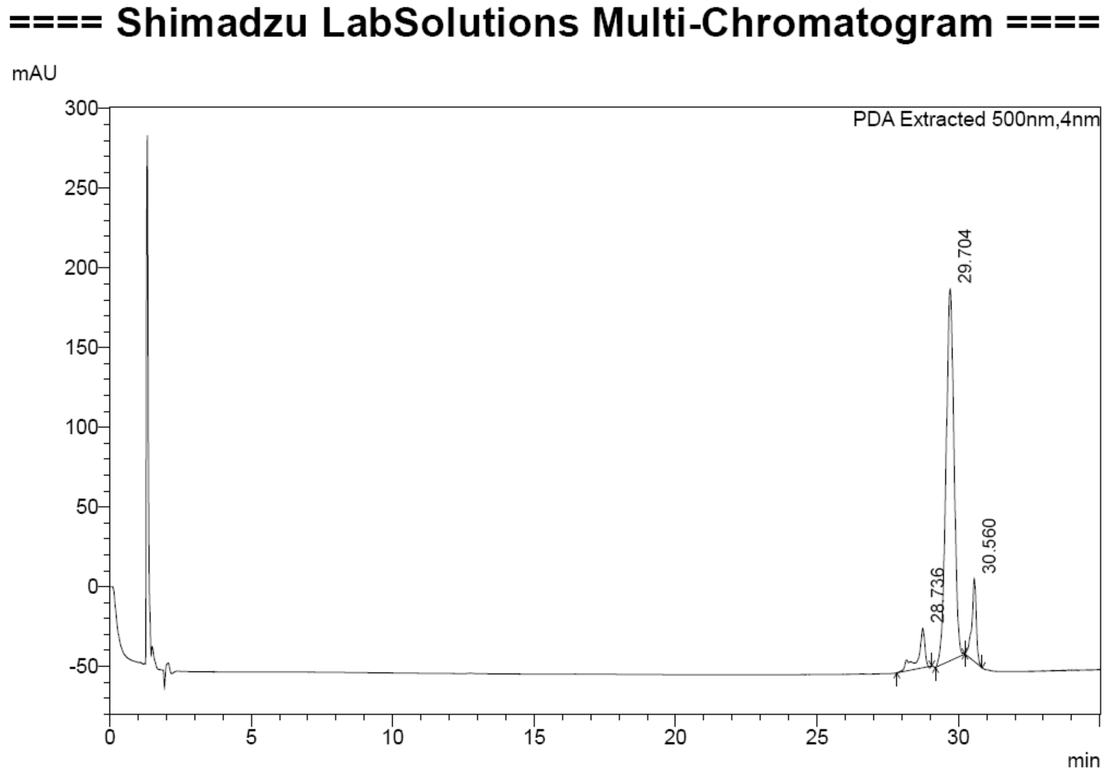

**MS of compound S17**

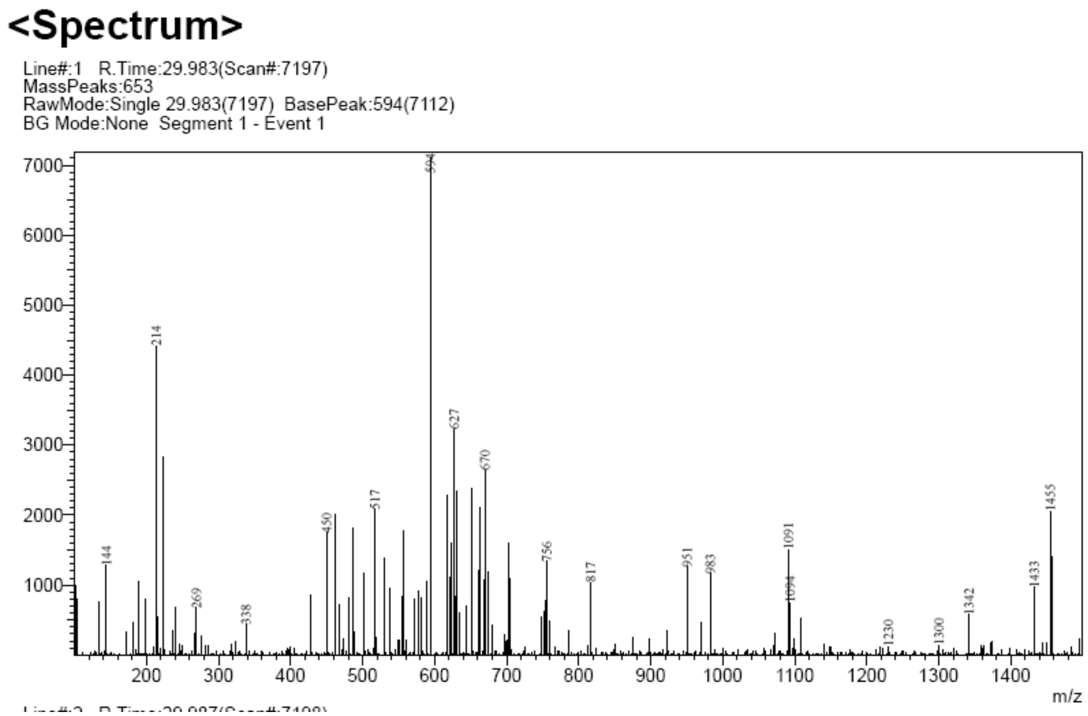

**LC of compound S18**

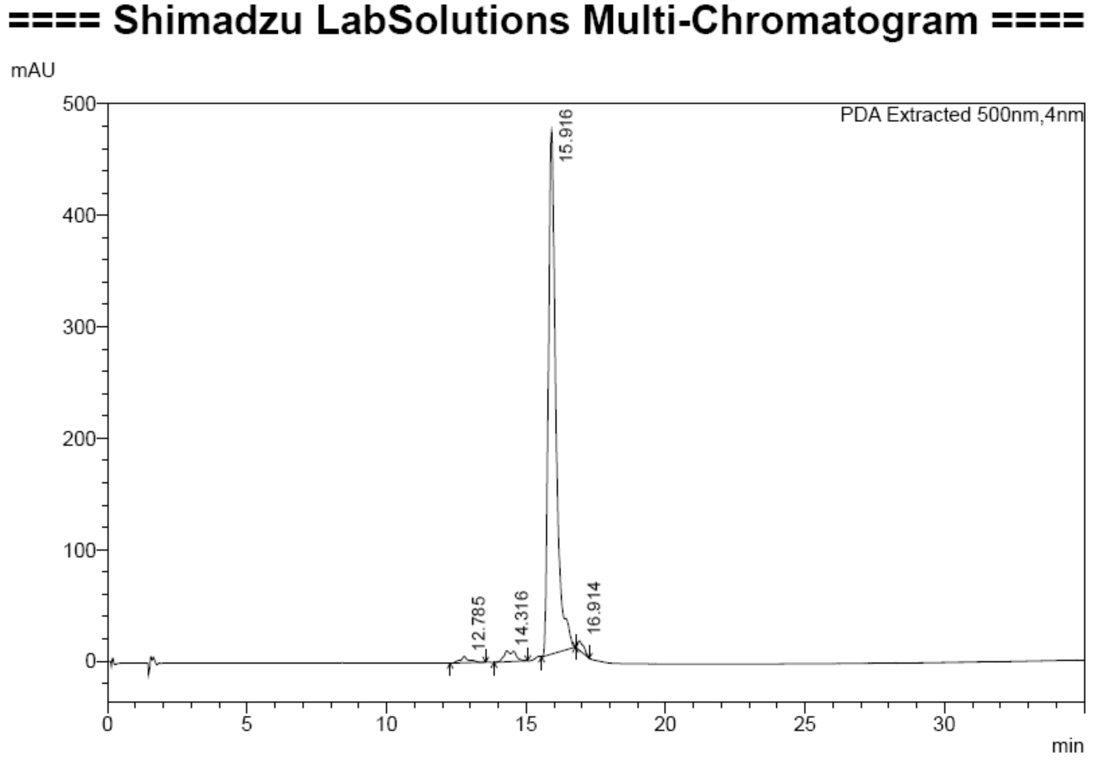

**MS of compound S18**

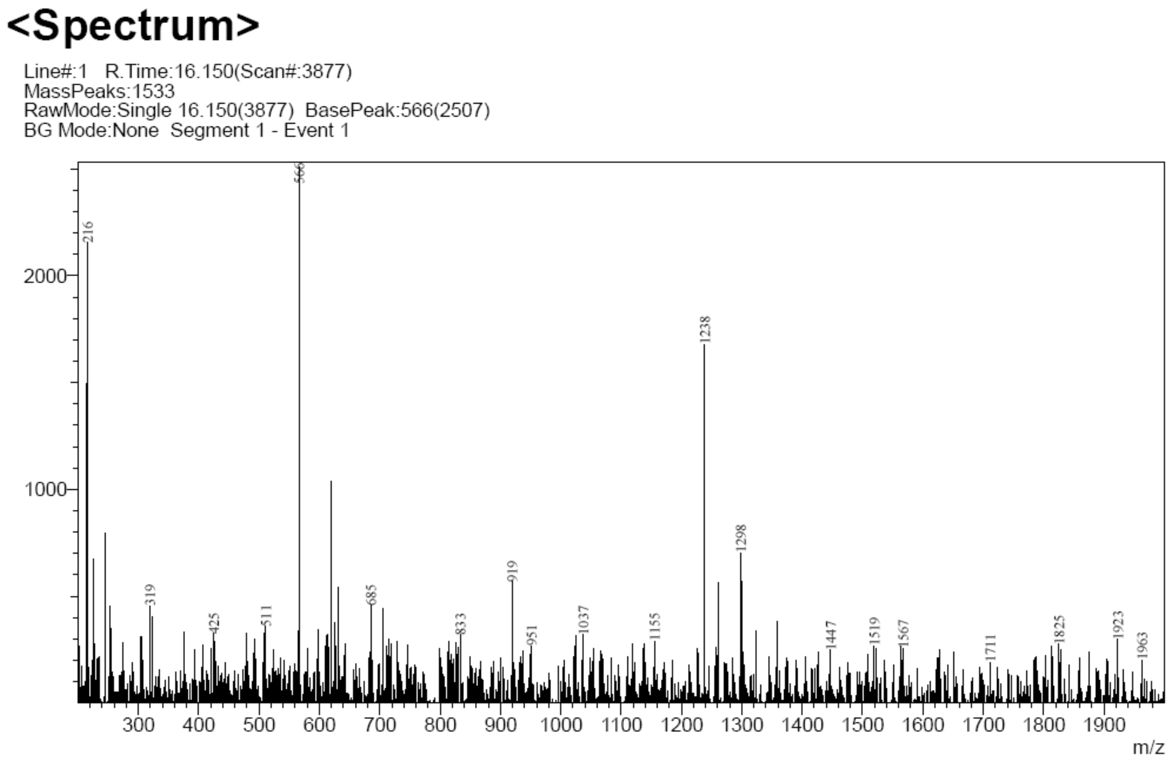

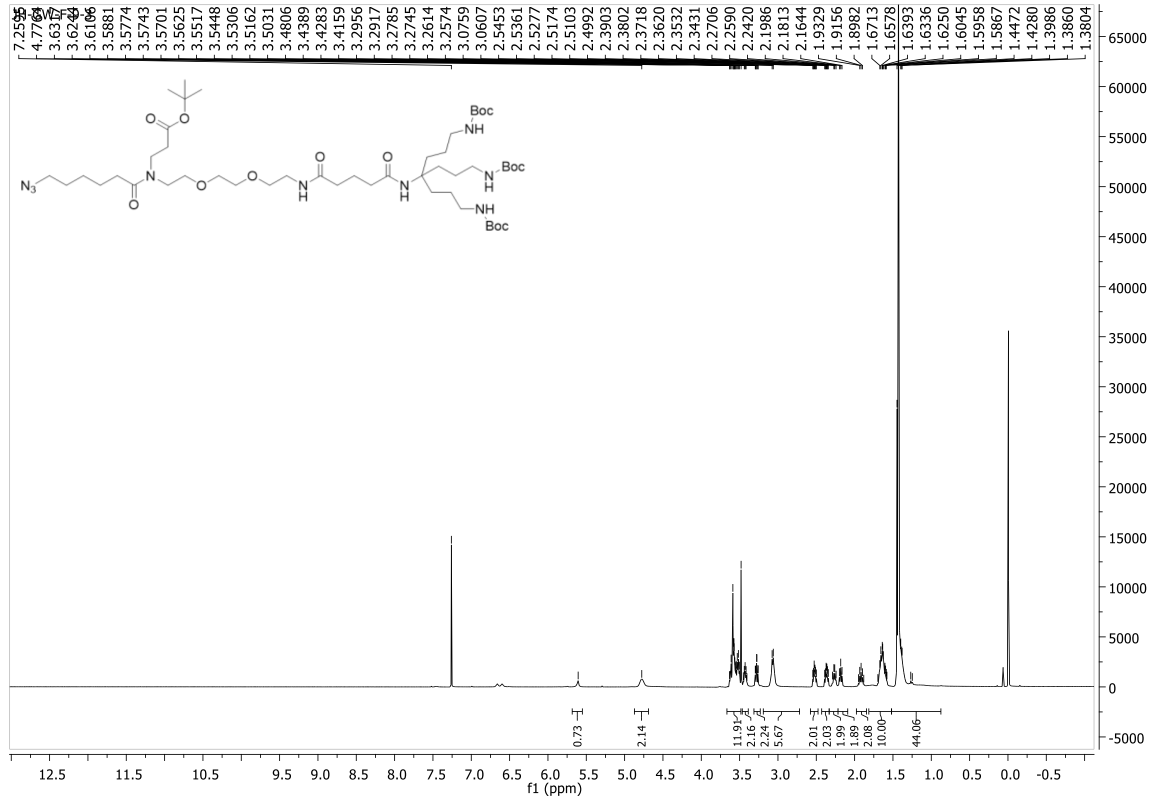

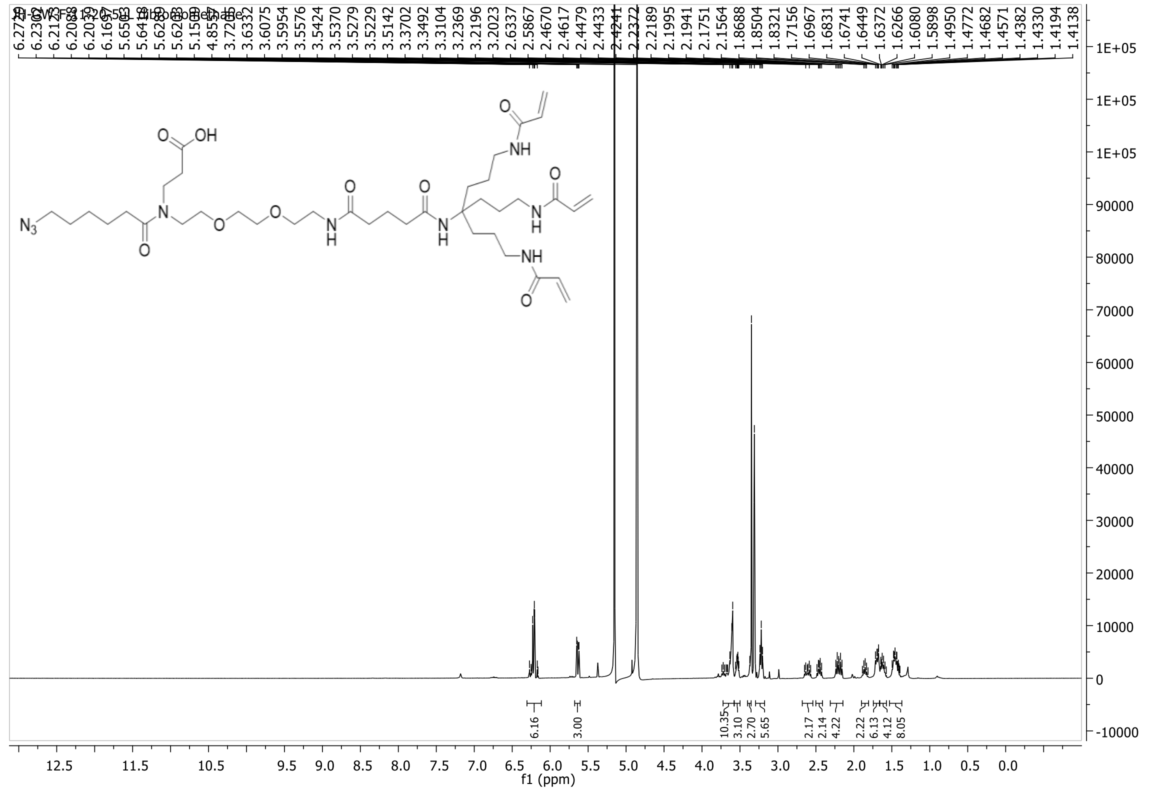

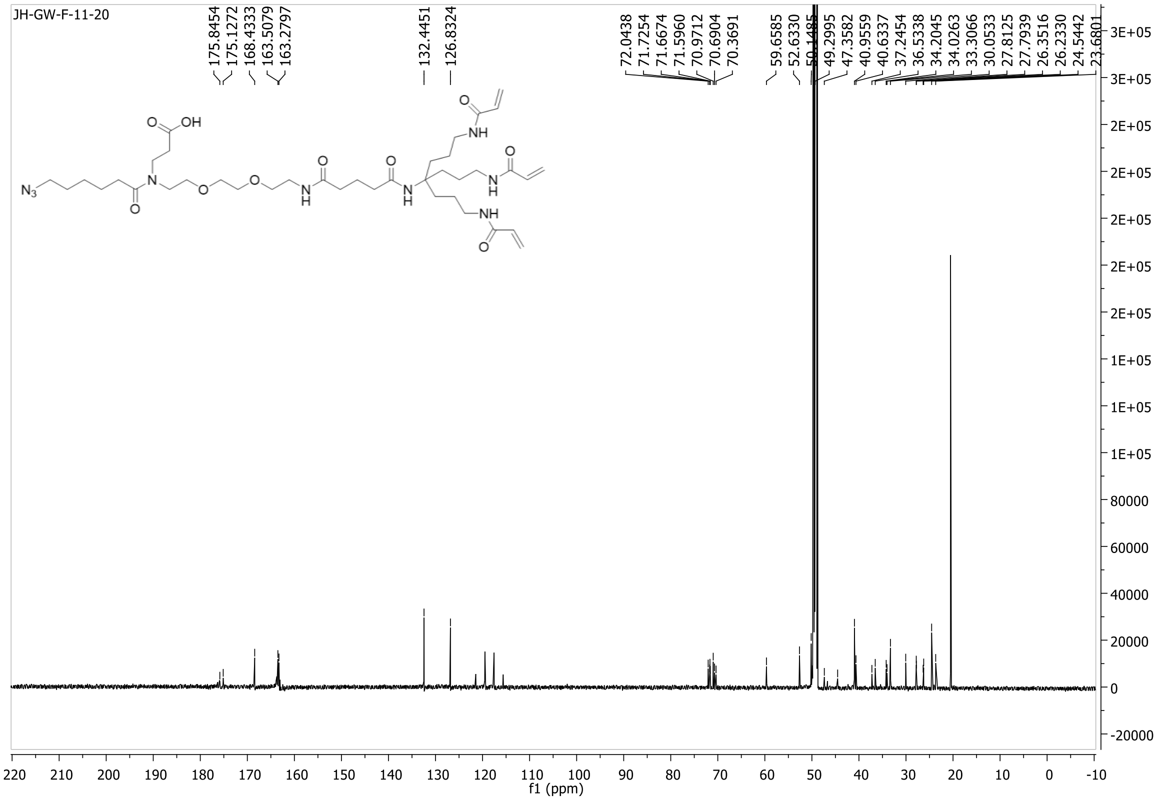
JH-GW-F-11-20
175.8454
175.1272
168.4333
163.5079
163.2797
132.4451
126.8324
72.0438
71.7254
71.6674
71.5960
70.9712
70.6904
70.3691
59.6585
52.6330
49.2995
47.3582
40.9559
40.6337
37.2454
36.5338
34.2045
34.0263
33.3066
30.0533
27.8125
27.7939
26.3516
26.2330
24.5442
N3
O
OH
N
O
O
NH
NH
H
N
H
N
f1 (ppm)

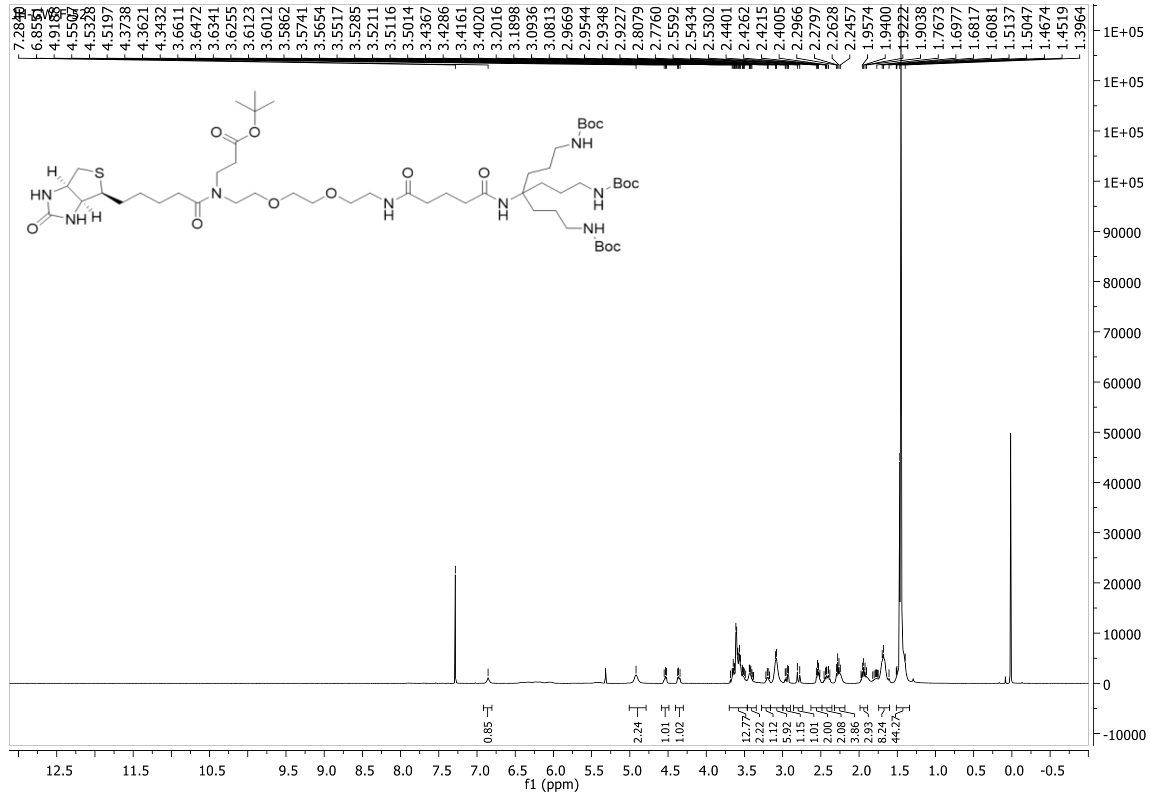
7.2844
6.8555
4.9188
4.5507
4.5328
4.5197
4.3738
4.3621
4.3432
3.6611
3.6472
3.6341
3.6255
3.6123
3.6012
3.5862
3.5741
3.5654
3.5517
3.5285
3.5211
3.5116
3.5014
3.4367
3.4286
3.4161
3.4020
3.2016
3.1898
3.0936
3.0813
2.9669
2.9544
2.9348
2.9227
2.8079
2.7760
2.5592
2.5434
2.5302
2.4401
2.4262
2.4215
2.4005
2.2966
2.2797
2.2628
2.2457
1.9574
1.9400
1.9222
1.9038
1.7673
1.6977
1.6817
1.6081
1.5137
1.5047
1.4674
1.4519
1.3964
Boc
NH
HN
S
H
O
0.85
2.24
1.01
1.02
12.77
2.22
1.12
5.92
1.15
1.01
2.00
2.08
3.86
2.93
8.24
44.27
f1 (ppm)

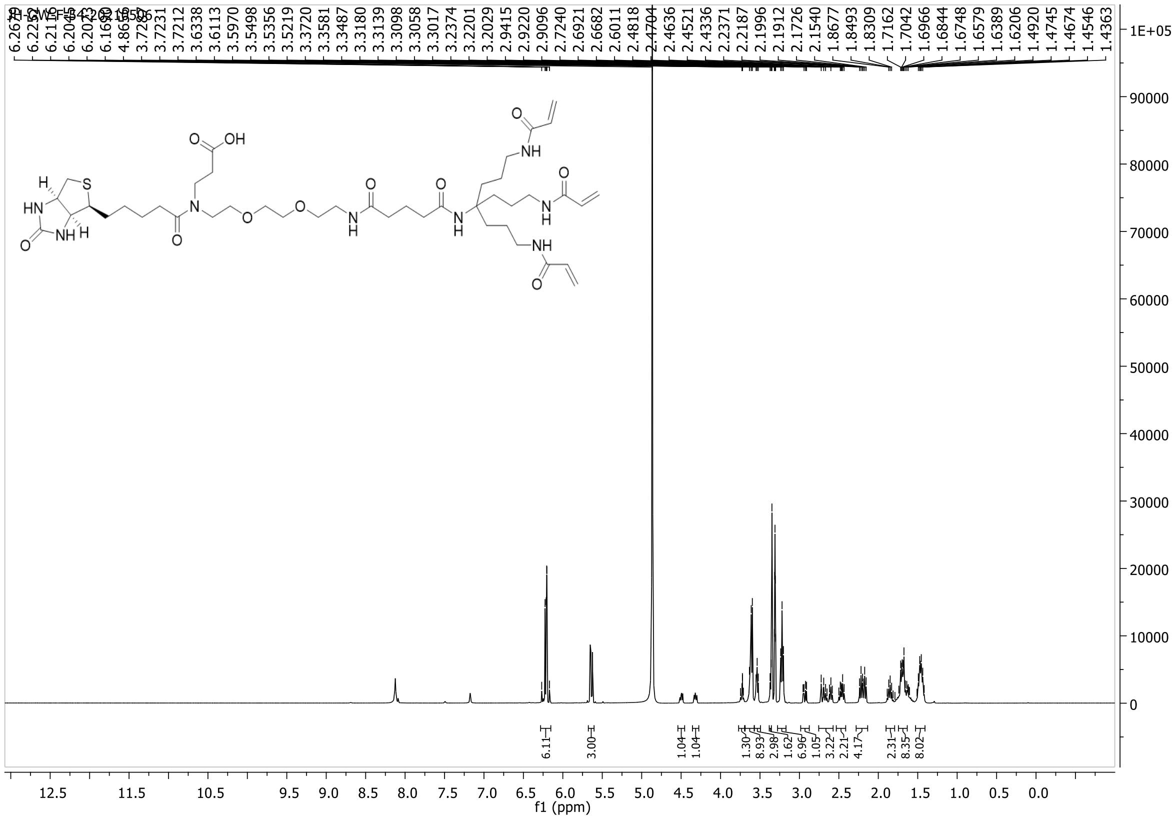
JH-GW-F-54-20210506
f1 (ppm)

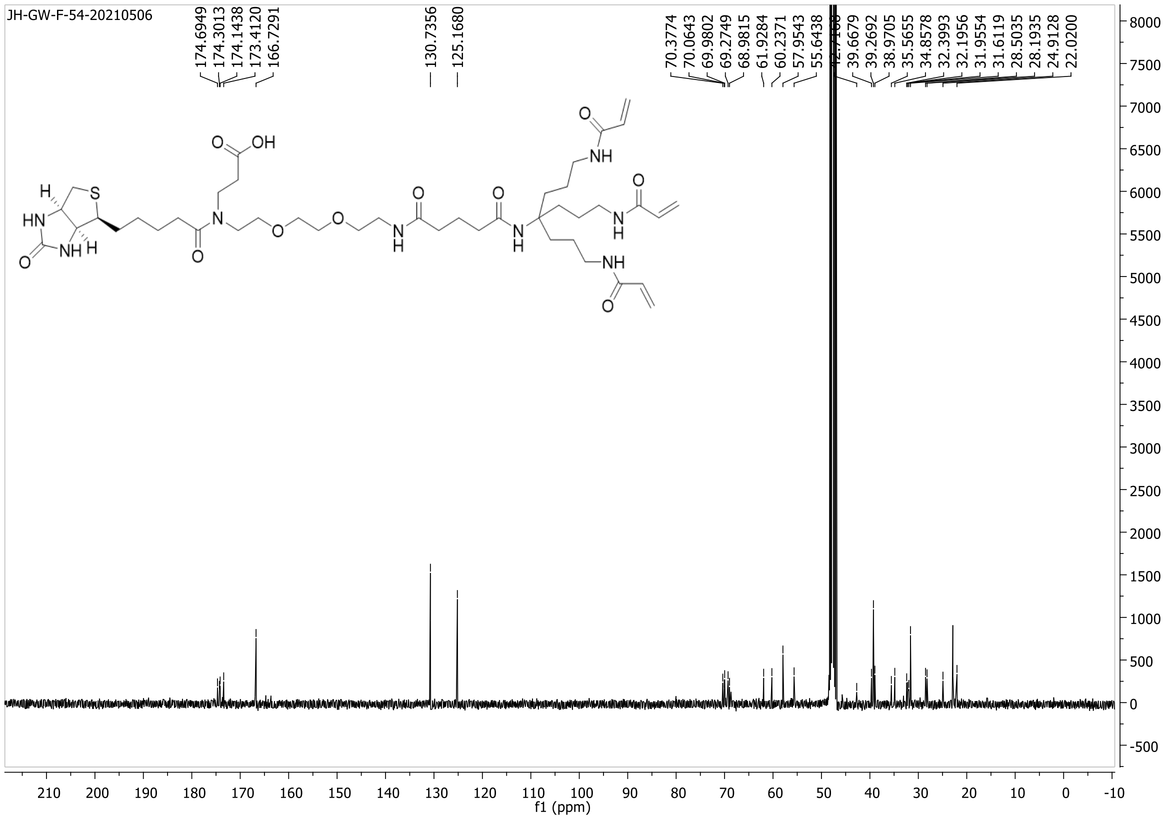
JH-GW-F-54-20210506
f1 (ppm)

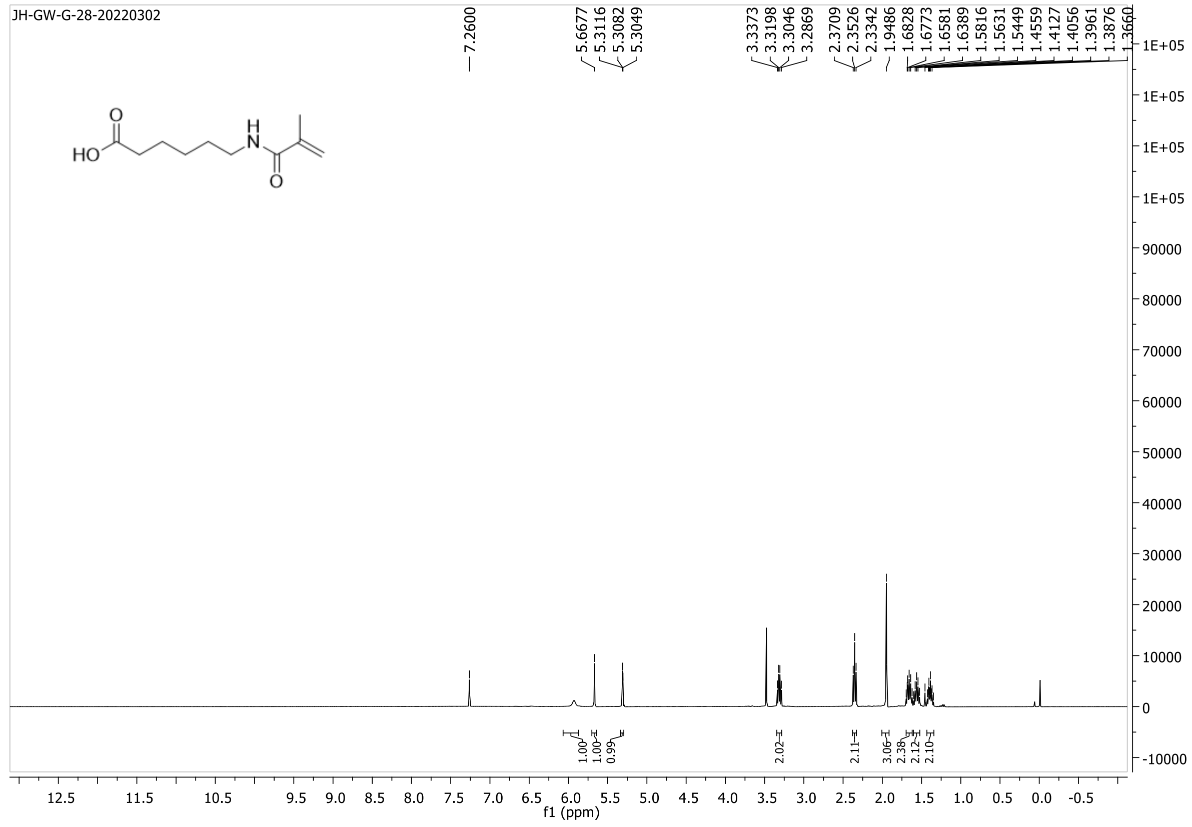
JH-GW-G-28-20220302
HO
O
H
N
O
f1 (ppm)

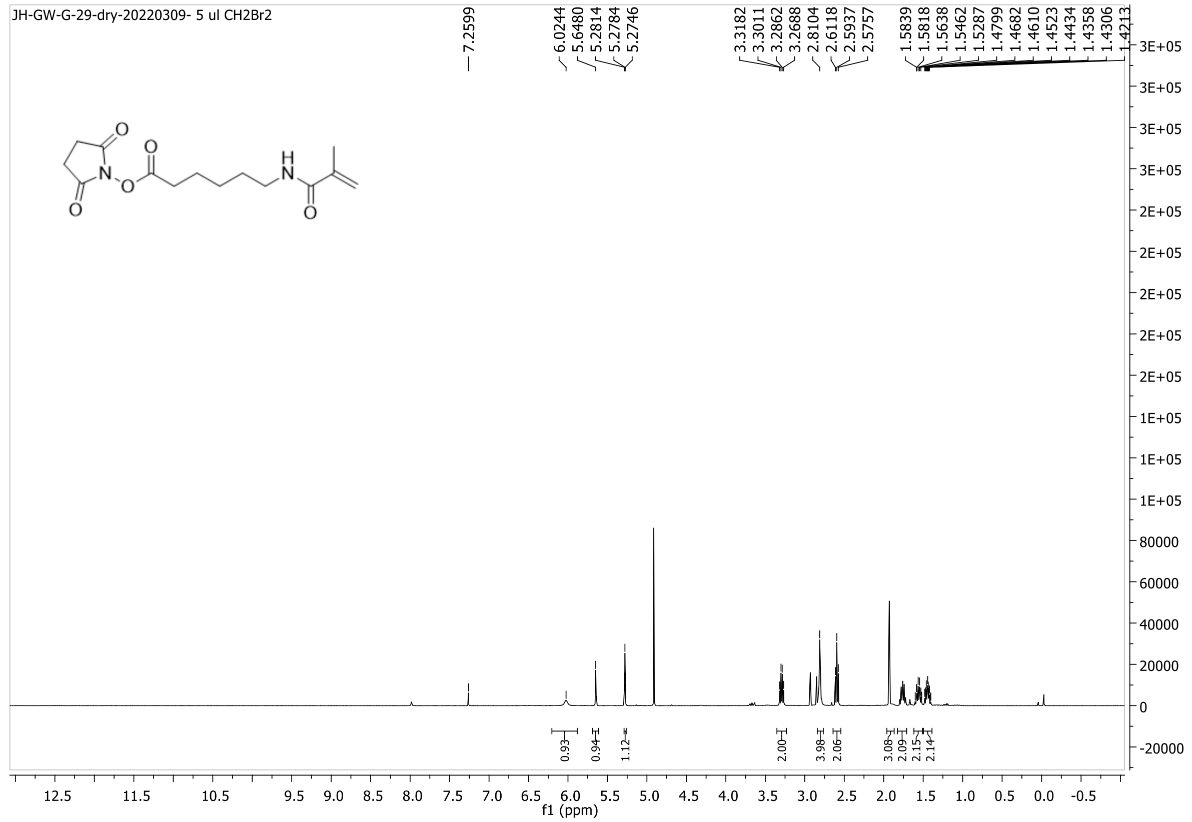
JH-GW-G-29-dry-20220309- 5 ul CH2Br2
O
N
O
O
O
H
N
O
f1 (ppm)

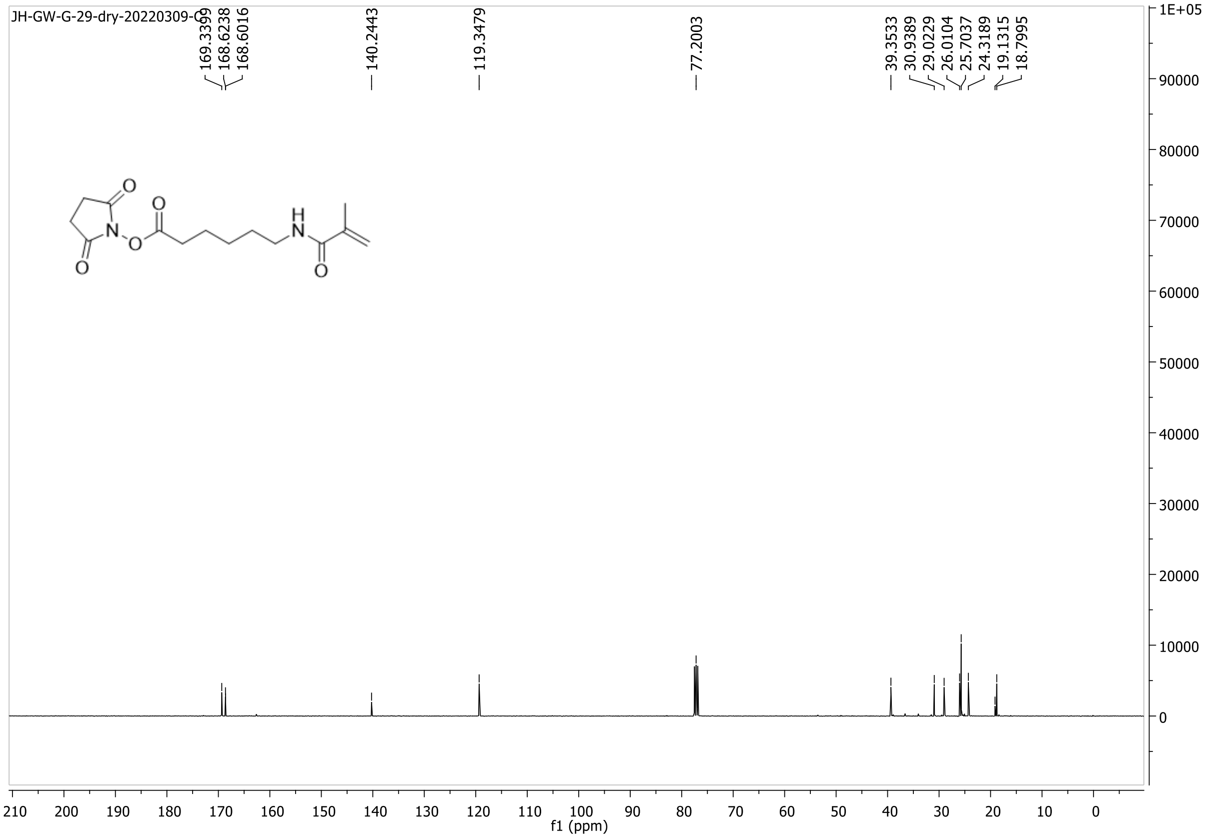
JH-GW-G-29-dry-20220309-C
169.3399
168.6238
168.6016
140.2443
119.3479
77.2003
39.3533
30.9389
29.0229
26.0104
25.7037
24.3189
19.1315
18.7995
f1 (ppm)

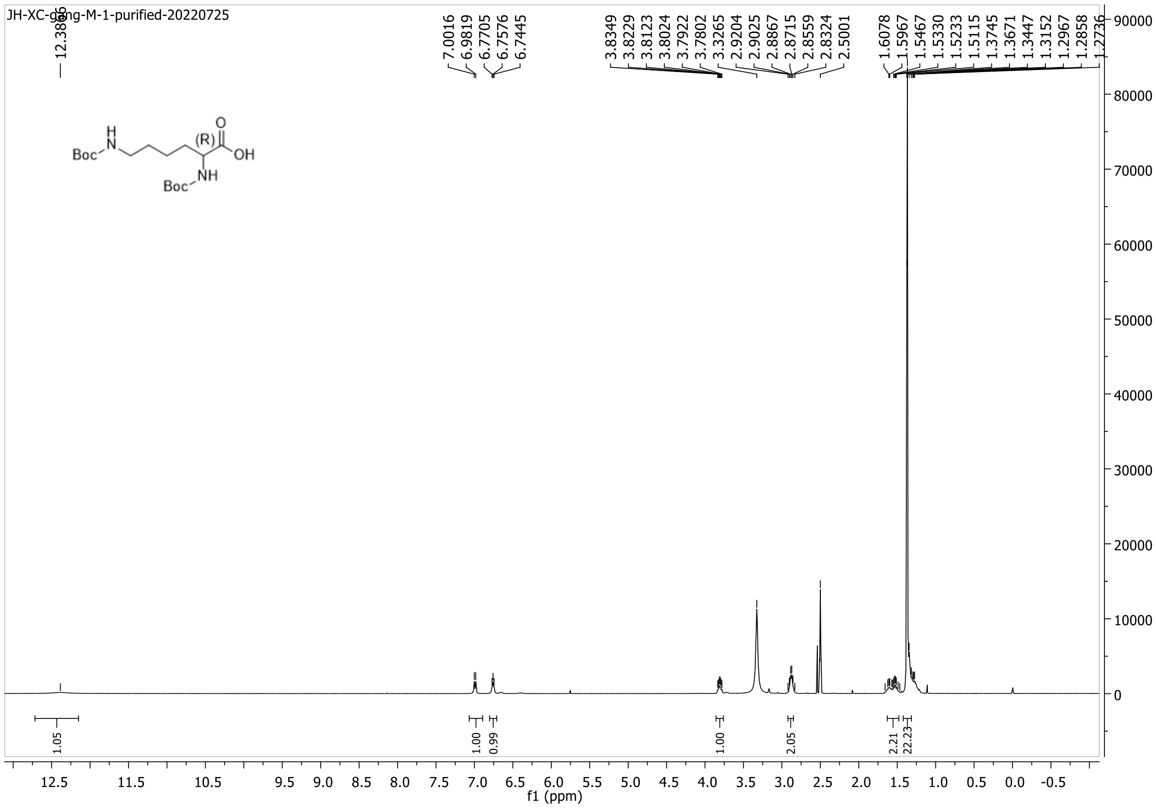
JH-XC-gang-M-1-purified-20220725
7.0016
6.9819
6.7705
6.7576
6.7445
3.8349
3.8229
3.8123
3.8024
3.7922
3.7802
3.3265
2.9204
2.9025
2.8867
2.8715
2.8559
2.8324
2.5001
1.6078
1.5967
1.5467
1.5330
1.5233
1.5115
1.3745
1.3671
1.3447
1.3152
1.2967
1.2858
1.2736
Boc
(R)
OH
NH
Boc
1.05
1.00
0.99
1.00
2.05
2.21
22.23
f1 (ppm)

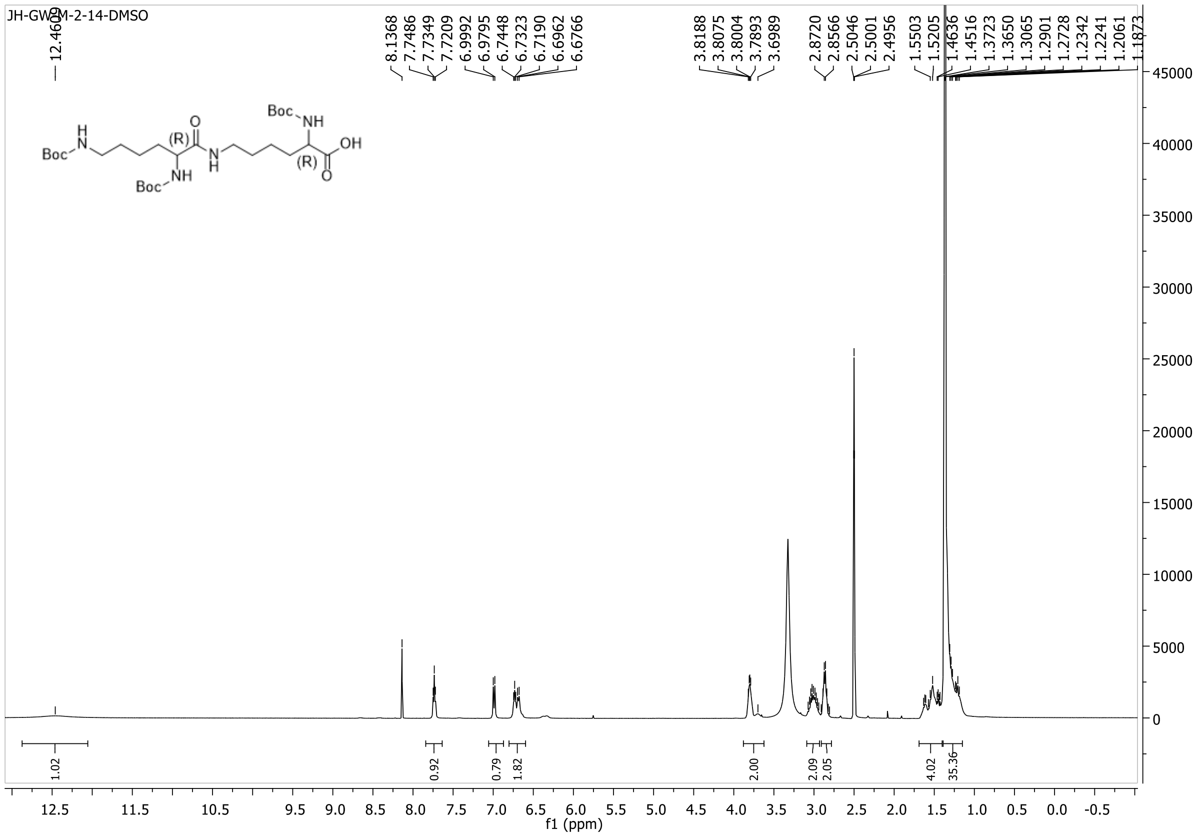

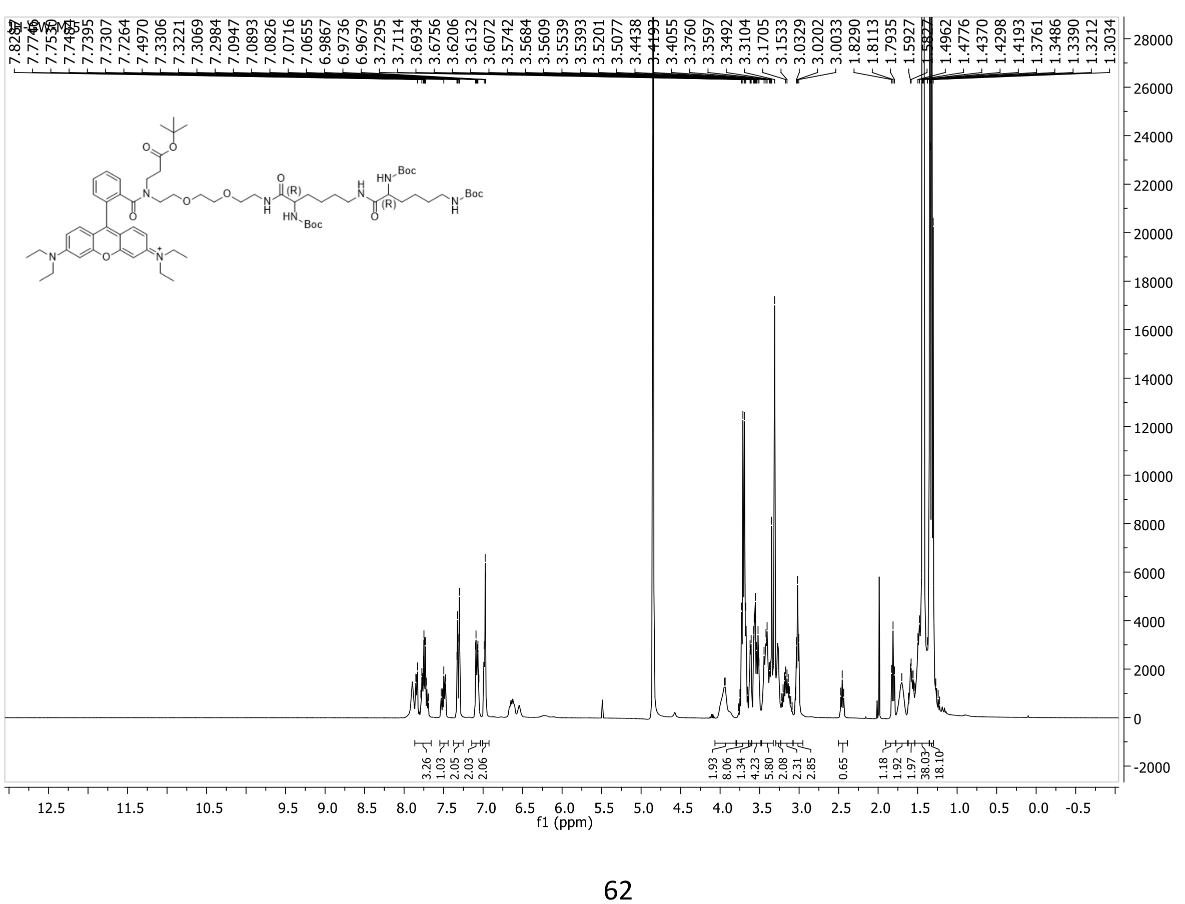

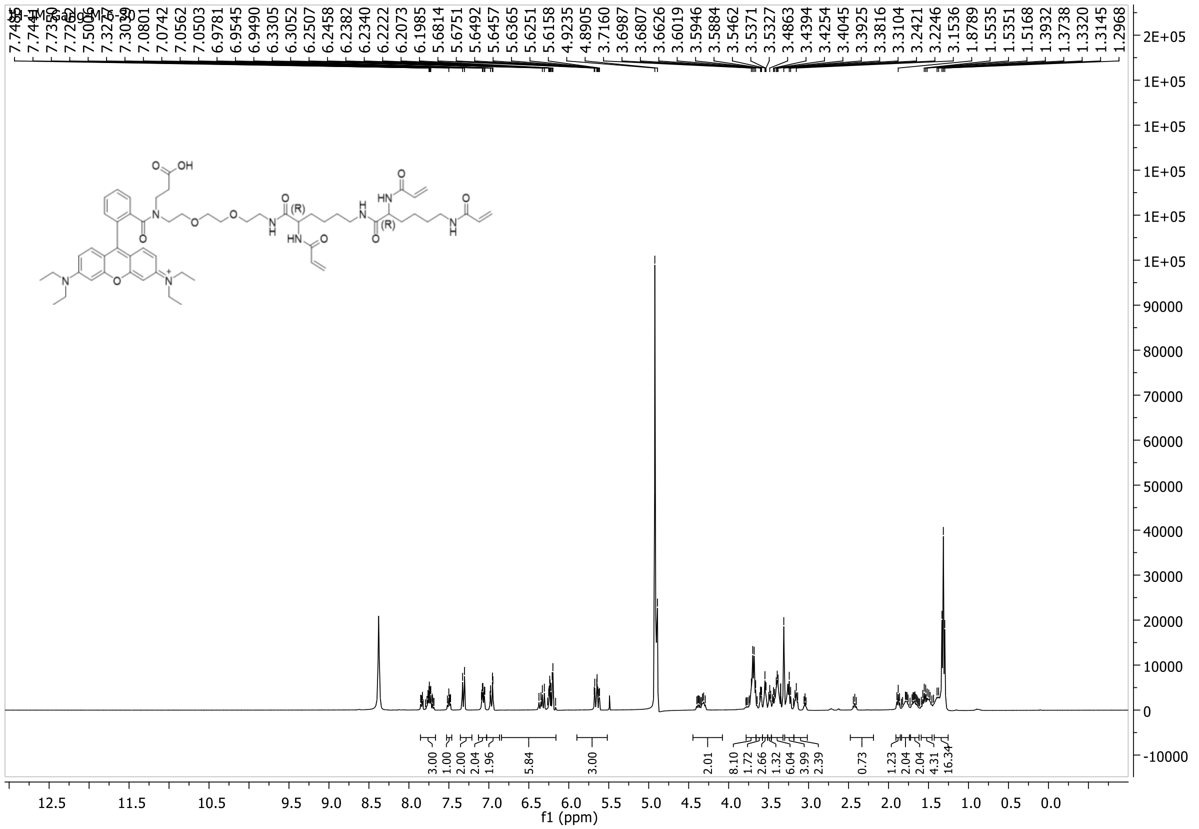
7.0801
7.0742
7.0562
7.0503
6.9781
6.9545
6.9490
6.3305
6.3052
6.2507
6.2458
6.2382
6.2340
6.2222
6.2073
6.1985
5.6814
5.6751
5.6492
5.6457
5.6365
5.6251
5.6158
4.9235
4.8905
3.7160
3.6987
3.6807
3.6626
3.6019
3.5946
3.5884
3.5462
3.5371
3.5327
3.4863
3.4394
3.4254
3.4045
3.3925
3.3816
3.3104
3.2421
3.2246
3.1536
1.8789
1.5535
1.5351
1.5168
1.3932
1.3738
1.3320
1.3145
1.2968
3.00
1.00
2.00
2.04
1.96
5.84
3.00
2.01
8.10
1.72
2.66
1.32
6.04
3.99
2.39
0.73
1.23
2.04
2.04
4.31
16.34
f1 (ppm)

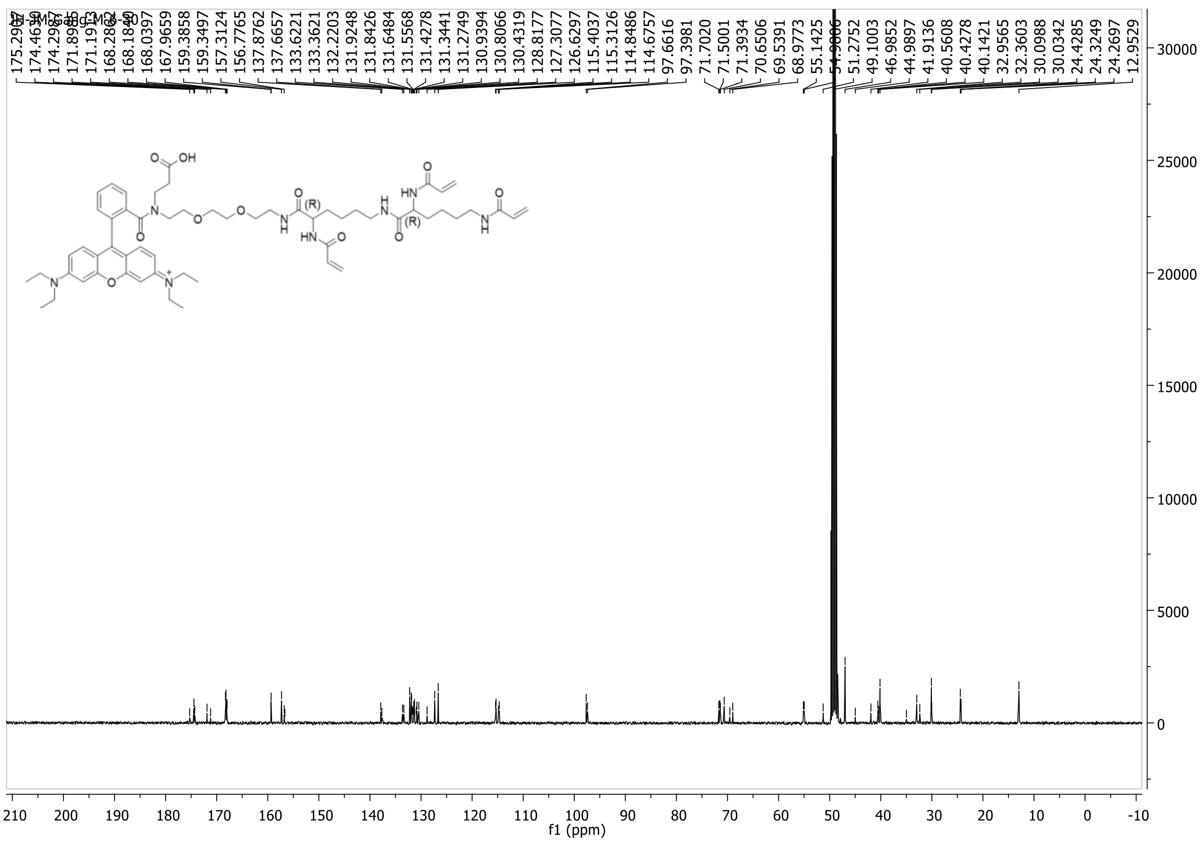
168.0397
167.9659
159.3858
159.3497
157.3124
156.7765
137.8762
137.6657
133.6221
133.3621
132.2203
131.9248
131.8426
131.6484
131.5568
131.4278
131.3441
131.2749
130.9394
130.8066
130.4319
128.8177
127.3077
126.6297
115.4037
115.3126
114.8486
114.6757
97.6616
97.3981
71.7020
71.5001
71.3934
70.6506
69.5391
68.9773
55.1425
51.2752
49.1003
46.9852
44.9897
41.9136
40.5608
40.4278
40.1421
32.9565
32.3603
30.0988
30.0342
24.4285
24.3249
24.2697
12.9529
f1 (ppm)

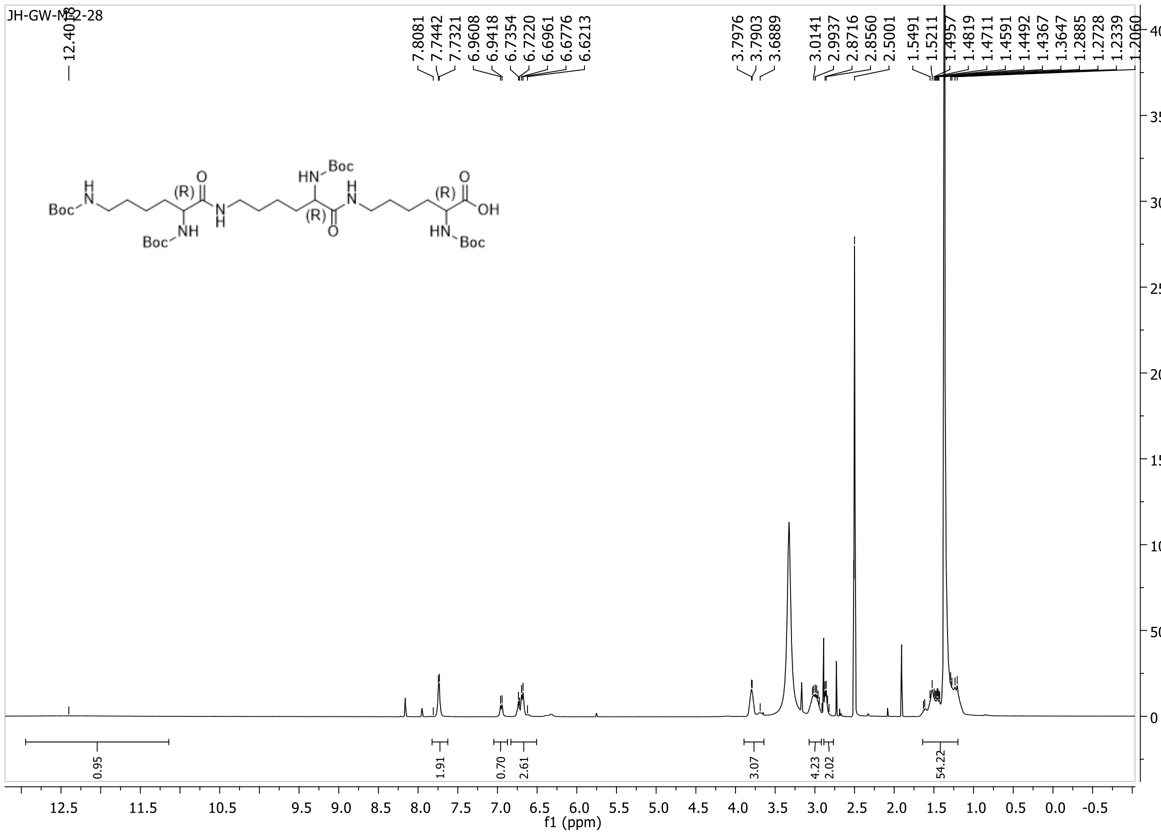

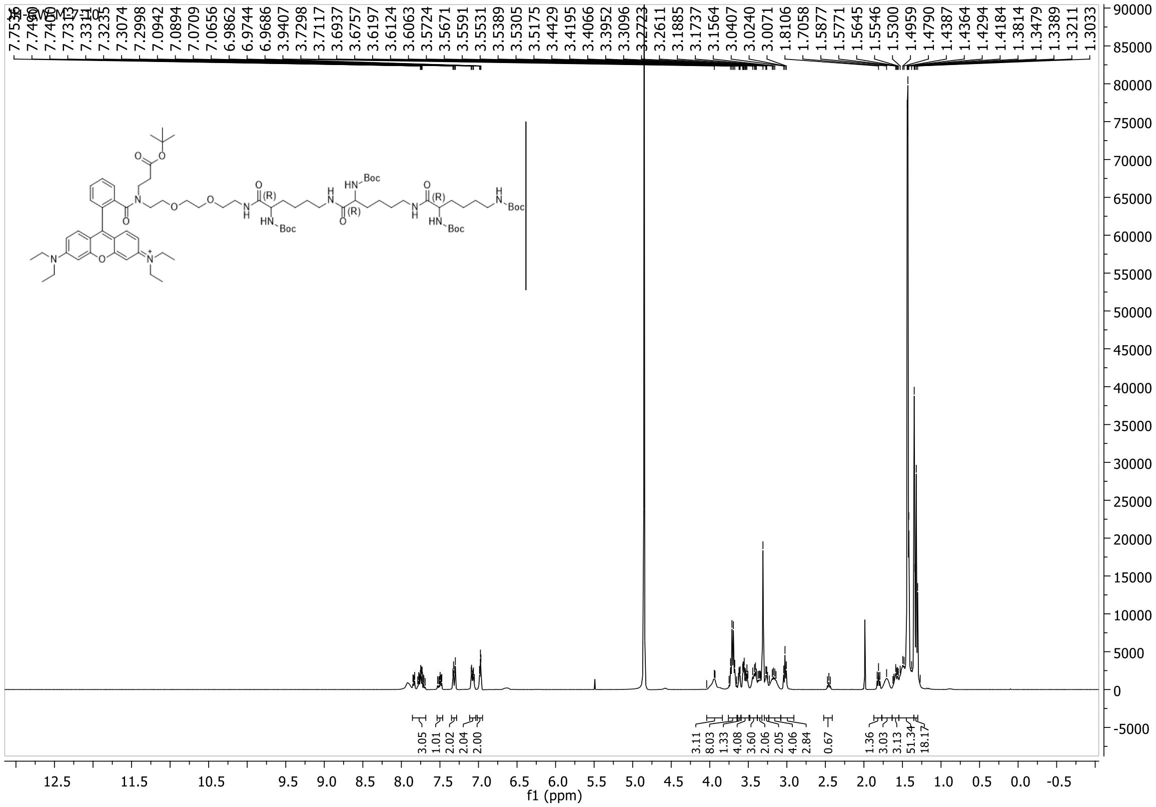

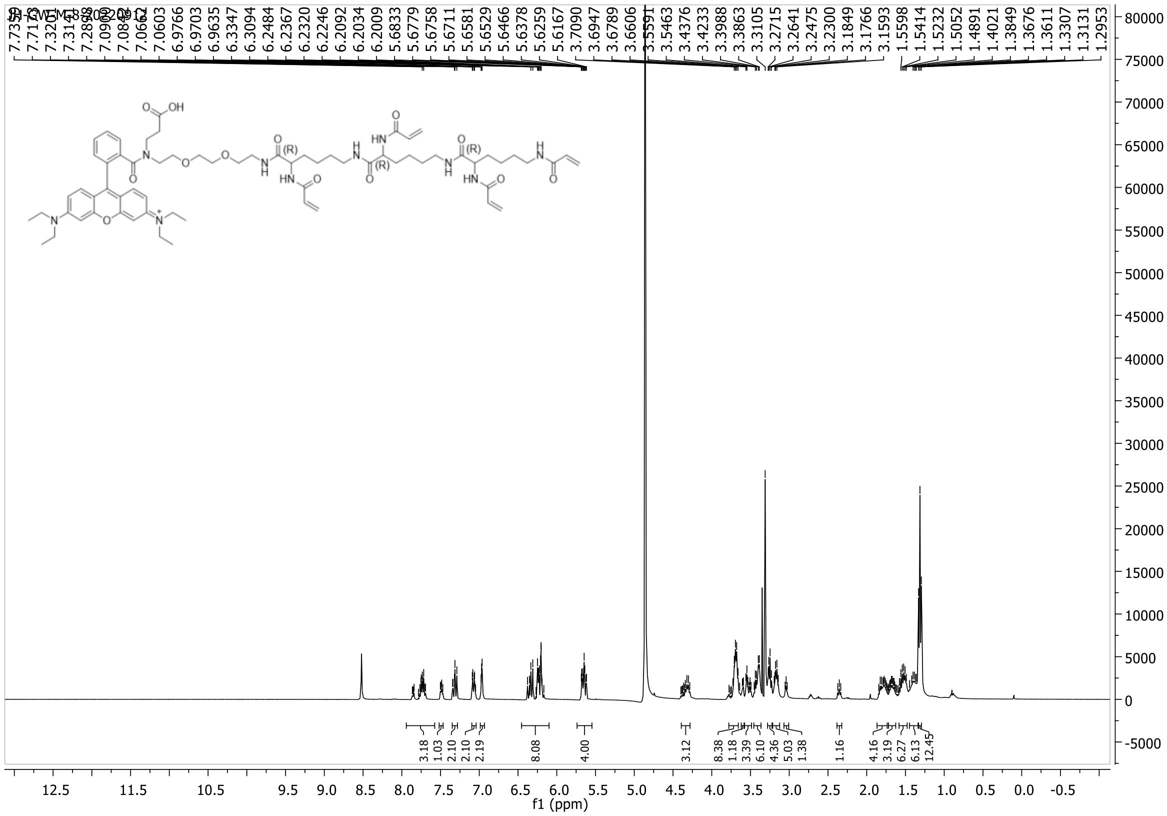
f1 (ppm)

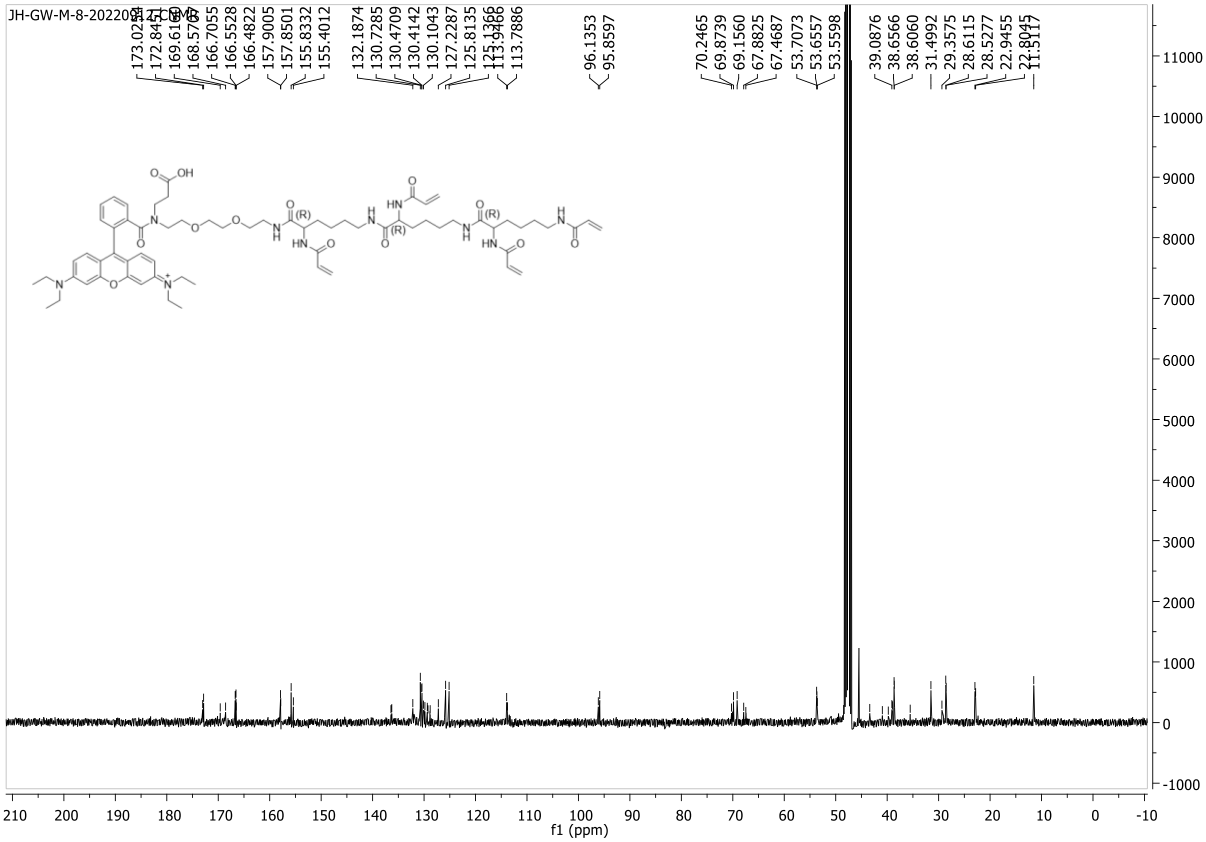
f1 (ppm)

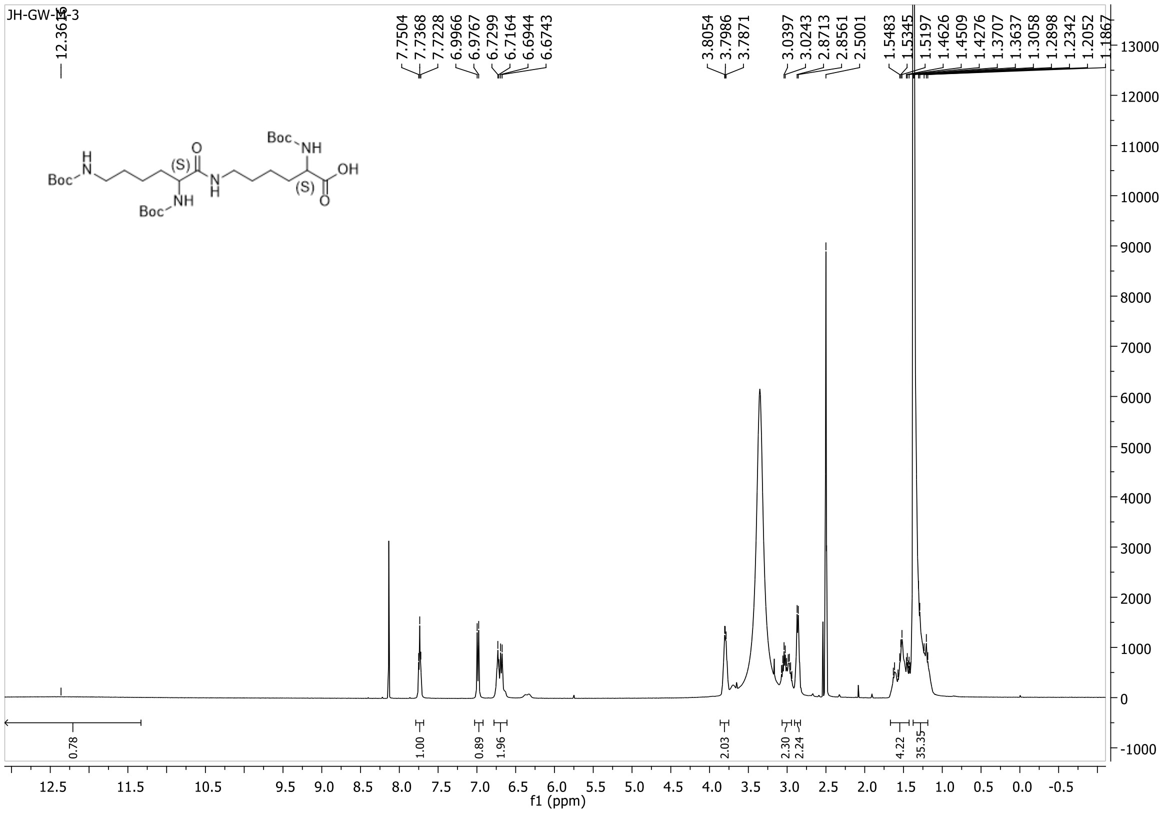
12.3616
7.7504
7.7368
7.7228
6.9966
6.9767
6.7299
6.7164
6.6944
6.6743
3.8054
3.7986
3.7871
3.0397
3.0243
2.8713
2.8561
2.5001
1.5483
1.5345
1.5197
1.4626
1.4509
1.4276
1.3707
1.3637
1.3058
1.2898
1.2342
1.2052
1.1867
0.78
1.00
0.89
1.96
2.03
2.30
2.24
4.22
35.35
f1 (ppm)
Boc
NH
(S)
OH

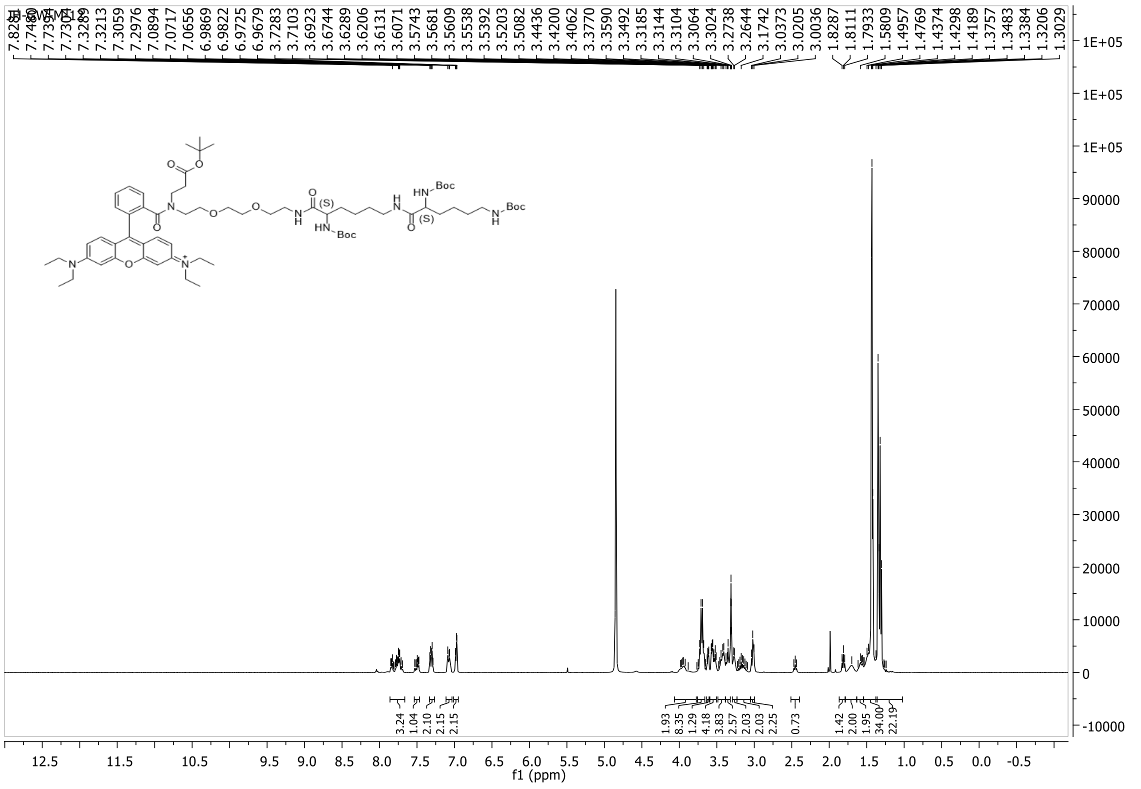
7.8287
7.7490
7.7353
7.7303
7.3289
7.3213
7.3059
7.2976
7.0894
7.0717
7.0656
6.9869
6.9822
6.9725
6.9657
3.7283
3.7103
3.6923
3.6744
3.6289
3.6206
3.6131
3.6071
3.5743
3.5681
3.5609
3.5538
3.5392
3.5203
3.5082
3.4436
3.4200
3.4062
3.3770
3.3590
3.3492
3.3185
3.3144
3.3104
3.3064
3.3024
3.2738
3.2644
3.1742
3.0373
3.0205
3.0036
1.8287
1.8111
1.7933
1.5809
1.4957
1.4769
1.4374
1.4298
1.4189
1.3757
1.3483
1.3384
1.3206
1.3029
3.24
1.04
2.10
2.15
2.15
1.93
8.35
1.29
4.18
3.83
2.57
2.03
2.03
2.25
0.73
1.42
2.00
1.95
34.00
22.19
f1 (ppm)
Boc
HN
(S)

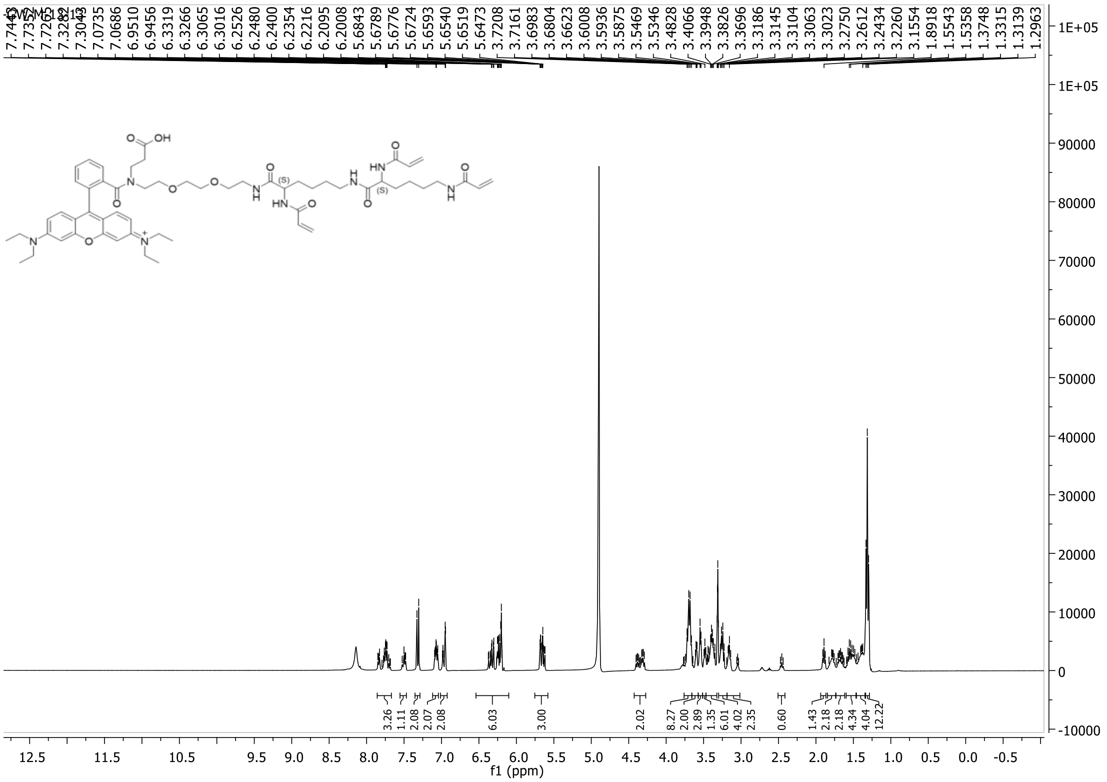
7.7443
7.7263
7.3282
7.3043
7.0735
7.0686
6.9510
6.9456
6.3319
6.3266
6.3065
6.3016
6.2526
6.2480
6.2400
6.2354
6.2216
6.2095
6.2008
5.6843
5.6789
5.6776
5.6724
5.6593
5.6540
5.6519
5.6473
3.7208
3.7161
3.6983
3.6804
3.6623
3.6008
3.5936
3.5875
3.5469
3.5346
3.4828
3.4066
3.3948
3.3826
3.3696
3.3186
3.3145
3.3104
3.3063
3.3023
3.2750
3.2612
3.2434
3.2260
3.1554
1.8918
1.5543
1.5358
1.3748
1.3315
1.3139
1.2963
OH
O
N
HN
NH
(S)
$N^+$
3.26
1.11
2.08
2.07
2.08
6.03
3.00
2.02
8.27
2.00
2.89
1.35
6.01
4.02
2.35
0.60
1.43
2.18
2.18
4.34
4.04
12.22
f1 (ppm)

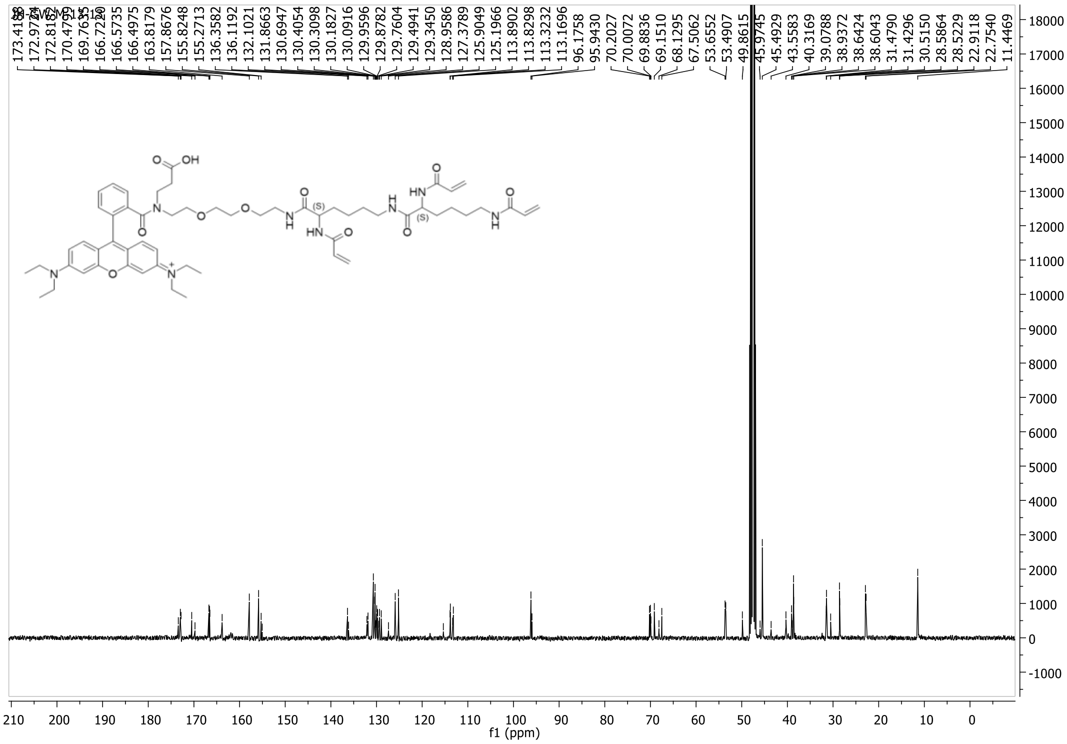
166.7220
166.5735
166.4975
163.8179
157.8676
155.8248
155.2713
136.3582
136.1192
132.1021
131.8663
130.6947
130.4054
130.3098
130.1827
130.0916
129.9596
129.8782
129.7604
129.4941
129.3450
128.9586
127.3789
125.9049
125.1966
113.8902
113.8298
113.3232
113.1696
96.1758
95.9430
70.2027
70.0072
69.8836
69.1510
68.1295
67.5062
53.6552
53.4907
49.8615
45.9745
45.4929
43.5583
40.3169
39.0788
38.9372
38.6424
38.6043
31.4790
31.4296
30.5150
28.5864
28.5229
22.9118
22.7540
11.4469
OH
O
N
HN
NH
(S)
$N^+$
f1 (ppm)
